# Chemical capture of diazo metabolites reveals biosynthetic hydrazone oxidation

**DOI:** 10.1101/2025.05.24.655970

**Authors:** Katarina Pfeifer, Devon Van Cura, Kelvin J. Y. Wu, Emily P. Balskus

**Author notes:** Authors contributed equally to the work (Katarina Pfeifer, Devon Van Cura).

## Abstract

Chemically reactive microbial natural products have enabled therapeutic development^1,2^ via their well-established bioactivities including anticancer,^3^ antibiotic,^4,5^ and antioxidant^6^ activities. However, discovery of reactive metabolites is particularly challenging because they may not tolerate traditional bioactivity-guided isolation workflows.^7^ Diazo-containing natural products are a subset of highly reactive microbial metabolites that display potent bioactivity^8–11^ and enable powerful (bio)synthetic transformations;^12,13^ however, instability of the diazo group to light,^14,15^ heat,^16,17^ mild acid,^18^ and mechanical shock^19^ has precluded their efficient discovery and application. Here, we develop a reactivity-based screening approach to capture diazo-containing metabolites and facilitate their discovery by mass spectrometry. This workflow revealed two novel diazo-containing natural products, 4-diazo-3-oxo-butanoic acid and diazoacetone, from the human lung pathogen *Nocardia ninae*. Biosynthetic investigations revealed a distinct enzymatic logic for diazo formation involving hydrazone oxidation catalyzed by the metalloenzyme Dob3, and biochemical characterization of Dob3 suggests promising future applications in biocatalysis. Overall, our work highlights the power of reactivity-guided strategies for identifying reactive metabolites and facilitating the discovery of unique enzymatic transformations.

## Introduction

Microorganisms produce various natural products containing the diazo functional group (C=N^+^=N^-^) including diazobenzofluorenes, diazobenzoquinones, ⍺-diazoketones, and ⍺-diazoesters (**Figure 1A**).^8^ The diazo group is highly reactive owing to the strong thermodynamic driving force of N_2_ release and can impart cytotoxicity to natural products through covalent and radical-induced damage of cellular targets. This has led to the exploration of diazo-containing metabolites as potential antibiotics and cancer chemotherapeutics.^8,20,21^ Diazo compounds are also used extensively as reagents and intermediates in synthetic chemistry as they enable powerful chemical transformations.^22^ Additionally, diazo compounds have emerged as important substrates for engineered heme-dependent enzymes that perform non-biological reactions in the context of biocatalysis^23^ and synthetic biology.^13,24^

**Figure 1:**
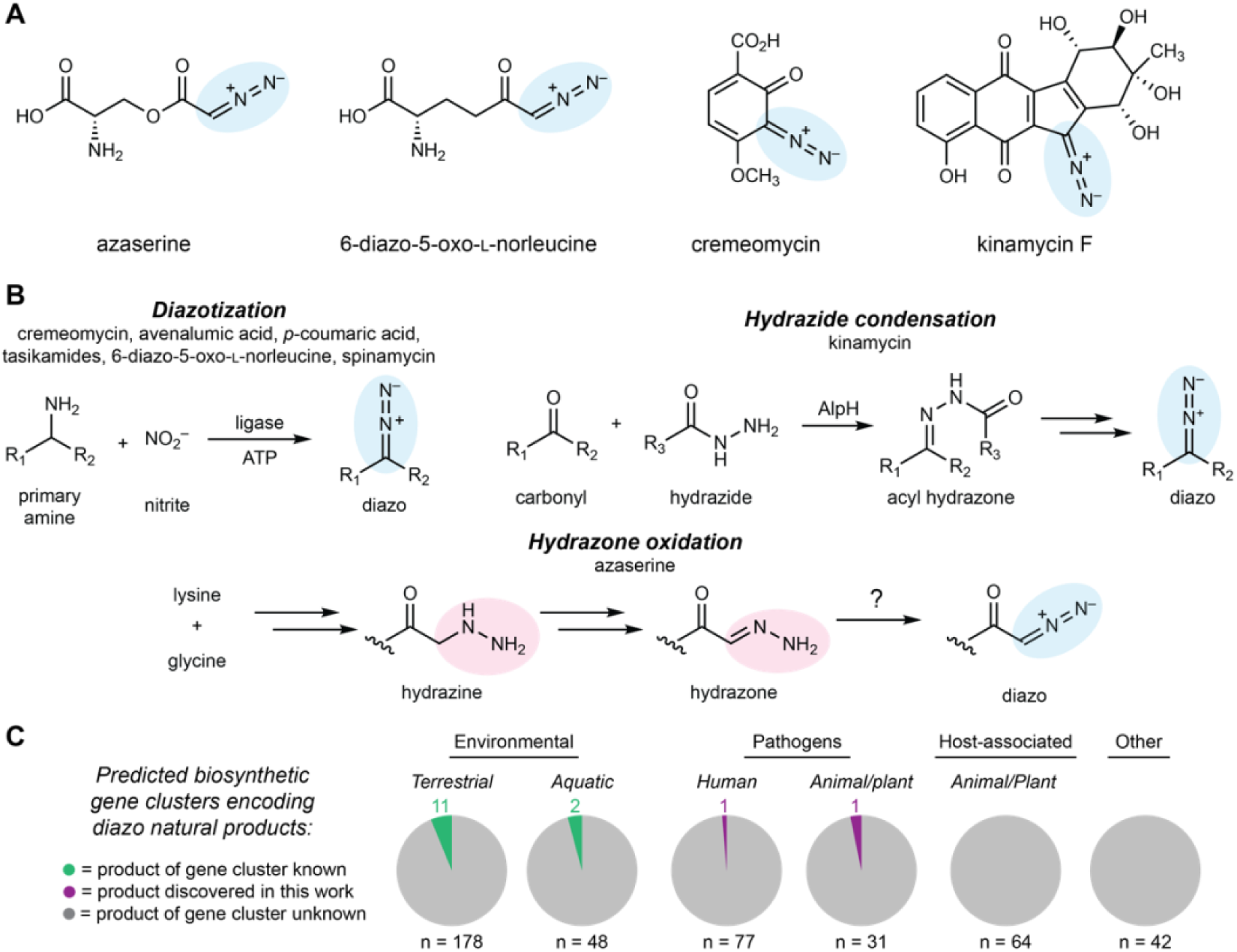
Discovery of new diazo-containing natural products may reveal chemical and biosynthetic diversity. **A)** Select diazo-containing natural products. **B)** Known diazo-forming enzymes perform diazotization or hydrazide condensation followed by non-enzymatic oxidation. Azaserine biosynthesis proceeds through enzymatic hydrazone formation, however, the hydrazone oxidoreductase has not yet been identified. **C)** Our genome mining efforts combined with previously reported efforts reveal 440 putative diazo-natural product-encoding biosynthetic gene clusters from diverse sources. As previous publications used enzymes from only four diazo-containing natural product biosynthetic pathways as queries, there are likely many additional diazo compound-encoding biosynthetic gene clusters that are not represented here.

The synthetic utility and biological activities of diazo-containing natural products have motivated efforts to understand their biosynthesis. To date, two distinct biosynthetic strategies to generate this functional group have been demonstrated biochemically. In the first, ATP-dependent ligases catalyze diazotization of primary amines using enzymatically produced nitrite (**Figure 1B**).^14,25–29^ The second strategy involves condensation of a hydrazide intermediate with a ketone followed by non-enzymatic oxidation to form the diazo group (**Supplementary** Figure 1)^30^ A third strategy, which was proposed in the azaserine biosynthetic pathway but has not yet been biochemically reconstituted, involves generation of a hydrazine intermediate, oxidation to an ⍺-hydrazonoacetyl intermediate, and conversion to an ⍺-diazoester by 2-electron *N*-oxidation.^31–33^ However, a dedicated hydrazone oxidoreductase has not been identified in this or any other diazo biosynthetic pathway.

Although <30 diazo-containing natural products have been reported, bioinformatic investigations suggest a much larger potential for microbial production of these metabolites. Compilation of previous genome mining efforts and our own searches has identified 440 biosynthetic gene clusters encoding putative diazo metabolites in diverse bacteria, including human, plant, and animal pathogens.^8,14,25,27,28,31,34^ However, only 13 of these gene clusters (3.2%) have known products (**Figure 1C**). We hypothesized that the discrepancy between the number of predicted and observed diazo-containing natural products is likely due to their chemical instability. Diazo compounds are sensitive to light,^14,15^ temperature,^16,17^ low pH,^18^ and mechanical shock,^19^ potentially complicating their discovery by traditional workflows. Challenges identifying and isolating these diazo metabolites therefore likely impede the discovery of natural products with intriguing biological activities and biosynthetic pathways.

In this study, we develop a reactivity-based natural product discovery workflow that uses a chemical trap to capture unstable diazo metabolites, enabling their identification and characterization by mass spectrometry. We use this approach to discover two related diazo-containing metabolites, 4-diazo-3-oxobutanoic acid (DOBA) and diazoacetone (DAC) from pathogenic *Nocardia* strains. Identification and in vitro reconstitution of the DOBA/DAC biosynthetic pathway reveals a ferritin-like diiron oxygenase/oxidase (FDO) enzyme, Dob3, that catalyzes the 2-electron *N*-oxidation of an ⍺-hydrazonoacetyl intermediate to produce the ⍺-diazoketone functional group. This discovery demonstrates a novel biosynthetic strategy for diazo formation, expands the known chemistry of the FDO enzyme family, and provides a starting place for biocatalytic diazo compound production.

## Results

### Reactivity-based screening workflow development

Traditional natural product discovery workflows such as bioactivity-guided fractionation, genome mining, heterologous expression, and genetic knockouts do not address the key challenge of diazo group instability. We envisioned employing an alternative discovery strategy by leveraging the reactive diazo functional group as a handle for chemical derivatization. Reactivity-based screening leverages chemical probes designed to react with a specific natural product functional group with minimal off-target reactivity, labeling metabolites of interest and facilitating their detection using LC–MS-based metabolomics.^7^ This approach has been employed to discover and/or isolate natural products with various functional groups, including polyyne- and isonitrile-containing natural products (**Supplementary** Figure 2).^35–40^ Further, chemical probes may be specially designed to enhance the stability and detection properties of labeled metabolites (e.g. UV absorption, ionization efficiency, etc.). While diazo compounds are known to participate in bioorthogonal 1,3-dipolar cycloadditions with alkynes,^41–44^ this reaction has not yet been exploited for diazo-containing natural product discovery.

To identify a chemical probe for our proposed discovery workflow, we examined previously developed diazo-alkyne cycloaddition reactions due to the prior applications of this reaction in complex biological matrices.^43^ We also expected the resulting pyrazole products^41^ to have enhanced ionization efficiency and increased lipophilicity relative to the corresponding diazo precursors, improving detection by reversed-phase LC–MS. We began by evaluating reactions between the model diazo-containing natural product azaserine and several alkynes. Of the alkynes tested, only strained cyclooctynes exhibited robust formation of the corresponding regioisomeric pyrazole products by LC–MS/MS (**Extended Data** Figure 1). Dibenzocyclooctyne C-6 acid (DBCO-acid) was selected for further optimization due to its superior yields.

The model reaction between DBCO-acid and azaserine was subsequently optimized (**Extended Data** Figure 2A-D). Aqueous solvents balanced high yields and facile sample handling, while a 16 h overnight incubation balanced product yield and experimental efficiency. Investigation of the reactivity of DBCO-acid toward other diazo compounds revealed good reactivity toward monosubstituted ⍺-diazocarbonyls (**Extended Data** Figure 3). During these analyses, we observed two peaks in the extracted ion chromatograms (EICs) of the DBCO-acid adducts, corresponding to the two product regioisomers produced during the 1,3-dipolar cycloaddition reaction. Products were not observed with disubstituted diazo-containing compounds, including the diazoquinone natural product cremeomycin, despite previous reports of successful reactions between cyclooctynes and disubstituted diazo-containing compounds.^41,45^

To assess the ability of DBCO-acid to label diazo-containing natural products in complex biological matrices, we performed the reaction using spent culture medium from the azaserine producing bacterium *Glycomyces harbinensis*.^46^ Targeted LC–MS analysis of spent culture medium revealed a mass feature with the same retention time, exact mass, and MS/MS fragmentation as an authentic standard of azaserine-DBCO, validating the applicability of the workflow for detecting diazo-containing natural products in complex metabolite mixtures (**Extended Data** Figure 4). We next used *G. harbinensis* spent medium to develop an untargeted LC–MS-based comparative metabolomics workflow (**Extended Data** Figure 5). Comparative metabolomics of DBCO-acid treated spent medium at reaction initiation (t_0_) versus a 16 h incubation (t_16_) revealed 7 metabolites that were significantly increased (p < 0.05, fold change ≥ 2), including the expected azaserine-DBCO regioisomers. These data validated our reactivity-based screening approach for diazo-containing natural product discovery.

### Discovery of diazo-containing metabolites

We then sought to apply our workflow to discover previously unidentified diazo-containing microbial natural products (**Figure 2A**). To prioritize organisms for screening, we mined bacterial genomes in the National Center for Biotechnology Information database for biosynthetic gene clusters encoding putative diazo-containing natural products. We chose to target diazo-containing metabolites likely to be produced by a hydrazone *N*-oxidation biosynthetic strategy to facilitate the study of this enigmatic transformation proposed to be important for azaserine biosynthesis.^31–33^ We identified biosynthetic gene clusters of interest by searching for homologs of hydrazone biosynthetic enzymes^31–33,47–50^ (**Supplementary** Figure 3) encoded alongside at least one additional predicted oxidoreductase. This analysis identified 129 biosynthetic gene clusters spanning a diverse range of Actinobacteria and Proteobacteria, including many *Streptomyces* species. Analysis of the gene clusters using prettyClusters^51^ identified five conserved biosynthetic gene cluster architectures in addition to the known azaserine, triacsin, and s56-p1 gene clusters (**Extended Data** Figure 6). These gene clusters encode many additional enzymes beyond the hydrazone biosynthetic enzymes, including potential diazo-forming oxidoreductases, indicating the potential for production of an array of diverse natural product structures.

**Figure 2:**
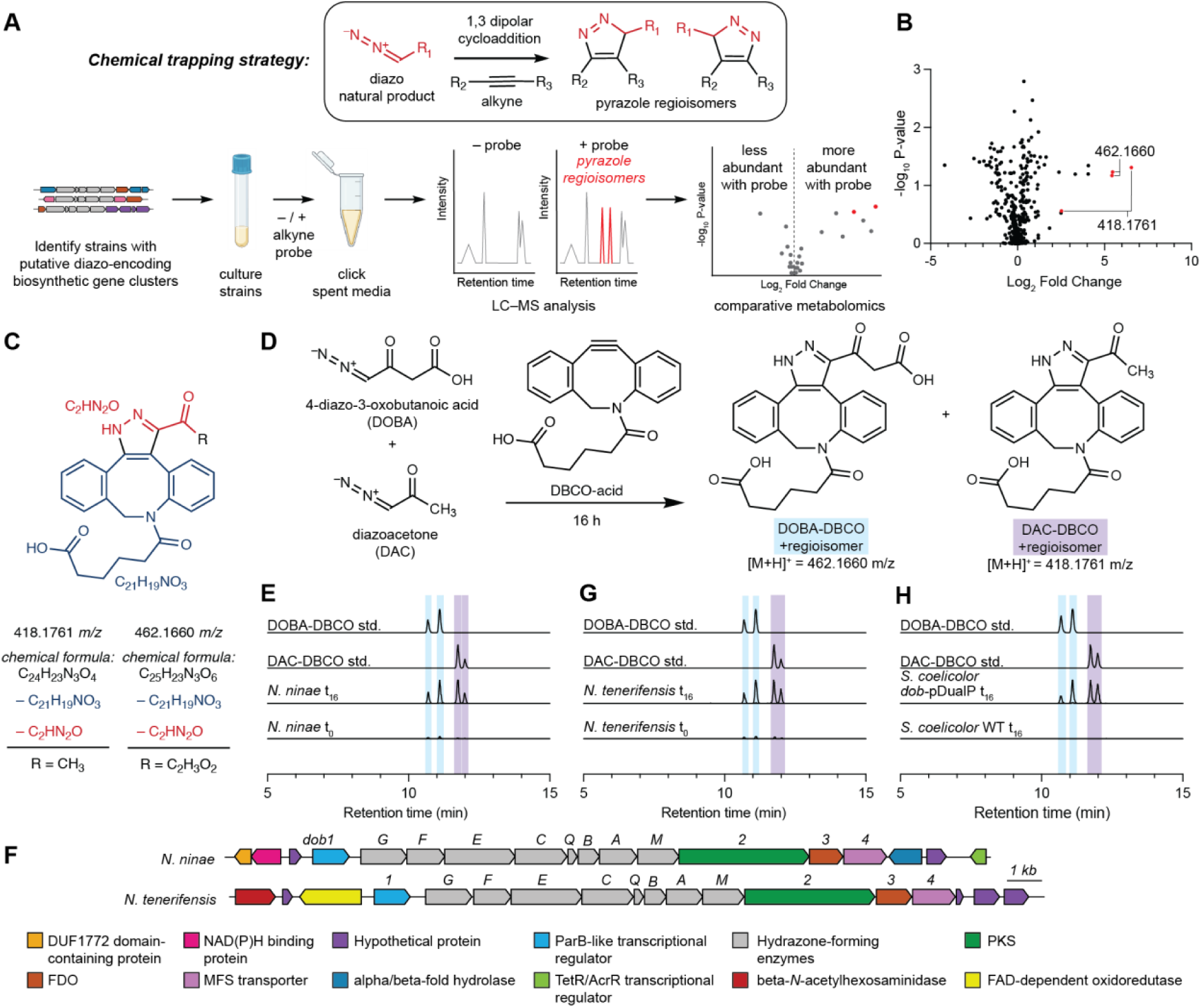
A reactivity-guided comparative metabolomics workflow uncovers DOBA and DAC from the human pathogen *Nocardia ninae*. **A)** Reactivity-based screening workflow. **B)** Volcano plot for comparative metabolomics of *N. ninae* spent medium. **C)** Analysis of predicted chemical formulas suggests DOBA and DAC as novel diazo-containing natural products. **D)** Reaction of DAC and DOBA with DBCO-acid yields DAC-DBCO and DOBA-DBCO. **E)** Extracted ion chromatograms of *m/z* = 418.1761 ± 5 ppm and *m/z* = 462.1660 ± 5 ppm in DBCO-treated *N. ninae* spent media at t = 0 h and t = 16 h compared to DOBA-DBCO and DAC-DBCO synthetic standards. EICs for 462.1660 and 418.1761 are superimposed. t_0_ and t_16_ data was normalized to the t_16_ maximum separately for DOBA-DBCO and DAC-DBCO. **F)** The *dob* biosynthetic gene cluster in *N. ninae* and *N. tenerifensis*. PKS = polyketide synthase, NRPS = non-ribosomal peptide synthetase, FDO = ferritin-like diiron oxidase or oxygenase. Extracted ion chromatograms of *m/z* = 418.1761 ± 5 ppm and *m/z* = 462.1660 ± 5 ppm in **G)** DBCO-treated *Nocardia tenerifensis* spent media and **H)** *Streptomyces coelicolor dob*-pDualP spent media at t = 0 h and t = 16 h compared to DOBA-DBCO and DAC-DBCO synthetic standards. Experiments were performed in biological triplicate and representative results are shown. EICs for 462.1660 and 418.1761 are superimposed. t_0_ and t_16_ data was normalized to the t_16_ maximum separately for DOBA-DBCO and DAC-DBCO.

We selected 10 commercially available strains identified in this analysis, spanning 3 of the 5 identified gene cluster architectures, for cultivation and analysis using our optimized comparative metabolomics workflow. Cultures were grown in up to 6 different growth media (molasses production media, ISP4, GYM, R2B, and MSF) for 5 to 7 days. Spent culture media were incubated with DBCO-acid, and LC–MS-based comparative metabolomics was performed in positive ion mode. Potential hits were observed from *Streptomyces yunnanensis* and *Nocardia ninae* (**Figure 2B**). Only two pairs of mass features obtained from spent media of *N. ninae* were present in t_16_ but not t_0_ samples and featured the doubled EIC peaks characteristic of DBCO-acid adduct regioisomers (418.1761 *m/z* and 462.1660 *m/z*) (**Supplementary Data**). Furthermore, their neutral molecular formulas, C_24_H_23_N_3_O_4_ (*m/z* = 418.1761) and C_25_H_23_N_3_O_6_ (*m/z* = 462.1660), contained sufficient nitrogen atoms to support the presence of a pyrazole functional group. The molecular formula of each putative diazo-containing natural product was determined by subtracting the molecular formula of DBCO-acid (C_21_H_19_NO_3_) to give C_4_H_4_N_2_O_3_ and C_3_H_4_N_2_O (**Figure 2C**). Based on the reactivity profile of DBCO-acid and the hydrazone-forming biosynthetic enzymes present in this strain, we hypothesized that both natural products contained a hydrazone-derived ⍺-diazoacetyl group. Subtracting the formula of the diazoacetyl group (C_2_HN_2_O) from the molecular formulas of both putative diazo-containing natural products yielded C_2_H_3_O_2_ and CH_3_ as the remaining atoms in each molecule, suggesting the original diazo natural products were 4-diazo-3-oxobutanoic acid (DOBA) and diazoacetone (DAC), respectively (**Figure 2C,D**).

To verify the structures of these natural products, we attempted to synthesize authentic standards of DOBA-DBCO and DAC-DBCO. While DAC-DBCO was readily accessible, DOBA-DBCO rapidly decarboxylated, precluding its isolation and purification. Accordingly, we synthesized DOBA-DBCO methyl ester (OMe-DOBA-DBCO) which was stable and could be readily deprotected enzymatically *in situ* to provide DOBA-DBCO under mild conditions. LC–MS analysis of DAC-DBCO and deprotected OMe-DOBA-DBCO standards revealed exact masses, retention times, and MS/MS fragmentation patterns identical those of the 462.1660 and 418.1761 masses observed from *N. ninae*, confirming the derivatized structures as DOBA-DBCO and DAC-DBCO, respectively (**Figure 2E**, **Supplementary** Figure 4). Notably, we were unable to observe underivatized DOBA and DAC in spent culture medium using LC–MS. This suggests these metabolites would not have been readily identified using standard analytical techniques, demonstrating the utility of our reactivity-guided discovery workflow. DOBA and DAC have not previously been identified in living systems, demonstrating the ability of this workflow to identify novel natural products.

### Gene cluster discovery and validation

We next set out to identify the biosynthetic gene cluster responsible for DOBA/DAC production to facilitate studies of diazo biosynthesis. Our genome mining analysis highlighted two potential diazo-forming biosynthetic gene clusters in *N. ninae*, “*dob”* and “*nin”* (**Supplementary** Figure 5), which both encode hydrazone-forming enzymatic machinery and an additional oxidoreductase. To determine the gene cluster responsible for DOBA/DAC production, we formulated a biosynthetic hypothesis for these metabolites. Related hydrazone-forming pathways use a conserved set of biosynthetic enzymes to generate a carrier protein-bound ⍺-hydrazonoacetyl thioester that is transferred to a nucleophilic intermediate.^31–33,50^ The structures of DOBA and DAC require the formation of a C–C bond between the ⍺-hydrazonoacetyl thioester intermediate and a carbon-based nucleophile. Polyketide synthases (PKSs) catalyze C–C bond-forming Claisen-type condensation reactions. Based on the structure of DOBA, we hypothesized that a PKS might catalyze C–C bond formation between malonyl-CoA and the ⍺-hydrazonoacetyl thioester intermediate to yield 4-hydrazono-3-oxobutanoic acid (HOBA) which could feasibly undergo a subsequent 2-electron *N*-oxidation to produce the diazo functional group of DOBA. We hypothesized that DAC could be produced by spontaneous non-enzymatic decarboxylation of DOBA.

Only the *dob* gene cluster encoded a predicted PKS consistent with a role in DOBA/DAC biosynthesis (**Figure 2F**). BLAST searches and genomic neighborhood analysis using EFI-GNT indicated that 14 additional *Nocardia* strains contain the *dob* gene cluster (**Supplementary Table 5**).^52,53^ To test the link between this gene cluster and DOBA/DAC biosynthesis, a selection of these strains was cultivated, spent media was derivatized with DBCO-acid, and targeted LC– MS/MS analysis was performed to assess DOBA-DBCO and DAC-DBCO production. One of the strains, *Nocardia tenerifensis,* produced mass features with exact masses, retention times, and MS/MS fragmentation patterns identical to those of DOBA-DBCO and DAC-DBCO, strengthening the relationship between the *dob* gene cluster and DOBA/DAC production (**Figure 2G**, **Supplementary** Figure 6).

To confirm the role of the *dob* gene cluster in DOBA/DAC biosynthesis, we expressed the gene cluster in a heterogenous host. The *dob* biosynthetic gene cluster was cloned into a dual-inducible vector using modified Gibson Assembly to produce *dob*-pDualP (**Supplementary** Figure 7). The *dob*-pDualP vector was transferred into the heterologous host *S. coelicolor* M1152 by conjugation with *E. coli* pUZ8002/ET12567. *S. coelicolor dob*-pDualP was cultured in a variety of growth media supplemented with ε-caprolactam and oxytetracycline inducers. Spent culture medium was derivatized with DBCO-acid and assayed for DOBA-DBCO and DAC-DBCO production. Only the heterologous host harboring *dob*-pDualP, and not the wild-type strain, produced DOBA-DBCO and DAC-DBCO, definitively linking the *dob* gene cluster to DOBA and DAC biosynthesis (**Figure 2H, Supplementary** Figure 8).

### Discovery of a diazo-forming metalloenzyme

With the role of the *dob* gene cluster in DOBA/DAC production validated, we next sought to assign biosynthetic roles to each of the encoded enzymes. DobG, E, F, B, C, Q, M, and A are homologous to hydrazone-forming enzymes which produce a carrier protein-bound ⍺-hydrazonoacetyl thioester, hydrazonoacetic acid (HYAA), from L-lysine and glycine (**Supplementary** Figure 3).^31–33,50^ Analysis of the proteins encoded by the remaining *dob* genes suggested the Type I PKS Dob2 as a likely candidate to catalyze C–C bond formation, and the ferritin-like diiron oxidase/oxygenase (FDO) Dob3 as a potential diazo-forming oxidoreductase. The remaining biosynthetic genes are predicted to encode a transcriptional regulator (Dob1) and a Major Facilitator Superfamily transporter (Dob4).

To investigate DOBA/DAC biosynthesis, we first sought to heterologously express (**Supplementary** Figure 9) and reconstitute the activity of the putative hydrazone biosynthetic enzymes previously described in the triacsin and azaserine pathways.^31–33,50^ As expected, the *N*-oxygenase DobG catalyzed the NAD(P)H- and FAD-dependent oxidation of L-lysine to *N*-6-hydroxylysine (**Supplementary** Figure 10). Attempts to reconstitute activity of the MetRS/cupin fusion enzyme DobE in vitro were unsuccessful; however, biotransformations using *E. coli* expressing DobE incubated with *N*-6-hydroxylysine, glycine, and ATP yielded *N-*6-(carboxymethylamino)lysine (**Supplementary** Figure 11). Hydrazinoacetic acid (HAA) was succinylated by the GCN5-*N*-acetyltransferase DobB (**Supplementary** Figure 12).

We reconstituted the activity of the remaining hydrazone biosynthetic enzymes in a single cascade reaction. HAA was incubated with succinyl-CoA, DobB, ATP, DobC, *holo*-DobQ, FAD, DobM, and DobA. As expected, we observed production of succinyl-HAA-DobQ and HYAA-DobQ intermediates (**Supplementary** Figure 13). In contrast to previous reports, we also observed accumulation of a mass consistent with succinyl-HYAA-DobQ in reactions lacking the C45 peptidase DobA. As such, we hypothesize that during DOBA biosynthesis, HAA is first succinylated by DobB, then succinyl-HAA is loaded onto the carrier protein DobQ by the adenylase DobC, succinyl-HAA-DobQ is oxidized to succinyl-HYAA-DobQ by the FAD-dependent oxidoreductase DobM, and finally succinyl-HYAA-DobQ is hydrolyzed by DobA to release HYAA-DobQ. Our observation of hydrolysis after oxidation differs from previous reports which suggested that hydrolysis either precedes^33^ or is concurrent with^31,50^ oxidation.

After confirming that the hydrazone-forming enzymes produce the expected ⍺-hydrazonoacetyl intermediate HYAA-DobQ, we sought to investigate the unique enzymatic transformations leading to DOBA/DAC production. Type I PKS Dob2 is predicted to contain thioesterase, ketosynthase, acyltransferase, and acyl carrier protein domains (TE-KS-AT-ACP). The Dob2 AT domain was predicted to load methylmalonyl-CoA by antiSMASH,^54^ but based on the structures of DAC and DOBA, we hypothesized that this domain could use malonyl-CoA. Based on canonical PKS logic,^55^ we hypothesized that HYAA may be translocated from DobQ to a conserved Cys residue in the KS domain of Dob2 prior to KS domain-catalyzed C–C bond-forming decarboxylative Claisen condensation with a malonyl extender unit to form 4-hydrazono-3-oxobutanoic acid-Dob2 (HOBA-Dob2). Finally, the TE domain of Dob2 might catalyze translocation to a conserved Ser residue prior to hydrolytic release of the mature PKS product 4-hydrazono-3-oxobutanoic acid (HOBA; **Supplementary** Figure 14).

The putative diazo-forming enzyme Dob3 was the sole unassigned oxidoreductase encoded in the *dob* gene cluster and therefore the most likely candidate for hydrazone oxidation. A structure homology search of the Dali Webserver^56^ using an AlphaFold^57^ predicted structure of Dob3 revealed strong structural homology to the characterized ferritin-like diiron oxygenases (FDOs) AurF (Z=40.1, 31% sequence identity) and CmlI (Z=40.9, 47% sequence identity), which catalyze 6-electron *N*-oxidation of aryl amines to aryl nitro groups during the biosynthesis of aureothin and chloramphenicol, respectively (**Supplementary** Figure 15).^58,59^ Based on this analysis, we hypothesized that Dob3 could catalyze *N*-oxidation of a hydrazone to a diazo group during DOBA biosynthesis. However, the order of reactions catalyzed by Dob2 and Dob3 in the pathway remained unclear.

We next set out to characterize these enzymes by first attempting to reconstitute the activity of Dob2 in vitro. However, incubation of *holo*-Dob2 and malonyl-CoA with enzymatically produced HYAA-DobQ failed to produce a mass feature consistent with the predicted hydrazone product 4-hydrazono-3-oxobutanoic acid (HOBA). We suspected this could be due to a combination of low yields and poor ionization efficiency rather than incorrect assignment of the biochemical transformation. Accordingly, we attempted to use HOBA as a substrate to reconstitute the activity of the putative diazo-forming enzyme Dob3. As the susceptibility of HOBA to decarboxylation precluded its direct synthesis, we synthesized HOBA *O*-methyl ester (OMe-HOBA) and used enzymatic deprotection to generate HOBA in situ. Incubation of OMe-HOBA with porcine liver esterase (PLE), Dob3, phenazine methosulfate (PMS), NADPH, and DBCO-acid revealed production of DOBA-DBCO and DAC-DBCO, demonstrating the ability of Dob3 to catalyze hydrazone *N*-oxidation to form the diazo group of DOBA (**Figure 3A**). Given the facile detection of DOBA-DBCO and DAC-DBCO by LC–MS, we then revisited the activity of Dob2 in a coupled enzyme assay with Dob3 followed by DBCO-acid derivatization. Enzymatically generated HYAA-DobQ was incubated with Dob2, malonyl-CoA, Dob3, PMS, NADH/NADPH, and DBCO-acid and resulted in the production of DOBA-DBCO and DAC-DBCO, confirming the role of Dob2 in this pathway (**Supplementary** Figure 16).

**Figure 3:**
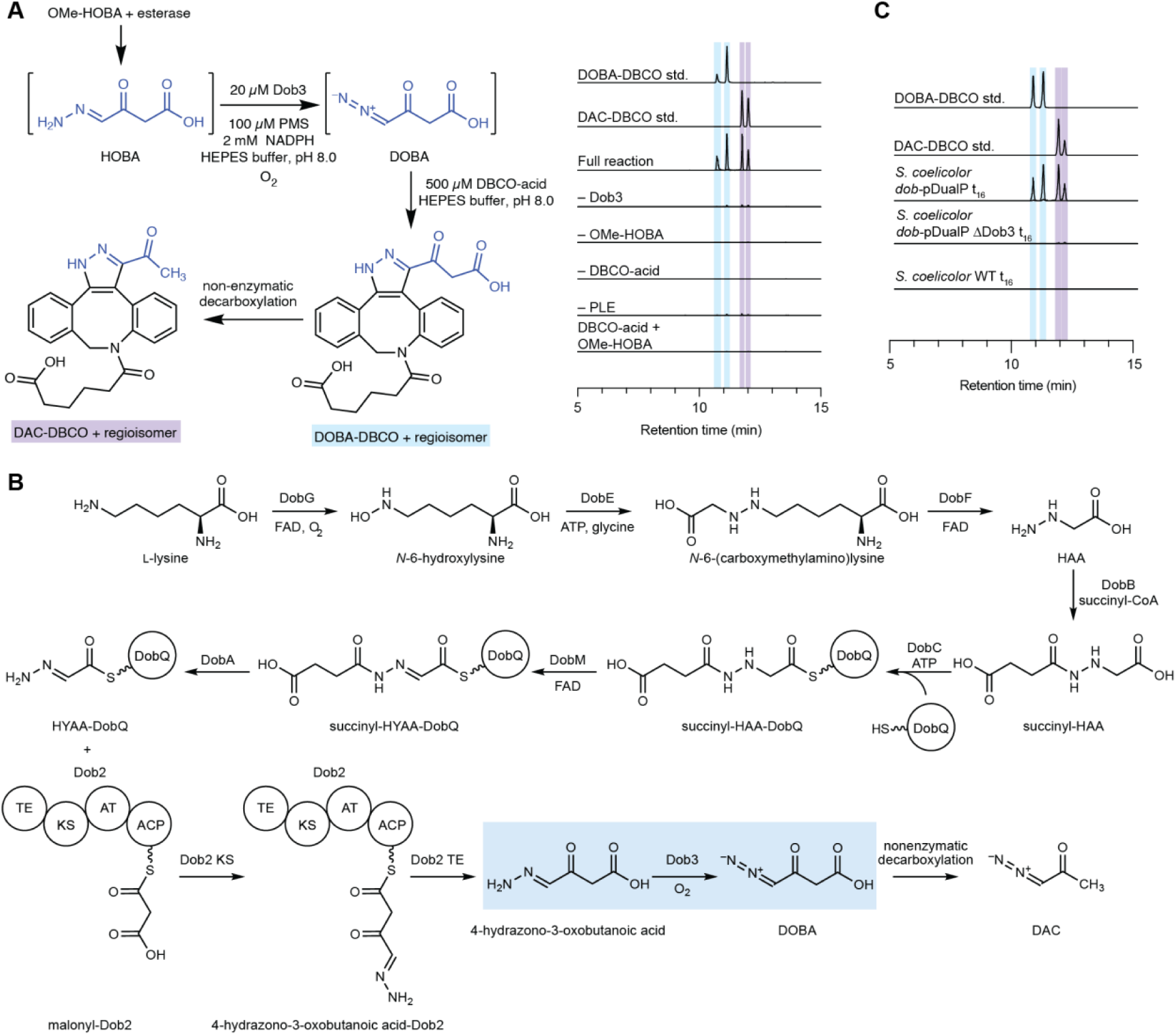
Reconstitution of Dob3 activity confirms hydrazone oxidation as a novel diazo-forming biosynthetic strategy. **A)** Overnight incubation of Dob3 with OMe-HOBA, PMS, PLE, NADPH, and DBCO-acid at pH 8.0 yields DOBA and DAC, demonstrating the Dob3 catalyzes *N*-oxidation of free HOBA to produce DOBA. Extracted ion chromatograms of *m/z* = 418.1761 ± 5 ppm and *m/z* = 462.1660 and ± 5 ppm. EICs for 462.1660 and 418.1761 are superimposed. The DOBA-DBCO standard was prepared by incubation of OMe-DOBA with DBCO-acid and PLE. Experiments were performed in biological triplicate. Representative results are shown. **B)** Proposed biosynthetic pathway of DOBA based on in vitro biochemical experiments. **C)** Extracted ion chromatograms of *m/z* = 418.1761 ± 5 ppm and *m/z* = 462.1660 ± 5 ppm in probe-treated *Streptomyces coelicolor dob*-pDualP ΔDob3 spent media at t = 16 h compared to probe-treated *Streptomyces coelicolor dob*-pDualP. t_0_ and t_16_ data was normalized to the t_16_ maximum separately for DOBA-DBCO and DAC-DBCO. EICs for 462.1660 and 418.1761 are superimposed. All experiments were performed in biological triplicate. Representative results are shown.

To evaluate the importance of Dob3 in DOBA/DAC biosynthesis in vivo, we designed the heterologous expression vector *dob*-pDualP ΔDob3 in which Dob3 is replaced with an ampicillin resistance gene (**Supplementary** Figure 17). The *dob*-pDualP ΔDob3 vector was transferred into *S. coelicolor* M1152 via conjugation, and the *S. coelicolor dob-*pDualP ΔDob3 strain was cultured to assess DOBA and DAC production. Spent media was derivatized with DBCO-acid, and LC– MS analysis demonstrated loss of DOBA-DBCO and DAC-DBCO (**Figure 3B**), supporting the necessity of Dob3 for DOBA/DAC biosynthesis in vivo.

To probe the timing of diazo formation in DOBA/DAC biosynthesis, we investigated whether Dob3 could oxidize HYAA-DobQ prior to C–C bond formation by Dob2. Dob3 and PMS were incubated with the HYAA biosynthetic enzymes and DBCO-acid in the absence of Dob2. However, no pyrazole products were detected. This suggests that C–C bond-formation by Dob2 precedes diazo formation by Dob3 (**Figure 3C**). We next attempted to probe the ability of Dob3 to oxidize PKS-tethered HOBA (HOBA-Dob2) using Ppant ejection and whole protein LC–MS assays. However, we did not detect the expected diazo product, potentially due to intermediate instability. Attempts to trap labile intermediates using DBCO-acid were also unsuccessful. As the ability of Dob3 to oxidize HOBA-Dob2 cannot be ruled out, the order of thioester hydrolysis versus diazo formation during DOBA/DAC biosynthesis remains uncertain.

### Dob3 is a unique FDO enzyme

We next sought to explore the biochemical basis for the biosynthetically unprecedented hydrazone oxidation reaction catalyzed by the FDO, Dob3. Members of the FDO family have been demonstrated to catalyze a variety of C–H oxygenation, C–H desaturation/oxygenation, and N– H oxygenation reactions. Reactions catalyzed by characterized FDO *N*-oxygenases include 4-electron and 6-electron oxidation of amines to produce nitroso,^60–62^ nitro,^63–69^ and nitronate^70^ functional groups, as well as 4-electron oxidation of hydrazines to produce the azoxy^71,72^ functional group. However, no FDO *N*-oxygenase has previously been described to catalyze 2-electron oxidation of the hydrazone functional group.

Structural predictions with AlphaFold suggested Dob3 is likely dimeric (**Supplementary** Figure 18), consistent with size-exclusion chromatography of purified Dob3 (**Supplementary** Figure 19). Comparison of the predicted structure of Dob3 to known FDO structures revealed the six conserved metal-binding residues present in all FDOs (E101, E137, H140, E198, E229, H232) as well as a seventh conserved histidine residue characteristic of FDO *N*-oxygenases (H225) (**Figure 4A**, **Supplementary** Figure 20).^73^

**Figure 4:**
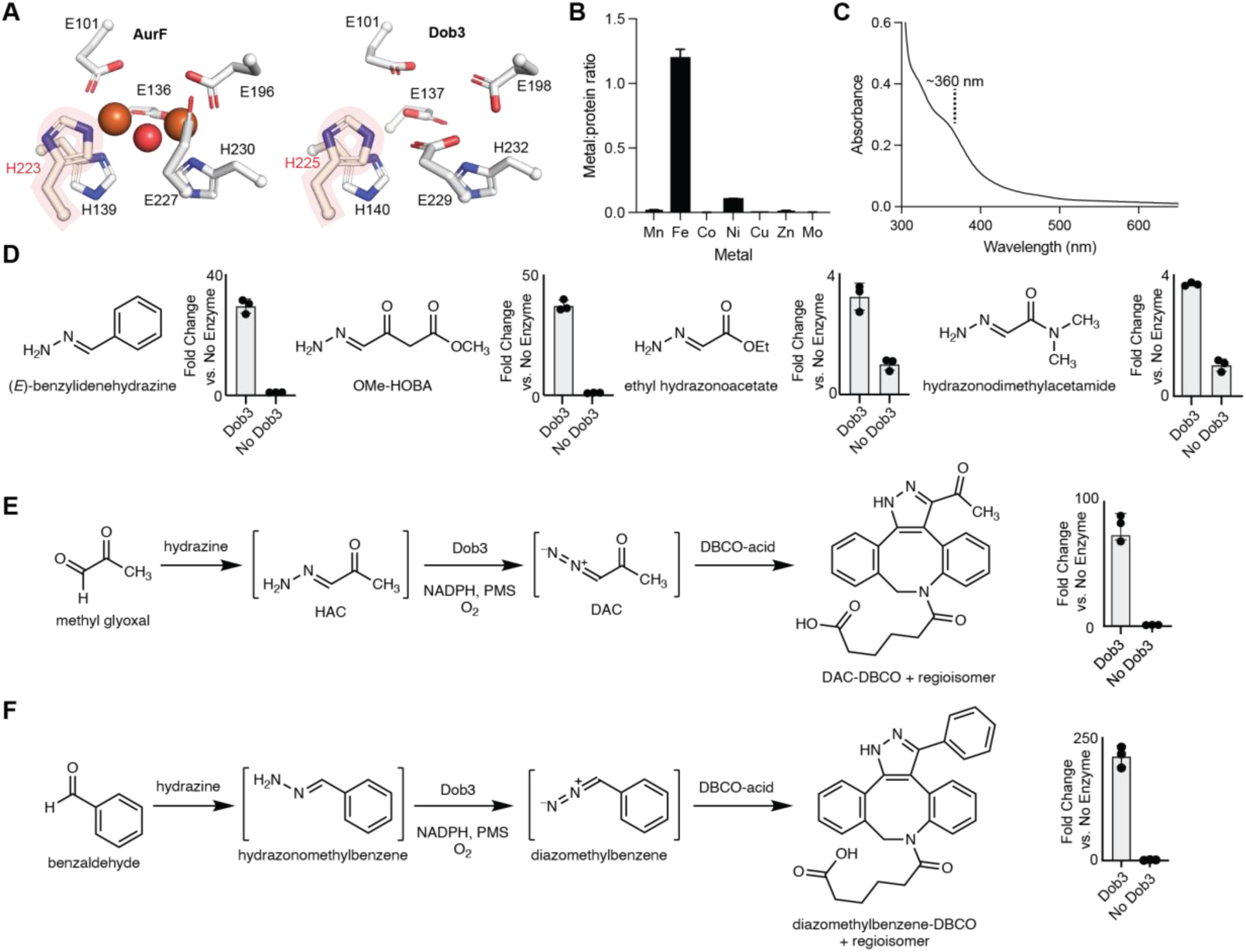
Dob3 catalyzes hydrazone oxidation. **A)** Comparison of active site residues in the AurF crystal structure and Dob3 AlphaFold structure. **B)** ICP-MS reveals Fe is the most abundant metal in Dob3 and is present at 1.4 equivalent Fe per monomer of protein. **C)** UV-vis analysis of Dob3 indicates a µ-oxo-cofactor in as-isolated protein. **D)** Dob3 oxidizes varied hydrazone substrates. 1 mM hydrazone was incubated overnight with 20 µM Dob3, 100 µM PMS, 2 mM NADPH, and 500 µM DBCO-acid in HEPES buffer pH 8.0. Peak areas were calculated as the sum of the regioisomer peak areas. **E)** One-pot incubation of methyl glyoxal, hydrazine, Dob3, and DBCO-acid produces DAC-DBCO. 1 mM aldehyde was incubated overnight with 1 mM hydrazine hydrate, 20 µM Dob3, 100 µM PMS, 2 mM NADPH, and 500 µM DBCO-acid in HEPES buffer pH 8.0. Peak areas were calculated as the sum of the regioisomer peak areas. **F)** One-pot incubation of benzaldehyde, hydrazine, Dob3, and DBCO-acid produces diazomethylbenzene-DBCO. 1 mM aldehyde was incubated overnight with 1 mM hydrazine hydrate, 20 µM Dob3, 100 µM PMS, 2 mM NADPH, and 500 µM DBCO-acid in HEPES buffer pH 8.0. Peak areas were calculated as the sum of the regioisomer peak areas.

FDOs employ dimetal cofactors such as diiron or mixed Mn/Fe cofactors.^59,65,73–76^ To characterize the active cofactor of Dob3, we attempted to reconstitute apo-Dob3 with various metals. However, expression of Dob3 in minimal medium lacking metal supplementation or in minimal medium supplemented with manganese resulted in insoluble protein. Similarly, expression of variants in which each of the putative metal-binding residues were mutated to alanine resulted in insoluble protein, suggesting the importance of metal binding for Dob3 stability. Inductively coupled plasma mass spectrometry (ICP-MS) of the as-isolated protein revealed that iron was the most abundant metal, and 1.4 equivalents of Fe per Dob3 monomer were present (**Figure 4B**). UV-vis spectroscopy of as-isolated Dob3 revealed a broad feature at ∼360 nm, consistent with a µ-oxo-diferric cofactor and analogous to the UV-vis spectra of as-isolated AurF and VlmB (**Figure 4C**).^65,71,72^ Together, these data strongly suggest that Dob3 employs a diiron cofactor. We hypothesize that the diiron cofactor reacts with molecular O_2_ to produce an intermediate which either hydroxylates the terminal nitrogen atom of a hydrazone intermediate followed by dehydration or promotes direct dehydrogenation to produce the diazo functional group (**Supplementary** Figure 21).

The synthetic utility of diazo compounds inspired us to investigate the substrate scope of Dob3 to assess its early potential as a diazo-forming biocatalyst. We tested the activity of Dob3 toward structurally varied hydrazones including ⍺-hydrazonoketones, ⍺-hydrazonoesters, ⍺-hydrazonoamides, and arylhydazones, using derivatization with DBCO-acid to facilitate product detection. Conversion to the corresponding pyrazole product was observed for all substrates to varying degrees, demonstrating the potential of Dob3 to oxidize a variety of hydrazone substrates (**Figure 4D**). We also showed that Dob3 could oxidize hydrazones generated in situ from condensation of hydrazine and commercially available aldehydes (**Figure 4E and F**). Low levels of non-enzymatic hydrazone oxidation were observed for some substrates (ranging from 3.4% to 31% of enzymatic levels), (**Supplementary** Figure 22 **and Supplementary** Figure 23).^77–79^ Overall, Dob3’s activity toward additional substrates, including precursors to important chemical reagents like ethyldiazoacetate, highlights its promise as a starting point for engineering diazo producing biocatalysts.

## Discussion

Microbes produce a variety of natural products that contain chemically reactive functional groups, and such molecules often have potent biological activity and unusual structural features that may be assembled using unprecedented enzymatic chemistry. However, traditional workflows for natural product discovery may not be suitable for identifying and characterizing reactive microbial metabolites. A prominent recent example is colibactin, a chemically unstable microbial genotoxin linked to colorectal cancer that, despite almost two decades of effort, has still not been isolated and structurally characterized using conventional methods.^80–82^ It is likely that many additional chemically reactive microbial natural products have eluded detection. Developing improved methods to enable the discovery of such natural products could reveal novel bioactive metabolites and biosynthetic enzymes.

Previously, reactivity-based screening approaches have been used to detect metabolites containing labile functional groups including ⍺,β-unsaturated carbonyls,^35,83–86^ polyynes,^36–38^ isonitriles,^40,40^ epoxides,^84^ azoxys,^87^ and reactive carbonyls.^88,89^ We have demonstrated that reactivity-based screening strategies can also be applied in the context of highly reactive diazo-containing natural products. Our approach addresses long-standing challenges of instability that have hindered the study of these metabolites. Notably, our workflow identified the novel diazo-containing natural products DOBA and DAC from the human pathogen *N. ninae*, enabling elucidation of their biosynthesis. These diazo-containing natural products could not be directly detected from microbial cultures but reacted with strained cyclooctynes to yield stable pyrazole derivatives that were readily identified using LC–MS-based comparative metabolomics.

Previously characterized diazo-containing metabolites have potent biological activity and may play important biological roles for producing microbes. The importance of these natural products is further emphasized by the phylogenetic and ecological diversity of strains containing putative diazo biosynthetic gene clusters, including microbes found in soil, aquatic environments, and infections of plants, insects, and mammals. The potent cytotoxicity of diazo-containing natural products such as lomaiviticin, azaserine, and 6-diazo-5-oxo-L-norleucine may indicate a role in microbe-microbe or microbe-host competition. DOBA and DAC are the first diazo-containing natural products identified from a human pathogen and their discovery raises interesting questions about the biological roles of these metabolites in virulence and human disease.

The discovery of diazo-containing natural products also has broader relevance to synthetic chemistry. Carbene transfer reactions using engineered heme proteins are a powerful emerging class of stereoselective biocatalytic transformations.^12,13,23,24^ Recent work on the integration of unnatural carbene transfer reactions into microbial biosynthesis demonstrated the production of chiral cyclopropanes from glucose, presenting an economically and environmentally attractive approach for pharmaceutical bioproduction.^27^ However, metabolic engineering using unnatural carbene transfer pathways is currently limited by the small number of diazo-containing natural products that can act as carbene donors.^26^ DAC is a powerful and extensively utilized synthetic reagent which has previously enabled carbene transferase-catalyzed reactions including C–H functionalization of pyrrolidines^90^ and alkene cyclopropanation en route to the antiviral Grazoprevir.^91^ The DAC biosynthetic enzymes described here can now be deployed in engineered biosynthetic carbene transfer pathways with potential impacts in renewable biomanufacturing.

Characterizing DOBA/DAC biosynthesis led to the discovery of Dob3, a new member of the FDO enzyme family and the first diazo-forming biosynthetic enzyme to catalyze 2-electron *N*-oxidation of hydrazones. Elucidation of this pathway provides biochemical confirmation that hydrazone oxidation is employed to produce diazo-containing natural products, as previously proposed for azaserine biosynthesis.^31–33^ However, the *aza* biosynthetic gene cluster lacks an encoded FDO, suggesting a different enzyme family catalyzes hydrazone oxidation in this pathway.

Wild-type Dob3 provides a promising starting place for developing engineered biocatalysts to produce synthetically useful diazo compounds from readily accessible hydrazone intermediates. This enzymatic activity was unknown prior to recent work describing the use of vanadium-dependent haloperoxidase (VHPO) enzymes to oxidize non-native hydrazone substrates.^92^ These enzymes are hypothesized to activate the terminal hydrazone nitrogen via H_2_O_2_-dependent halogenation, followed by elimination to form the diazo compound. We hypothesize that diazo formation catalyzed by Dob3 follows a distinct chemical logic whereby Dob3 uses a diiron cofactor and molecular O_2_ to oxidize the hydrazone, potentially through *N*-oxygenation followed by dehydration, or direct dehydrogenation. The discovery of Dob3 expands the scope of transformations performed by FDO enzymes and illustrates how biosynthetic pathways that construct reactive natural products are rich sources of unappreciated enzymatic reactions with the potential to enable chemical bioproduction.

Finally, our work suggests that many chemically reactive microbial natural products may have eluded discovery and characterization due to their instability. Natural product biosynthetic gene clusters predicted to generate reactive functional groups are found in numerous microbial strains including human pathogens, suggesting that reactive microbial metabolites play important yet underappreciated biological roles. We anticipate that additional reactivity-guided approaches may facilitate further exploration of this microbial ‘dark matter’ for natural product and biosynthetic enzyme discovery.

## Methods

### General experimental procedures

Primer synthesis and DNA sequencing was performed by Genewiz (Waltham, MA). DNA purification of recombinant DNA purification of recombinant plasmids was performed using an EZNA Plasmid DNA Mini Kit from Omega Bio-Tek (Norcross, GA). Restriction enzymes were purchased from New England Biolabs (Ipswich, MA) and digests were performed according to the manufacturer’s protocol. Gibson Master Mix was purchased from New England Biolabs (Ipswich, MA) and Gibson Assembly was performed according to the manufacturer’s protocol. Nickel nitriloacetic acid agarose (Ni-NTA) resin was purchased from Qiagen (Germantown, MD) and ThermoFisher Scientific (Waltham, MA). Benzylidenehydrazine was purchased from AstaTech (Bristol, PA). NADPH was purchased from Fisher Scientific (Hampton, NH). DBCO-acid was purchased from Conju-Probe (San Diego, CA) Novex Tris-Glycine SDS-PAGE gels were purchased from ThermoFisher Scientific (Waltham, MA). Protein concentrations were determined by measuring UV-absorption at 280 nm using a Thermo Scientific NanoDrop 2000. ExPASy ProtParam was used to calculate extinction coefficients. Optical densities of *E. coli* cultures were measured at 600 nm using a Beckman Coulter DU730 Life Sciences UV/Vis spectrophotometer. All water used was purified using a MilliQ water purification system. The remaining materials were purchased from Sigma-Aldrich unless otherwise indicated.

### General methods for mass spectrometry

#### Agilent 6530 qTOF spectrometer with Dual AJS source

Unless otherwise indicated, samples were analyzed by LC–MS using an Agilent 1200 series LC system coupled to an Agilent 6530 qTOF spectrometer with Dual AJS source. Drying gas temperature was 300 °C, drying gas flow was 11 L/min, nebulizer pressure was 45 psi, sheath gas temperature was 275 °C, sheath gas flow was 11 L/min, capillary voltage was 3500 V, nozzle voltage was 500 V, fragmentor voltage was 125 V, and skimmer voltage was 65 V. Mass spectra were acquired in positive ion mode. The LC column was a Cogent Diamond Hydride column (4 µm, 100 Å, 3 × 150 mm, Microsolv Technology Corp). The flow rate was 0.5 mL/min using solvent A = 0.1% formic acid in water and solvent B = 0.1% formic acid in acetonitrile. The LC conditions were: 0–1 min at 70% B isocratic, 1–7 min at 70–50% B, 7–9 min at 50% B isocratic, 9–10 min at 50–70% B, 10–14 min at 70% B isocratic. MS/MS spectra were collected with 10V collision energy. A mass window of 10 ppm was used for extracted ion chromatograms.

#### Thermo Orbitrap IQ-X Tribrid

Unless otherwise indicated, samples were analyzed using a Horizon Vanquish UHPLC coupled to a Thermo Orbitrap IQ-X Tribrid mass spectrometer. The LC column was a Kinetex C18 column (1.7 µm, 100 Å, 150 x 2.1 mm, Kinetex). 2 µL of sample was injected. The flow rate was 0.4 mL/min using mobile phase A = 0.1% formic acid in water and B = 0.1% formic acid in acetonitrile. The LC conditions were 0-2 min 10% B isocratic, 2-29 min 10-90% B, 29-32 min 90% B isocratic, 32-32 min 90-10% B, 32-35 min 10% B isocratic. The MS settings were: mass range 400-600 *m/z,* 120000 resolution, the RF lens was 35%, the standard AGC target was used, and the auto maximum injection time mode was used. MS/MS spectra were acquired with a 1 *m/z* isolation window, 15000 resolution, standard AGC target, auto maximum injection time mode, and HCD fragmentation with 30% collision energy. Spectra were acquired in positive mode. A mass window of 5 ppm was used for extracted ion chromatograms.

#### Comparative Metabolomics using AcquireX and Compound Discoverer

Data-dependent MS/MS acquisition was performed via AcquireX (deep scan mode) from 4 replicate injections with media + DBCO-acid providing an exclusion list. Data-dependent MS/MS spectra were acquired with 2.0E4 intensity threshold, exclusion after 1 time, 2.5 s exclusion duration, a 3 ppm isolation window, and HCD, CID, and UVPD fragmentation. HCD fragmentation was performed with a 1 m/z isolation window, stepped collision energies of 20, 35, 50, 75, and 100%, and 15,000 resolution. CID fragmentation was performed with a 1 m/z isolation window, 30% collision energy, 15,000 resolution, and 22 ms maximum injection time. UVPD fragmentation was performed with a 1.6 *m/z* isolation window, molecular weight-dependent UVPD activation time, 12% activation time, 15,000 resolution, and 22 ms maximum injection time.

Comparative metabolomics for t_0_ and t_16_ samples were performed using Compound Discoverer: The raw spectra of all samples were imported into Compound Discoverer 3.3 (Thermo Scientific) where peak alignment, compound detection, and compound grouping was performed. Compound detection was performed with a 3 ppm mass tolerance, a 10,000 minimum peak intensity, 1.5 chromatographic signal/noise threshold, and isotope grouping for Br and Cl ions. All ions were considered. Compound grouping was performed with a 3 ppm mass tolerance, a 0.2 min retention time tolerance, preferred ions of [M+H]^+^ and [M–H]^-^ and area integration of the most common ion. Peak rating contributions were 3 for area contribution, 10 for CV contribution, 5 for FWHM to Base Contribution, 5 for Jaggedness Contribution, 5 for Modality Contribution, and 5 for Zig-Zag Index Contribution. Peak rating filters were 3 for Peak Rating Threshold and 2 for Number of Files. Gap filling was performed with a mass tolerance of 5 ppm and a signal to noise threshold of 1.5. SERRF QC Correction was applied with a 50% minimum QC coverage, a 30% maximum QC area RSD, a 25% maximum corrected QC Area RSD, 2 batches, and interpolated gap-filling. Background compounds were marked with a max sample/blank ratio of 5, a max blank/sample ratio of 0 and the background compounds were then hidden. MS spectra were assigned as t_16_ or t_0_ and differential analysis was performed following spectra processing. Differential analysis was performed with area values transformed to log10. Peak rating contributions for differential analysis were 3 for area contribution, 10 for CV contribution, 5 for FWHM to Base Contribution, 5 for Jaggedness Contribution, 5 for Modality Contribution, and 5 for Zig-Zag Index Contribution. Following statistical analysis, results were filtered to include only those with a calculated molecular weight and *m/z* greater than 333.

### Reaction of azaserine with alkyne probes

100 µL reaction mixtures containing 100 µM azaserine and 100 µM of the specified alkyne were prepared in water. The reaction mixtures were incubated for 16 h at room temperature. 10 µL of reaction mixture was then diluted with 150 µL of LC–MS H_2_O, 20 µL of LC–MS ACN, and 20 µL of MeOH and samples were filtered (0.2 µm, VWR, nylon). Detection of probe adducts was accomplished using the method described above for the Thermo Orbitrap IQ-X Tribrid mass spectrometer.

### Optimization of the reaction with azaserine and DBCO-acid: solvent screen

100 µL reaction mixtures containing 100 µM azaserine and 100 µM DBCO-acid in H_2_O, 1:1 H_2_O:MeOH, MeOH, 1:1 H_2_O:ACN, or ACN were incubated overnight at room temperature. 10 µL of the reaction mixture was then diluted with 150 µL of LC–MS H_2_O, 20 µL of LC–MS ACN, and 20 µL of MeOH and samples were filtered (0.2 µm, VWR, nylon). Detection of azaserine-DBCO was accomplished using the method described above for the Thermo Orbitrap IQ-X Tribrid mass spectrometer.

### Optimization of the reaction with azaserine and DBCO-acid: reaction time screen

100 µL reaction mixtures containing 100 µM azaserine and 500 µM DBCO-acid in H_2_O were incubated for the specified amount of time at room temperature. At the specified time point, 10 µL of the reaction mixture was diluted with 150 µL of LC–MS H_2_O, 20 µL of LC–MS ACN, and 20 µL of MeOH and samples were filtered (0.2 µm, VWR, nylon). Detection of azaserine-DBCO was accomplished using the method described above for the Thermo Orbitrap IQ-X Tribrid mass spectrometer.

### Optimization of the reaction with azaserine and DBCO-acid: DBCO-acid equivalents

100 µL reaction mixtures containing 100 µM azaserine and the specified equivalents of DBCO-acid in H_2_O were incubated overnight at room temperature. 10 µL of the reaction mixture was then diluted with 150 µL of LC–MS H_2_O, 20 µL of LC–MS ACN, and 20 µL of MeOH and samples were filtered (0.2 µm, VWR, nylon). Detection of azaserine-DBCO was accomplished using the method described above for the Thermo Orbitrap IQ-X Tribrid mass spectrometer.

### DBCO-acid substrate scope

250 µL reaction mixtures containing 100 µM diazo substrate and 100 µM DBCO-acid in H_2_O were incubated for 18 h at room temperature. 10 µL of the reaction mixture was diluted into 150 µL of H_2_O + 20 µL of MeOH + 20 µL of ACN and filtered (0.2 µm, VWR, nylon). Detection of DBCO-acid adducts were accomplished using the method described above for the Thermo Orbitrap IQ-X Tribrid mass spectrometer.

### *G. harbinensis* t_0_ vs t_16_ comparative metabolomics

*G. harbinensis* was grown on NZ amine plates (10 g/L glucose, 20 g/L soluble starch, 5 g/L yeast extract, 5 g/L N-Z-amine, 1 g/L CaCO_3_, 15 g/L agar) at 30 °C for 7-10 days and was then used to inoculate 5 mL of NZ amine medium. The NZ amine culture was incubated for 7-10 days at 30 °C with shaking at 200 rpm. 400 µL of the liquid culture was used to inoculate 20 mL of molasses production medium (10 g/L glucose, 5 g/L Bacto peptone, 20 g/L molasses, 1 g/L CaCO_3_) which was incubated at 30 °C with shaking at 200 rpm for 14 days. 200 µL of culture was spun down for 5 min at max speed. 1 µL of 25 mM DBCO-acid in MeOH was added to 49 µL of supernatant. The t_0_ control was obtained by dilution of 10 µL of the reaction mixture into 150 µL of H_2_O + 20 µL of ACN + 20 µL of MeOH, filtration (0.2 µm, VWR, nylon), and storage at –80 °C prior to analysis. The t_16_ sample was obtained by a 16 h incubation of the remaining reaction mixture at room temperature, followed by dilution of 10 µL of the reaction mixture into 150 µL of H_2_O + 20 µL of ACN + 20 µL of MeOH, and filtration (0.2 µm, VWR, nylon). Comparative metabolomics was performed using a Thermo Orbitrap IQ-X Tribrid mass spectrometer as described above.

### Genome mining for biosynthetic gene clusters encoding hydrazone *N*-oxidation

The protein sequence of AzaE was queried against the NCBI nucleotide collection database and the non-redundant protein sequence database in tblastn and blastp searches, respectively. Blastp results with an E value < E-98 that were non-redundant were added to the tblastn search results. GenBank files were downloaded using the accession number for each result. The files were prepared according to the prettyClusters^51^ workflow, detailed below. The GenBank documents were annotated with Prokka.^93^ Amino acid sequences were extracted and added to a local database. AzaE was queried against this database in a blastp search and homologs with an E-value < E-31 were taken forward. Gene metadata and neighbor sequences and metadata were generated with the gbToIMG function using default values except NeighborNumber was set to 15. InterProScan was used to generate additional metadata for all genes and neighbors. This data was merged with the previous metadata using the incorpIprScan function. The prepNeighbors function was run with NeighborNumber = 10 and trimShortClusters = false. AzaE homologs were dereplicated using the EFI-SSN tool and the repNodeTrim function. Manual curation identified gene neighborhoods containing homologs of hydrazone-forming enzymes (neighborhoods contained at least one representative each of pfam02770, pfam13434, pfam01266, pfam00501, pfam13302, and pfam03417. pfam00550 was not considered as acyl carrier proteins are commonly not annotated). The analyzeNeighbors function was run on the HYAA-cassette containing clusters with neighborThreshold = 2.5 and tgCutoff = 56. Results were visualized with Cytoscape.^94^ This analysis identified 113 putative diazo encoding biosynthetic gene clusters. Compilation of these results with bioinformatic analysis from previous studies resulted in Figure 1C.

### Diazo-containing natural product discovery from *N. ninae*

*N. ninae* was grown on GYM agar (4 g/L glucose, 4 g/L yeast extract, 10 g/L malt extract, 2 g/L CaCO_3_, 20 g/L agar) plates for 7-10 days at 30 °C. Colonies were used to inoculate 5 mL of GYM medium (4 g/L glucose, 4 g/L yeast extract, 10 g/L malt extract), and the culture was incubated for 7-10 days at 30 °C with shaking at 200 rpm. 100 µL of the culture was used to inoculate 5 mL of GYM medium and this was incubated for 7-10 days at 30 °C with shaking at 200 rpm. 200 µL of spent culture medium was spun down at max speed for 5 min. 1 µL of 25 mM DBCO-acid in MeOH was added to 49 µL of spent culture medium and incubated overnight at room temperature. 10 µL of the reaction mixture was diluted into 150 µL of H_2_O + 20 µL of ACN + 20 µL of MeOH and filtered (0.2 µm, VWR, nylon). Samples were analyzed using a Thermo Orbitrap IQ-X Tribrid mass spectrometer as described above with minor modifications. The MS settings were: mass range 70-700 *m/z*. Comparative metabolomics for t_0_ and t_16_ samples were performed using Compound Discoverer as described in the general methods above.

### Detection of DOBA-DBCO and DAC-DBCO from *N. tenerifensis*

*N. tenerifensis* was grown on GYM agar plates for 7-10 days at 30 °C. Colonies were used to inoculate 5 mL of GYM medium and the culture was incubated for 7-10 days at 30 °C with shaking at 200 rpm. 100 µL of the culture was used to inoculate 5 mL of molasses production medium, and this culture was incubated for 7-10 days at 30 °C with shaking at 200 rpm. 200 µL of spent culture medium was spun down at max speed for 5 min. 1 µL of 25 mM DBCO-acid in MeOH was added to 49 µL of spent culture medium and incubated overnight at room temperature. 10 µL of the reaction mixture was diluted into 150 µL of H_2_O + 20 µL of ACN + 20 µL of MeOH and filtered (0.2 µm, VWR, nylon). Detection of DAC-DBCO and DOBA-DBCO was accomplished using a Thermo Orbitrap IQ-X Tribrid mass spectrometer as described above with minor modifications. The MS settings were: mass range 70-700 *m/z*. MS/MS was accomplished using HCD fragmentation with 30% normalized collision energy or CID fragmentation with 30% normalized collision energy, 10 ms activation time, and 0.25 activation Q for DAC-DBCO and DOBA-DBCO, respectively.

### Detection of DOBA-DBCO and DAC-DBCO from *S. coelicolor dob*-pDualP

*S. coelicolor dob-*pDualP was grown on MSF (20 g/L mannitol, 20 g/L soy flour, 20 g/L agar) + nalidixic acid (30 µg/mL) + apramycin (50 µg/mL) agar plates at 30 °C for 5-7 days and then used to inoculate 5 mL of YEME medium. *S. coelicolor* WT was grown on MSF agar plates at 30 °C for 5-7 days and used to inoculate 5 mL of YEME medium. The YEME cultures were incubated at 30 °C with shaking at 200 rpm for 7 days. 100 µL of the YEME cultures was used to inoculate 5 mL of molasses production medium which was incubated at 30 °C with shaking at 200 rpm for 14 days. ε-caprolactam and oxytetracycline were added to the cultures for final concentrations of 0.1% w/v and 2.5 mM respectively. After incubation of the induced cultures at 30 °C with shaking at 200 rpm for 7 days, 200 µL of spent culture medium was derivatized and analyzed as described above with minor modifications. The MS settings were: mass range 400-480 *m/z*. MS/MS spectra were acquired with a 5 ppm isolation window, 15000 resolution, 1000% AGC target, dynamic maximum injection time, and HCD fragmentation with 30% normalized collision energy or CID fragmentation with 30% collision energy, 10 ms activation time, and 0.25 activation Q for DAC-DBCO and DOBA-DBCO, respectively.

### Cloning *dob-*pDualP

Construction of the *dob*-pDualP vector was performed as previously reported for *aza*-pDualP, with minor modifications.^31^ Cloning of *dob*-pDualP was performed by Terra Bioforge (Madison, WI). *N. ninae* cells were lysed using proprietary methods to maintain genomic DNA integrity. Genomic DNA >50 kb was isolated using proprietary gel free methods. The *dob* gene cluster was excised from *N. ninae* genomic DNA using Cas9 with guide RNAs targeting upstream and downstream of the gene cluster. A *Streptomyces* compatible dual-inducible vector (pDualP) containing Potr^95^ (oxytetracycline) and PnitA^96,97^ (ε-caprolactam) inducible promoters flanking the insertion site was amplified using PCR with primers designed to create ∼40 bp of overlap specific to the dob gene cluster. Modified Gibson Assembly of the *dob* gene cluster and PCR amplified linear pDualP was performed using a proprietary reaction mix to produce the *dob*-pDualP expression vector. The *dob*-pDualP vector was transformed into *E. coli* BacOpt2.0 (*E. coli* DH10b derivative) and clones were verified by whole plasmid sequencing.

### Cloning *dob-*pDualP ΔDob3

The deletion of Dob3 was performed by Terra Bioforge (Middleton, WI). Briefly, *dob-*pDualP was restricted *in vitro* with CRISPR nuclease complexed with 2 guide RNAs targeting both the 5’ and 3’ ends of the FDO gene. The digested DNA was then repaired in an isothermal assembly reaction with the AmpR cassette from pUC19, which was amplified by PCR using the primers gaaatggagggatagccgtgtaagtttaaacggcacttttcgggg and aaattggacagcagccgcttgtttaaacgttaccaatgcttaatcagtgagg. The assembly reaction to remove the FDO gene and replace it with the AmpR cassette was then transformed via electroporation to *E. coli* DH10B. The transformation culture was then plated on LB agar containing 25 µg/mL apramycin and 100 µg/mL ampicillin. Transformant colonies were picked and plasmid DNA extracted. Whole Plasmid Sequencing was performed by Plasmidsaurus using Oxford Nanopore Technology with custom analysis and annotation.

### Conjugation of *dob*-pDualP into *S. coelicolor* M1152

*S. coelicolor* M1152 was grown on MSF agar for 7 days at 30 °C. A single colony was used to inoculate 5 mL of YEME (3 g/L yeast extract, 3 g/L malt extract, 5 g/L peptone, 10 g/L glucose, 340 g/L sucrose) medium which was incubated at 30 °C with shaking at 200 rpm for 7 days. 5 mL of LB + apramycin (50 µg/mL) + kanamycin (50 µg/mL) + chloramphenicol (20 µg/mL) medium was inoculated with *E. coli* ET12567/pUZ8002 and grown at 37 °C with shaking at 200 rpm for 2 days. The culture was spun down at 3000 rpm for 10 min. 5 mL of LB medium was added and the pellet was resuspended. The culture was spun down at 3000 rpm for 10 min and the supernatant was discarded. This wash step was then repeated before the pellet was resuspended in 100 µL of LB medium. The *S. coelicolor* culture was spun down at 4000 rpm for 10 min on the same day. 2 mL of 10% sucrose was added, the pellet was resuspended, and the culture was spun down at 4000 rpm for 10 min. This was repeated before the pellet was resuspended in 100 µL of LB. 100 µL of *S. coelicolor* suspension was combined with 100 µL of *E. coli* suspension and mixed. This was plated on MSF + 10 mM MgCl_2_ agar and grown at 30 °C. After 18 h, the plate was flooded with 1 mL of sterile water containing 1 mg apramycin and 0.5 mg nalidixic acid before being incubated at 30 °C. Ex-conjugants were grown on MSF + apramycin (50 µg/mL) + nalidixic acid (30 µg/mL) agar and confirmed by genome sequencing.

### Conjugation of *dob*-pDualP ΔDob3 into *S. coelicolor* M1152

5 mL of LB + apramycin (25 µg/mL) + chloramphenicol (10 µg/mL) + kanamycin (25 µg/mL) medium was inoculated with *E. coli* ET12567/pUZ8002 *dob*-pDualP ΔDob3 and grown at 37 °C with shaking at 200 rpm overnight. After the culture reached OD_600_ >0.4, it was spun down at 3000 rpm for 10 min and the supernatant was discarded. 5 mL of LB medium was added, and the pellet was resuspended. The culture was spun down at 3000 rpm for 10 min and the supernatant was discarded. This wash step was repeated before the pellet was resuspended in 500 µL of LB medium. 10 µL of *S. coelicolor* M1152 spore stock was added to 500 µL of 2xYT medium and incubated for 10 min at 50 °C. Samples were then cooled to room temperature. 500 µL of *E. coli* ET12567/pUZ8002 *dob*-pDualP ΔDob3 in LB medium was added to 500 µL of *S. coelicolor* in 2xYT (16 g/L casein digest peptone, 10 g/L yeast extract, 5 g/L NaCl). Samples were spun down at 3000 rpm for 10 min and 850 µL of supernatant was removed. The pellet was resuspended in the remaining supernatant and plated on MSF + 50 mM MgCl_2_ agar. The plate was then incubated at 30 °C for 24 h before it was flooded with 1 mL of sterile water containing 1 mg apramycin and 0.5 mg nalidixic acid. Ex-conjugants were grown on MSF + apramycin + nalidixic acid agar and confirmed by genome sequencing.

### Genome sequencing verification of *S. coelicolor dob-*pDualP ΔDob3

*S. coelicolor dob*-pDualP and *S. coelicolor dob*-pDualP ΔDob3 were grown on MSF + apramycin agar for 7 days at 30 °C. Single colonies were used to inoculate 5 mL of YEME medium (3 g/L yeast extract, 3 g/L malt extracat, 5 g/L peptone, 10 g/L glucose, 340 g/L sucrose) which was incubated for 7 days at 30 °C with shaking at 200 rpm. Genomic DNA was extracted using the Lucigen MasterPure Gram Positive gDNA Purification Kit. Samples were incubated with Ready-Lyse lysozyme for 36 h and eluted with 10 µL of H_2_O. Genomic DNA sequencing was performed by Plasmidsaurus using Oxford Nanopore Technology with custom analysis and annotation.

### Comparison of *S. coelicolor dob*-pDualP vs *S. coelicolor dob*-pDualP ΔDob3 spent media

*S. coelicolor dob*-pDualP and *S. coelicolor dob*-pDualP ΔDob3 were grown on MSF + apramycin agar for 7 days at 30 °C. Single colonies were used to inoculate 5 mL of YEME medium which were incubated for 7 days at 30 °C with shaking at 200 rpm. 100 µL of starter culture was used to inoculate 5 mL of ISP4 medium (10 g/L soluble starch, 1 g/L MgSO_4_ • 7H_2_O, 1 g/L NaCl, 2 g/L (NH_4_)_2_SO_4_, 2 g/L CaCO_3_ 1 L H_2_O, 1 mL of trace salts solution (stock solution: 1 g/L FeSO_4_ • 7H_2_O, 1 g/L MnCl_2_ • 4H_2_O, 1 g/L ZnSO_4_ • 7H_2_O). The ISP4 cultures were incubated for 14 days at 30 °C with shaking at 200 rpm before they were induced with ε-caprolactam (0.1% w/v) and oxytetracycline (2.5 µM) and incubated overnight at 30 °C with shaking at 200 rpm. 200 µL of culture aliquots were spun down at max speed for 5 min. 1 µL of 25 mM DBCO-acid in MeOH was added to 49 µL of spent culture medium and incubated overnight. 10 µL of the reaction mixture was diluted into 150 µL of H_2_O + 20 µL of ACN + 20 µL of MeOH and filtered (0.2 µm, VWR, nylon). Detection of DAC-DBCO and DOBA-DBCO was accomplished using a Thermo Orbitrap IQ-X Tribrid mass spectrometer coupled to a Horizon Vanquish UHPLC according to the above method with minor modifications. The mass range was 70-700 *m/z*, MS/MS spectra were acquired with a 5 ppm isolation window, 15000 resolution, standard AGC target, auto maximum injection time mode, and HCD fragmentation with 30% normalized collision energy or CID fragmentation with 30% collision energy, 10 ms activation time, and 0.25 activation Q for DAC-DBCO and DOBA-DBCO, respectively.

### Isolation of *N. ninae* genomic DNA

*N. ninae* was grown on GYM agar plates for 7-10 days at 30 °C. Single colonies were used to inoculate 5 mL of GYM medium and the culture was incubated for 7-10 days at 30 °C with shaking at 200 rpm. Genomic DNA was extracted using the Lucigen MasterPure Gram Positive gDNA Purification Kit with a 16 h lysozyme incubation. Samples were eluted with 10 µL of H_2_O.

### Cloning DobA, DobB, DobQ, DobC, DobG, DobM, Dob2, and Dob3

Genes encoding each protein were amplified by PCR using the appropriate forward and reverse primers listed in **Supplementary Table 1**. PCR products were isolated using Zymoclean Gel DNA Recovery Kit and DNA Clean & Concentrator. PCR products were ligated into a NdeI/HindIII digested pET28a plasmid using Gibson assembly. Plasmids were transformed into *E. coli* Top10 and the sequence was confirmed by Sanger sequencing. Plasmids containing the *dobA, dobB, dobC, dobG, dobM,* and *dob3* inserts were transformed into *E. coli* BL21(DE3) for protein expression. Plasmids encoding the *dobQ* (carrier protein) and *dob2* (PKS) inserts were transformed into *E. coli* BAP1 for expression of phosphopantetheinylated holoenzymes.

### Expression and Purification of DobA, DobB, DobC, DobG, DobM, DobQ, Dob2, Dob2S88A, Dob3, and Dob3 mutants

A single colony was used to inoculate 30 mL of LB medium + kanamycin (50 µg/mL) which was incubated overnight at 37 °C with shaking at 200 rpm. A 20 mL aliquot was used to inoculate 1 L of LB medium + kanamycin (50 µg/mL) culture which was incubated at 37 °C with shaking at 180 rpm until an OD_600_ of at least 0.6 was reached. Cultures were cold-shocked on ice for 30 min before induction with 50 µg/mL IPTG. For DobG and DobM, 75 mg/L riboflavin was added upon induction. Cultures were incubated at 18 °C with shaking at 180 rpm overnight. After 16 h, cells were centrifuged at 4000 rpm at 4 °C for 20 min. Pellets were resuspended in 30 mL of 20 mM HEPES pH 8.0, 10 mM MgCl_2_, and 500 mM NaCl. For DobG and DobM, 1 mM FAD was added to the lysis buffer. Cells were lysed using a cell disruptor (Avestin EmulsiFlex-C3) and lysates were spun down at 15,000 × g for 40 min at 4 °C. Spent culture medium was incubated with 3 mL of Ni-NTA resin and nutated at 4 °C for 30 min. The resin was washed with buffer containing 25 mM imidazole before eluting proteins in buffer containing 200 mM imidazole. Eluted proteins were concentrated to < 2 mL using a MilliporeSigma Amicon Ultra-15 Centrifugal Filter Unit with an appropriate molecular weight cutoff. DobB, DobC, DobG, DobM, DobQ, Dob2, Dob2S88A, Dob3, and Dob3 mutants were desalted using buffer (20 mM HEPES pH 8.0, 10 mM MgCl_2_, 50 mM NaCl, 10% glycerol) and Cytiva PD-10 columns pre-packed with Sephadex G-25 resin. DobA was desalted and further purified on a BioRad BioLogic DuoFlow FPLC system with a Superdex 75 column using 20 mM HEPES pH 8.0, 10 mM MgCl_2_, 50 mM NaCl, and 10% glycerol as the mobile phase. Desalted proteins were concentrated using a MilliporeSigma Amicon Ultra-15 Centrifugal Filter Unit with an appropriate molecular weight cutoff before storing concentrated proteins at –80 °C.

### In vitro activity assay of DobG

100 µL reaction mixtures containing 20 µM DobG, 1 mM L-lysine, 500 µM NADH, 500 µM NADPH, and 100 µM FAD were prepared in buffer (20 mM HEPES pH 8.0, 10 mM MgCl_2_, 50 mM NaCl, 10% glycerol) and incubated aerobically at room temperature for 30 minutes. Reaction mixtures were diluted with 2.3 volumes of LC–MS grade acetonitrile, incubated at –20 °C for 20 minutes, and centrifuged at 21,000 × g to remove insoluble material. Supernatants were passed through a 0.2 µM filter (VWR, 13 mm, Nylon, 0.2 µm). Samples were analyzed by LC–MS and detection of *N*-6-hydroxylysine was accomplished by LC–MS using an Agilent 1200 series LC system coupled to an Agilent 6530 quantitative time-of-flight (qTOF) mass spectrometer with a Dual Agilent Jet Stream Technology (AJS) Ionization Source using the method described above. Assays were performed in biological triplicate and representative results are shown.

### In vivo activity assay of DobE

A single colony of *E. coli* BL21(DE3) containing the *azaE* plasmid or empty pET28a plasmid (control) was used to inoculate 5 mL of LB medium + kanamycin (50 µg/mL) and incubated at 37 °C with shaking at 200 rpm. After 16 h, 2% v/v of saturated overnight culture was used to inoculate 5 mL of M9 medium (1 x M9 salt solution, 100 µM CaCl_2_, 2 mM MgSO_4_, 0.4% glucose; 5 x M9 salt solution: 64 g/L Na_2_HPO_4_, 15 g/L KH_2_PO_4_, 2.5 g/L NaCl, 5 g/L NH_4_Cl) + kanamycin (50 µg/mL). Cultures were incubated at 37 °C with shaking at 200 rpm until OD_600_ reached 0.6. Cultures were cold shocked on ice for 30 min and then 50 µg/mL IPTG, 1 mM *N*-6-hydroxylysine, 1 mM glycine, and 4 mM ATP were added. Cultures were incubated overnight at 18 °C with shaking at 200 rpm. After 16 h, samples of each culture were prepared by diluting spent culture medium to a final concentration of 70% LC–MS grade acetonitrile and incubating at –20 °C for 30 min. The insoluble material was removed by centrifugation at 16,100 *× g*, and the sample was analyzed by LC–MS. Detection was accomplished using an Agilent 1200 series LC system coupled to an Agilent 6530 quantitative time-of-flight (qTOF) mass spectrometer using the method described above.

### In vitro activity of DobB

100 µL reaction mixtures containing 50 µM DobB, 100 µM succinyl-CoA, and 100 µM HAA were prepared in buffer (20 mM HEPES pH 6.7, 10 mM MgCl_2_, 50 mM NaCl, 10% glycerol) and incubated aerobically on ice for 5 minutes. Samples were diluted to 95:5 H_2_O:ACN, filtered (3kDa, Amicon, cellulose), and analyzed using an Agilent 1200 series LC system coupled to an Agilent 6530 quantitative time-of-flight (qTOF) mass spectrometer with a Dual Agilent Jet Stream Technology (AJS) Ionization Source. Drying gas temperature was 300 °C, drying gas flow was 11 L/min, nebulizer pressure was 45 psi, sheath gas temperature was 275 °C, sheath gas flow was 11 L/min, capillary voltage was 3500V, nozzle voltage was 500V, fragmentor voltage was 125V, and skimmer voltage was 65V. The LC column was an Eclipse XDB C-18 column (5 µm, 80 Å, 4.6 × 150 mm, Agilent). The flow rate was 0.5 mL/min using solvent A = 0.1% formic acid in water and solvent B = 0.1% formic acid in acetonitrile. The LC conditions were: 0–5 min at 5% B isocratic, 5–8 min at 5–95% B, 8–10 min at 95% B isocratic, 10–12 min at 95–5% B, 12–18 min at 5% B isocratic. A mass window of 10 ppm was used for extracted ion chromatograms. Assays were performed in biological triplicate and representative results are shown.

### PPANT ejection assay of DobA, DobB, DobC, DobM, DobQ

50 µL reaction mixtures containing 0.2 mM succinyl-CoA, 1 mM hydrazinoacetic acid (HAA), 5 mM ATP, 0.2 mM FAD, 40 µM DobA, 20 µM DobB, 20 µM DobC, 20 µM DobM, and 50 µM DobQ in exchange buffer (20 mM HEPES pH 8.0, 10 mM MgCl_2_, 50 mM NaCl, 10% glycerol) were incubated for 20 min at room temperature. Reaction mixtures were immediately diluted with 4 volumes of chilled water and kept on ice to slow reaction progress. Samples were immediately filtered (0.2 µm, VWR, nylon) and analyzed by LC–MS using an Agilent 1200 series LC system coupled to an Agilent 6530 qTOF spectrometer with Dual AJS source. Drying gas temperature was 325 °C, drying gas flow rate was 10 L/min, nebulizer pressure was 35 psi, sheath gas temperature was 275 °C, sheath gas flow was 11 L/min, capillary voltage was 4000 V, nozzle voltage was 1000 V, fragmentor voltage was 250 V, and skimmer voltage was 65 V. Mass spectra were acquired in positive ion mode. The LC column was an Aeries WIDEPORE XB-C18 column (3.6 µm, 200 Å, 4.6 x 150 mm, Phenomenex). 10 µL of sample was injected. The flow rate was 0.5 mL/min with solvent A = 0.1% formic acid in water and solvent B = 0.1% formic acid in acetonitrile. The LC conditions were: 1-5 min at 5% B isocratic, 5-23 min at 5-60% B, 23-26 min at 60-95% B, 26-27 min at 95% B isocratic, 27-28 min at 95-5% B, 28-35 min at 5% B isocratic. A mass window of 10 ppm was used for extracted ion chromatograms.

### In vitro activity of Dob3

50 µL reaction mixtures containing 100 µM PMS, 20 µM Dob3, 1 mM OMe-HOBA, 2 mM NADPH, 0.1 mg PLE, and 500 µM DBCO-acid in exchange buffer (20 mM HEPES pH 8.0, 10 mM MgCl_2_, 50 mM NaCl, 10% glycerol) were incubated overnight. 70 µL of buffer, 15 µL of ACN, and 15 µL of MeOH were added, and proteins were removed by 3 kDa centrifugal filter prior to LC–MS analysis. Detection of DOBA-DBCO and DAC-DBCO was accomplished using a Thermo Orbitrap IQ-X Tribrid mass spectrometer as described above with minor modifications. The MS settings were: mass range 70-700 *m/z*. MS/MS spectra of DAC-DBCO were obtained using HCD fragmentation with 30% normalized collision energy. MS/MS spectra of DOBA-DBCO were obtained using CID fragmentation with 30% collision energy, 10 ms activation time, and 0.25 activation Q.

### DobA, DobB, DobC, DobM, DobQ, Dob2, and Dob3 cascade reaction

50 µL reaction mixtures containing 1 mM HAA, 20 µM DobB, 200 µM succinyl-CoA, 50 µM DobQ, 20 µM DobC, 5 mM ATP, 20 µM DobM, 200 µM FAD, 40 µM DobA, 10 µM Dob2, 200 µM malonyl-CoA 20 µM Dob3, 100 µM phenazine methosulfate (PMS), 2 mM NADPH, and 500 µM DBCO-acid in exchange buffer (20 mM HEPES pH 8.0, 10 mM MgCl_2_, 50 mM NaCl, 10% glycerol) were incubated aerobically at room temperature for 16 h. Reaction mixtures were diluted with 70 µL of exchange buffer, 15 µL of MeOH, and 15 µL of ACN and filtered (3 kDa, Pall, Omega membrane).

Detection of DOBA-DBCO and DAC-DBCO was accomplished using a Thermo Orbitrap IQ-X Tribrid mass spectrometer as described above with minor modifications. The MS settings were: mass range 70-700 *m/z*. MS/MS spectra of DAC-DBCO were obtained using HCD fragmentation with 30% normalized collision energy. MS/MS spectra of DOBA-DBCO were obtained using CID fragmentation with 30% collision energy, 10 ms activation time, and 0.25 activation Q.

### ICP-MS of Dob3

60 µL of trace-metal free nitric acid was added to 200 µL of 1 mg/mL Dob3 in water. This mixture was incubated for 3 h at 60 °C before precipitates were removed by centrifugation. 200 µL of supernatant was diluted to a total volume of 3 mL with molecular biology grade water. Samples were analyzed on an Agilent 7900 Inductive Coupled Mass Spectrometer.

### Ferrozine assay of Dob3

12.5 µL of 50% aqueous TFA was added to 62.5 µL of 186 µM Dob3, mixed, and incubated for 5 min at room temp. 125 µM, 250 µM, 500 µM, and 1000 µM standards of ferrous ammonium sulfate were prepared analogously and blanks were prepared using exchange buffer (20 mM HEPES pH 8.0, 10 mM MgCl_2_, 50 mM NaCl, 10% glycerol). The sample was spun down at max speed for 2 min. 50 µL of supernatant was transferred to a new Eppendorf and 450 µL of H_2_O, 20 µL of 75 mM ascorbic acid, 20 µL of 10 mM ferrozine, and 120 µL of saturated ammonium acetate were added. Absorbance at 562 nm was measured on an Agilent Cary 3500 UV-Vis spectrophotometer. The average of the blank absorbances was subtracted from sample and standard measurements before further analysis.

### Substrate scope of Dob3

50 µL reaction mixtures containing 1 mM of the specified hydrazone substrate, 2 mM NADPH, 100 µM PMS, 20 µM Dob3, and 500 µM DBCO-acid in exchange buffer (20 mM HEPES pH 8.0, 10 mM MgCl_2_, 50 mM NaCl, 10% glycerol) were incubated overnight. Samples were diluted with 70 µL of exchange buffer, 15 µL of MeOH, and 15 µL of ACN and filtered (3 kDa, VWR, nylon). Detection of corresponding pyrazoles was accomplished using a Thermo Orbitrap IQ-X Tribrid mass spectrometer as described above. HCD fragmentation with 30% normalized collision. Spectra were acquired in positive ion mode. A mass window of 5 ppm was used for extracted ion chromatograms.

### One pot chemoenzymatic production of ethyl diazoacetate

50 µL reaction mixtures containing 100 µM PMS, 20 µM Dob3, 1 mM ethyl glyoxylate, 2 mM NADPH, 2 mM hydrazine hydrate, and 500 µM DBCO-acid in exchange buffer (20 mM HEPES pH 8.0, 10 mM MgCl_2_, 50 mM NaCl, 10% glycerol) were incubated overnight. 70 µL of buffer, 15 µL of ACN, and 15 µL of MeOH were added and the reactions were centrifugal filtered (3 kDa, Pall, Omega membrane). Detection of EDA-DBCO was accomplished using a Thermo Orbitrap IQ-X Tribrid mass spectrometer as described above with minor modifications. The MS settings were: mass range 70-700 *m/z*.

### One pot chemoenzymatic production of diazomethylbenzene

50 µL reaction mixtures containing 100 µM PMS, 20 µM Dob3, 2 mM benzaldehyde, 2 mM NADPH, 2 mM hydrazine hydrate, and 500 µM DBCO-acid in exchange buffer (20 mM HEPES pH 8.0, 10 mM MgCl_2_, 50 mM NaCl, 10% glycerol) were incubated overnight. 70 µL of buffer, 15 µL of ACN, and 15 µL of MeOH were added and the reaction mixtures were filtered (3 kDa, Pall, Omega membrane). Detection of EDA-DBCO was accomplished using a Thermo Orbitrap IQ-X Tribrid mass spectrometer as described above with minor modifications. The MS settings were: mass range 70-700 *m/z*.

### Anaerobic/aerobic hydrazone incubation

50 µL reaction mixtures containing 500 µM benzylidenehydrazine, 100 µM PMS, 2 mM NADPH, and 500 µM DBCO-acid in exchange buffer (20 mM HEPES pH 8.0, 10 mM MgCl_2_, 50 mM NaCl, 10% glycerol) was incubated for 22 h at room temperature either aerobically or anaerobically. Samples were quenched aerobically or anaerobically through addition of 70 µL of buffer, 15 µL of LC–MS ACN, and 15 µL of LC–MS MeOH and then filtered (0.2 µm, VWR, nylon). Anaerobic samples were moved to the autosampler approximately 5 min prior to injection. Detection of diazomethylbenzene-DBCO was accomplished using a Thermo Orbitrap IQ-X Tribrid mass spectrometer as described above with minor modifications. The MS settings were: mass range 70-700 *m/z*.

### Synthesis of 6-(3-acetyl-1,9-dihydro-8*H*-dibenzo[*b*,*f*]pyrazolo[4,3-*d*]azocin-8-yl)-6-oxohexanoic acid (DAC-DBCO regioisomers)

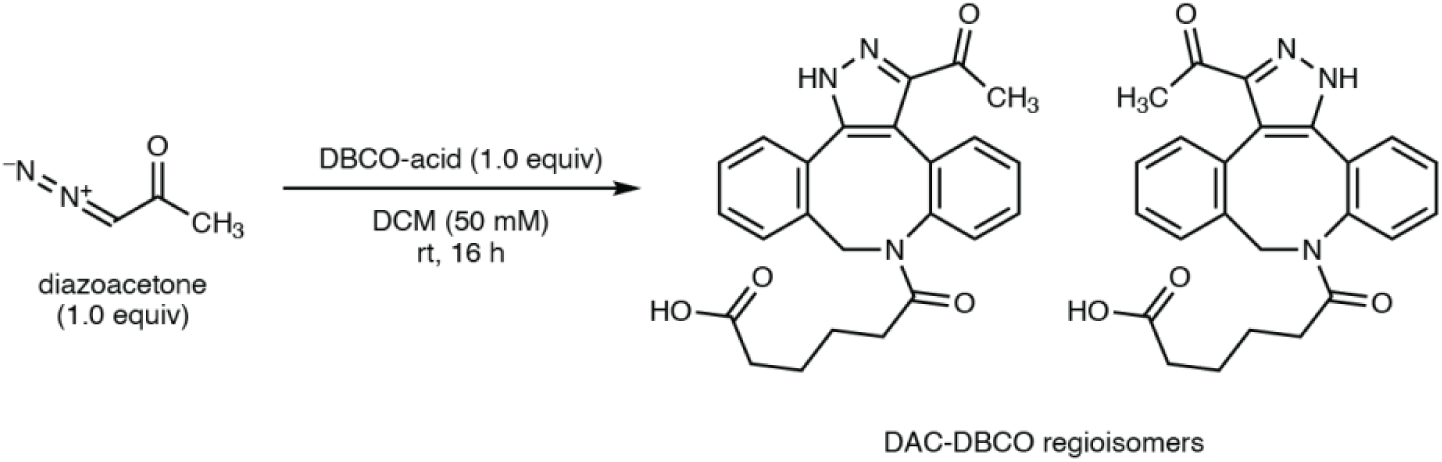

39.7 mg of DBCO-acid (0.12 mmol, 1.0 equiv) was added to 100 mg of diazoacetone (0.12 mmol, 1 equiv) in 2.4 mL of DCM in a capped vial. The reaction mixture was stirred overnight, and 0.5 mL of DCM was added the next morning to replace evaporated solvent. The mixture was concentrated under vacuum and purified using reversed phase chromatography on a Biotage Selekt using a 6 g C18 column with a 2% to 95% gradient of ACN in H_2_O. Fractions were concentrated under reduced pressure and then lyophilized. The product was re-purified using a Dionex UltiMate HPG-3200Bx Semi-Preparative HPLC using a HypersilGold column with a 5% to 95% ACN (+ 0.1% formic acid) in H_2_O (+ 0.1% formic acid) gradient. The fractions were concentrated under reduced pressure and lyophilized. NMR spectra were obtained using a Bruker AVANCE NEO 400B spectrometer (400 MHz, 100 MHz) and correspond to a 1:1 mixture of both pyrazole regioisomers. HRMS data were collected using a Thermo Orbitrap IQ-X Tribrid mass spectrometer coupled to a Horizon Vanquish UHPLC as described above.

**^1^H NMR:** (400 MHz, CDCl_3_) δ 7.50 – 7.32 (m, 7H), 7.30 – 7.27 (m, 2H), 7.25 – 7.17 (m, 11H), 7.07 – 6.99 (m, 2H), 5.92 (d, *J* = 16.9 Hz, 1H), 5.77 (d, *J* = 15.5 Hz, 1H), 4.42 (d, *J* = 15.5 Hz, 1H), 4.26 (d, *J* = 17.0 Hz, 1H), 2.64 (s, 3H), 2.31 (s, 3H), 2.26 – 2.12 (m, 3H), 1.84 (m, 3H), 1.71 (m 1H), 1.53 – 1.22 (m, 4H).

**^13^C NMR:** (101 MHz, CDCl_3_) δ 194.50, 190.58, 178.09, 177.44, 174.14, 173.27, 148.89, 145.76, 143.20, 140.45, 140.07, 139.78, 134.78, 134.49, 133.34, 132.52, 132.11, 131.99, 131.06, 130.76, 130.49, 129.84, 129.57, 129.27, 129.20, 128.63, 128.57, 128.46, 127.59, 127.11, 126.78, 126.72, 123.35, 117.42, 52.70, 52.26, 33.88, 33.66, 33.52, 28.63, 27.76, 24.27, 24.20, 23.93.

**HRMS** (ESI+, *m/z*): [M + H]^+^ calculated for C_24_H_24_N_3_O_4_^+^, 418.1761; found 418.1760.

### Synthesis of methyl 4-diazo-3-oxobutanoate (OMe-DOBA)

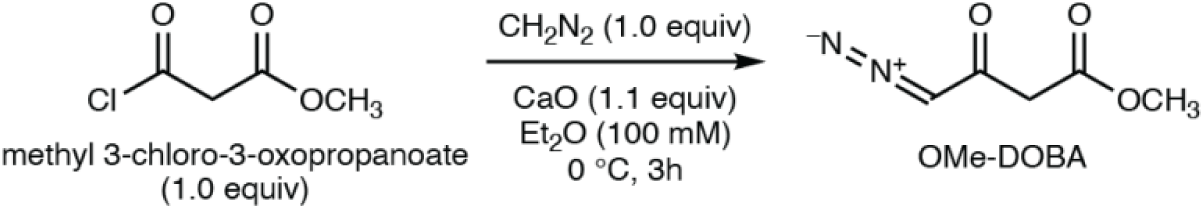

A freshly prepared and titrated solution of diazomethane in diethyl ether (0.25 M, 4.0 mL, 1.0 mmol, 1.0 equiv)^98^ was added to a suspension of calcium oxide (61.7 mg, 1.0 mmol, 1.1 equiv) in anhydrous diethyl ether (5.0 mL) at 0 °C.^99^ The suspension was stirred at 0 °C for 5 minutes, and methyl 3-chloro-3-oxopropanoate (0.107 mL, 1.0 mmol, 1.0 equiv) in anhydrous diethyl ether (0.5 mL) was added dropwise. The reaction mixture was stirred at 0 °C for 3 hours, allowed to warm to room temperature, filtered, and concentrated under reduced pressure (ambient water bath) to yield a crude light-yellow oil. Purification by flash-column chromatography (30 – 50% ethyl acetate in hexanes) afforded methyl 4-diazo-3-oxobutanoate (85.0 mg, 0.598 mmol, 60% yield) as a light-yellow oil.

**^1^H NMR:** (400 MHz, CDCl_3_) δ 5.53 (s, 1H), 3.74 (s, 3H), 3.36 (s, 2H).

**^13^C NMR:** (101 MHz, CDCl_3_) δ 186.07, 167.85, 56.12, 52.68, 46.87.

**FTIR** (neat), cm^-1^: 3102 (m), 2956 (w), 2107 (s), 1739 (s), 1636 (s).

**HRMS** (ESI+, *m/z*): [M+H–N_2_]^+^ calculated for C_5_H_7_O_3_^+^, 115.0390; found 115.0387.

### Synthesis of methyl (*E*)-4-hydrazono-3-oxobutanoate (OMe-HOBA)

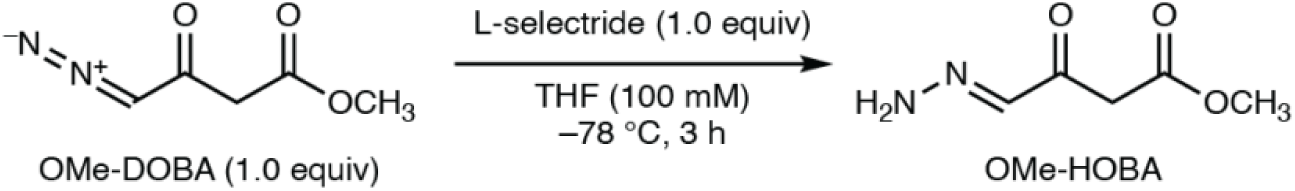

Methyl 4-diazo-3-oxobutanoate (OMe-DOBA) (15.0 mg, 106 µmol, 1.0 equiv) was dissolved in THF (1.06 mL) and cooled to –78 °C. L-selectride in THF (1.0 M, 106 µL, 1.0 equiv) was added dropwise and the resultant canary-yellow solution was stirred at the same temperature for 3 hours.^100^ The reaction mixture was quenched at –78 °C with methanol (42.7 µL, 10 equiv), warmed to room temperature, and concentrated under reduced pressure. Purification by flash-column chromatography (0 – 5% methanol in dichloromethane) afforded methyl (*E*)-4-hydrazineylidene-3-oxobutanoate (OMe-HOBA) (5.0 mg, 34.7 µmol, 33% yield) as a light-yellow film.

**^1^H NMR:** (400 MHz, CD_2_Cl_2_) δ 7.09 (s, 1H), 6.49 (s, 2H), 3.72 (s, 2H), 3.68 (s, 3H).

**^13^C NMR:** (101 MHz, CD_2_Cl_2_) δ 192.14, 169.02, 137.67, 52.57, 43.99.

**FTIR** (neat), cm^-1^: 3430 (m), 3308 (m), 3225 (m), 2954 (m), 1739 (s), 1664 (s), 1596 (s).

**HRMS** (ESI+, *m/z*): [M+H]^+^ calculated for C_5_H_9_N_2_O_3_^+^, 145.0608; found 145.0609.

NB: When dissolved in CDCl_3_, the (*E*)-hydrazone was observed to isomerize to the (*Z*)-hydrazone.

### Synthesis of *N*-6-hydroxylysine

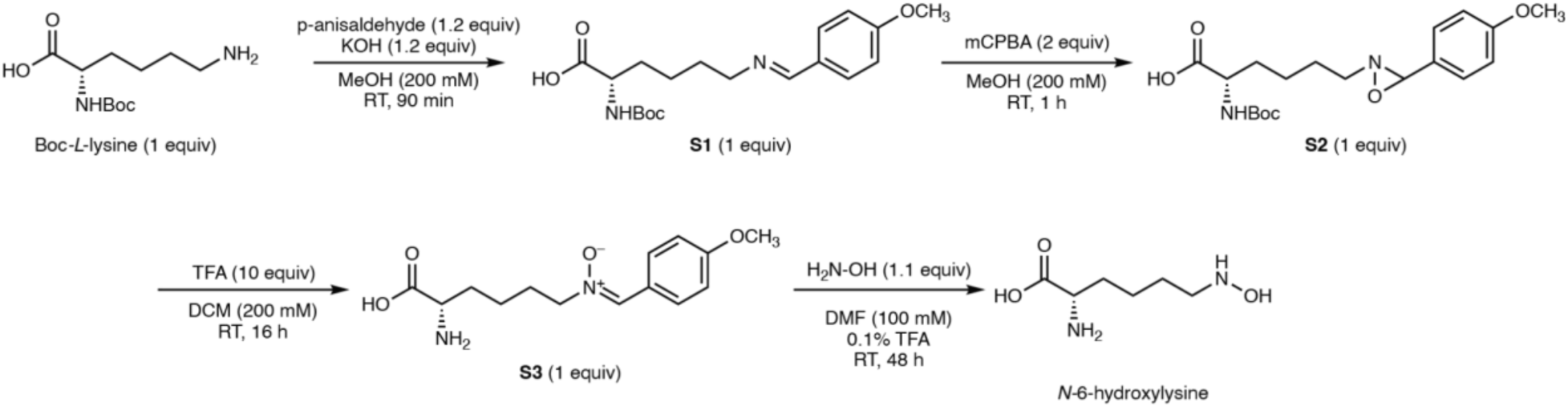

*N*-6-hydroxylysine was synthesized according to a previously reported procedure.^31^ NMR and HRMS data matched the previously reported values.

### Synthesis of HAA

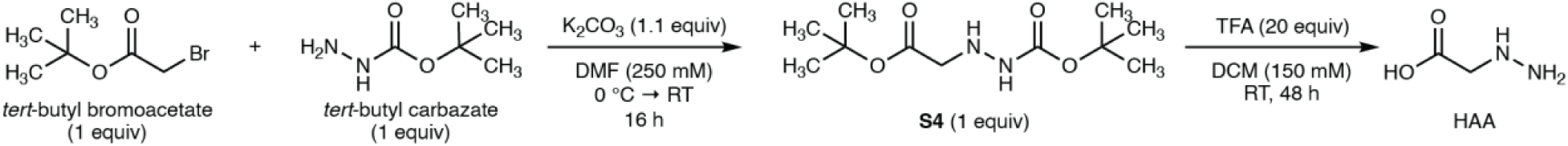

Hydrazinoacetic acid was synthesized according to a previously reported procedure.^31^ NMR and HRMS data matched previously reported values.

### Synthesis of 6-(3-(3-methoxy-3-oxopropanoyl)-1,9-dihydro-8*H*-dibenzo[*b*,*f*]pyrazolo[4,3-*d*]azocin-8-yl)-6-oxohexanoic acid & 6-(3-(3-methoxy-3-oxopropanoyl)-1,8-dihydro-9*H*-dibenzo[*b*,*f*]pyrazolo[3,4-*d*]azocin-9-yl)-6-oxohexanoic acid (OMe-DOBA-DBCO regioisomers)

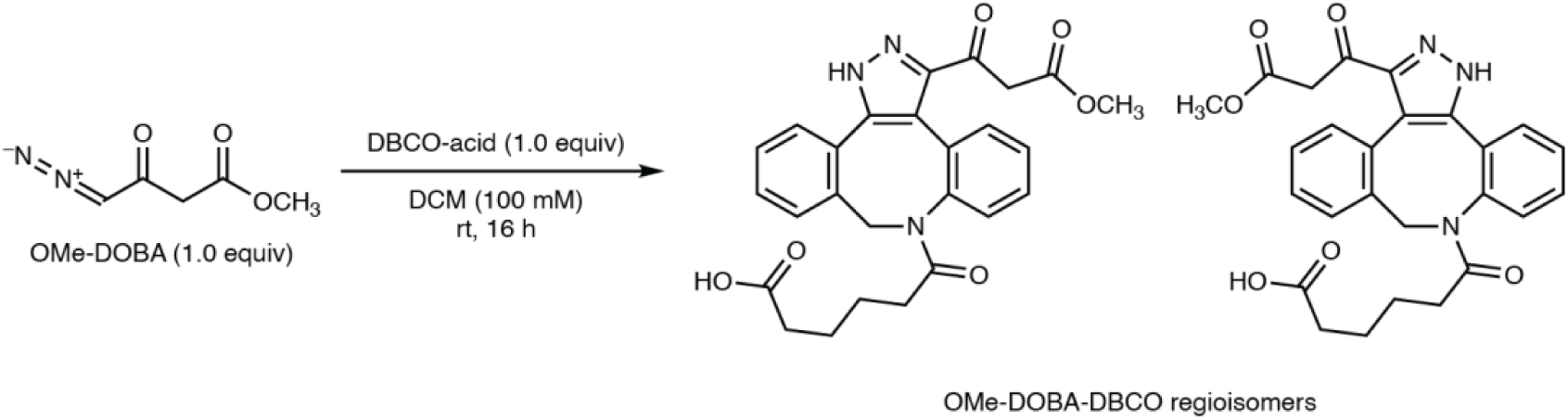

DBCO-C_6_-Acid (14.00 mg, 42.0 μmol, 1.0 equiv) was dissolved in dichloromethane (420 μL) and methyl 4-diazo-3-oxobutanoate (5.97 mg, 42.0 μmol, 1.0 equiv) dissolved in dichloromethane (420 μL) was added. The colorless solution was stirred for 16 hours, concentrated under reduced pressure, and purified by preparatory-HPLC (5 – 50% acetonitrile in water, with 0.1% HCOOH additive throughout, over 30 minutes) to yield an inseparable mixture of regioisomeric OMe-DOBA-DBCO-C_6_-acids in approximately a 2:1 ratio (16.7 mg, 35.1 μmol, 84%) as a fluffy white powder after lyophilization.

**^1^H NMR:** (400 MHz, CDCl_3_) δ 7.53 – 7.47 (m, 1H), 7.47 – 7.32 (m, 4H), 7.30 – 7.15 (m, 5H), 7.13 – 7.08 (m, 1H), 6.94 – 6.88 (m, 1H), 5.82 (d, *J* = 17.0 Hz, 1H), 5.62 (d, *J* = 15.3 Hz, 0.5H), 4.44 (d, *J* = 15.4 Hz, 0.5H), 4.39 (d, *J* = 16.3 Hz, 1H), 4.19 (d, *J* = 17.1 Hz, 1H), 4.00 (d, *J* = 16.2 Hz, 1H), 3.84 (d, *J* = 9.5 Hz, 0.5H), 3.77 (d, *J* = 8.4 Hz, 0.5H), 3.74 (s, 3H), 3.70 (s, 1.5H), 2.31 – 2.11 (m, 3H), 1.93 – 1.79 (m, 3H), 1.76 – 1.69 1.72 (m, 1H), 1.58 – 1.20 (m, 7H).

**^13^C NMR:** (101 MHz, CDCl_3_) δ 189.50, 185.77, 178.19, 177.24, 174.49, 173.03, 169.21, 167.78, 147.31, 146.04, 142.42, 141.18, 140.57, 139.65, 134.56, 134.33, 132.86, 132.20, 132.16, 132.08, 131.45, 131.10, 130.82, 130.65, 129.83, 129.62, 129.16, 129.11, 128.92, 128.63, 127.71, 126.91, 126.55, 126.13, 123.44, 117.25, 52.92, 52.64, 52.46, 52.18, 46.55, 45.97, 33.88, 33.49, 33.43, 24.36, 24.27, 24.23, 23.90.

**HRMS** (ESI+, *m/z*): [M + H]^+^ calculated for C_26_H_26_N_3_O_6_^+^, 476.1816; found 476.1824.

### Synthesis of Ethyl (*E*)-2-hydrazineylideneacetate

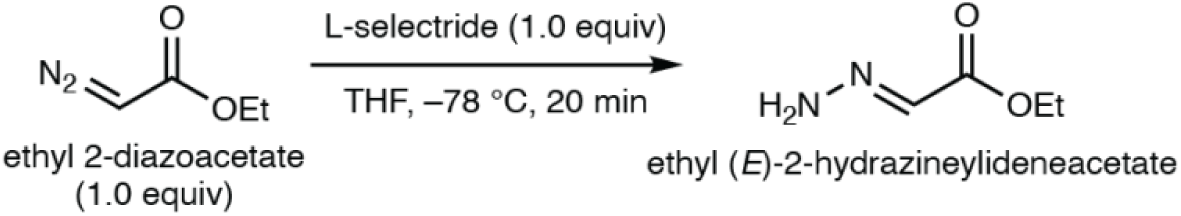

To a yellow solution of ethyl 2-diazoacetate (87 Wt% in dichloromethane, 100 μL, 827 μmol, 1.0 equiv.) dissolved in THF (4.14 mL) at –78 °C was added 1.0 M L-selectride in THF (827 μL, 827 μmol, 1.0 equiv.) dropwise. The reaction was stirred at the same temperature for 20 minutes and quenched at –78 °C by the addition of a minimal amount of ethanol. The solution was allowed to warm up to room temperature and concentrated to yield a crude blood-orange oil. The crude oil was purified by flash column chromatography (50% ethyl acetate in hexanes) to yield ethyl (*E*)-2-hydrazineylideneacetate (48.0 mg, 413 μmol, 50%) as a colorless film.

**^1^H NMR:** (400 MHz, CD_2_Cl_2_) δ 7.02 (s, 1H), 6.38 (s, 2H), 4.21 (q, *J* = 7.1 Hz, 2H), 1.29 (t, *J* = 7.2 Hz, 3H).

**^13^C NMR**: (101 MHz, CD_2_Cl_2_) δ 164.46, 130.02, 61.12, 14.60.

**HRMS** (ESI+, *m/z*): [M+H]^+^ calculated for C_4_H_9_N_2_O_2_^+^, 117.0659; found 117.0657.

### Synthesis of (*E*)-2-hydrazineylidene-*N*,*N*-dimethylacetamide

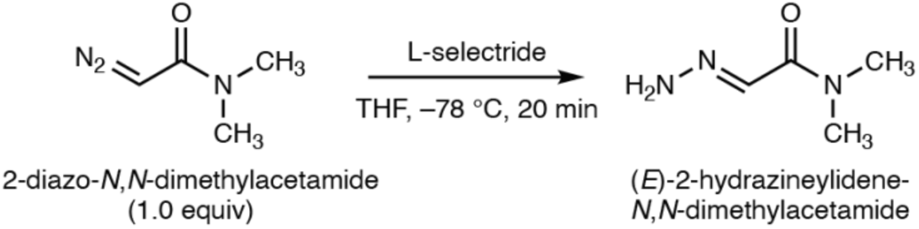

To a yellow solution of 2-diazo-*N,N*-dimethylacetamide^101^ (68 mg, 601 μmol, 1 equiv.) dissolved in THF (3.0 mL) at –78 °C was added 1.0 M L-selectride in THF (601 μL, 601 μmol, 1.0 equiv.) dropwise. The reaction was stirred at the same temperature for 20 minutes and quenched at – 78 °C by the addition of a minimal amount of ethanol. The solution was allowed to warm up to room temperature and concentrated to yield a crude orange oil. The crude oil was purified by flash column chromatography (0 – 5% methanol in dichloromethane) to yield (*E*)-2-hydrazineylidene-*N*,*N*-dimethylacetamide (38.2 mg, 332 μmol, 55%) as a white film.

**^1^H NMR:** (400 MHz, CD_2_Cl_2_) δ 7.33 (s, 1H), 5.94 (s, 2H), 3.14 (s, 3H), 2.95 (s, 3H).

**^13^C NMR:** (101 MHz, CD_2_Cl_2_) δ 164.56, 164.37, 134.46, 120.74, 38.11, 37.54, 36.20, 35.08.

NB: The (*E*)-hydrazone was observed to isomerize (∼50% conversion) to the (*Z*)-hydrazone over the timespan of the ^13^C NMR experiment in CD_2_Cl_2_. Therefore, eight ^13^C peaks, rather than the expected four, were observed.

**HRMS** (ESI+, *m/z*): [M+H]^+^ calculated for C_4_H_10_N_3_O^+^, 116.0818; found 116.0818.

### Synthesis of (*S*)-6-(3-((2-amino-2-carboxyethoxy)carbonyl)-1,9-dihydro-8*H*-dibenzo[*b*,*f*]pyrazolo[4,3-*d*]azocin-8-yl)-6-oxohexanoic acid & (*S*)-6-(3-((2-amino-2-carboxyethoxy)carbonyl)-1,8-dihydro-9*H*-dibenzo[*b*,*f*]pyrazolo[3,4-*d*]azocin-9-yl)-6-oxohexanoic acid (Azaserine-DBCO regioisomers)

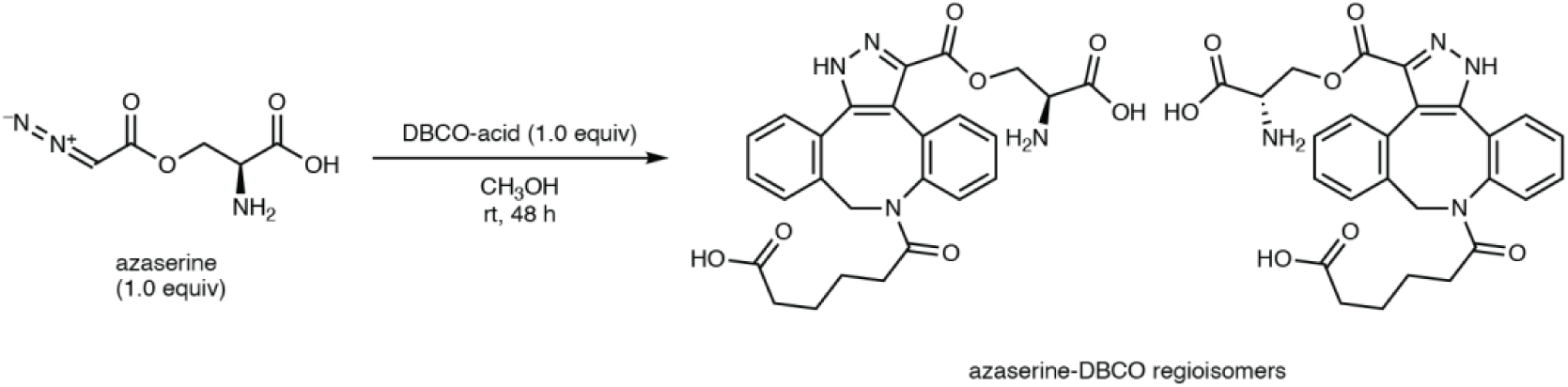

A suspension of azaserine (5.85 mg, 33.8 μmol, 1.0 equiv.) and DBCO-acid (11.3 mg, 33.8 μmol, 1.0 equiv.) in methanol (1.35 mL) was stirred for 48 hours, upon which the suspension dissolved completely to yield a colorless solution. The reaction mixture was directly purified by preparatory-HPLC (5 – 50% acetonitrile in water, with 0.1% HCOOH additive throughout, over 30 minutes) to yield an inseparable mixture of regioisomeric Azaserine-DBCO-C_6_-acids (5.0 mg, 9.87 μmol, 29%) as a fluffy white powder after lyophilization.

**^1^H NMR:** (400 MHz, CD_3_OD) δ 7.73 – 6.95 (m, 8H), 5.98 (d, *J* = 16.7 Hz, 0.6H), 5.66 (dd, *J* = 15.4, 9.9 Hz, 0.4H), 4.79 – 4.63 (m, 1.6H), 4.58 – 4.38 (m, 1.4H), 4.08 – 3.89 (m, 1H), 2.23 – 2.04 (m, 2H), 1.95 – 1.72 (m, 1.6H), 1.65 (dd, *J* = 15.8, 7.4 Hz, 0.4H), 1.45 – 1.25 (m, 4H).

**^13^C NMR:** (101 MHz, CD_3_OD) δ 177.84, 177.55, 175.12, 174.63, 170.81, 170.53, 141.71, 141.23, 141.19, 136.15, 134.81, 134.51, 134.41, 133.24, 133.22, 132.95, 132.77, 132.05, 130.91, 130.88, 130.66, 130.56, 130.52, 130.38, 129.84, 129.62, 129.58, 129.06, 128.51, 128.45, 128.41, 128.24, 128.16, 65.03, 64.78, 55.30, 55.19, 53.40, 52.87, 34.87, 34.85, 34.74, 34.70, 34.66, 25.59, 25.54, 25.51, 25.47, 25.40.

**HRMS** (ESI+, *m/z*): [M+H]^+^ calculated for C_26_H_27_N_4_O_7_^+^, 507.1874; found 507.1876

### Synthesis of 6-oxo-6-(3-phenyl-1,9-dihydro-8*H*-dibenzo[*b*,*f*]pyrazolo[4,3-*d*]azocin-8-yl)hexanoic acid & 6-oxo-6-(3-phenyl-1,8-dihydro-9*H*-dibenzo[*b*,*f*]pyrazolo[3,4-*d*]azocin-9-yl)hexanoic acid (Diazomethylbenzene-DBCO regioisomers)

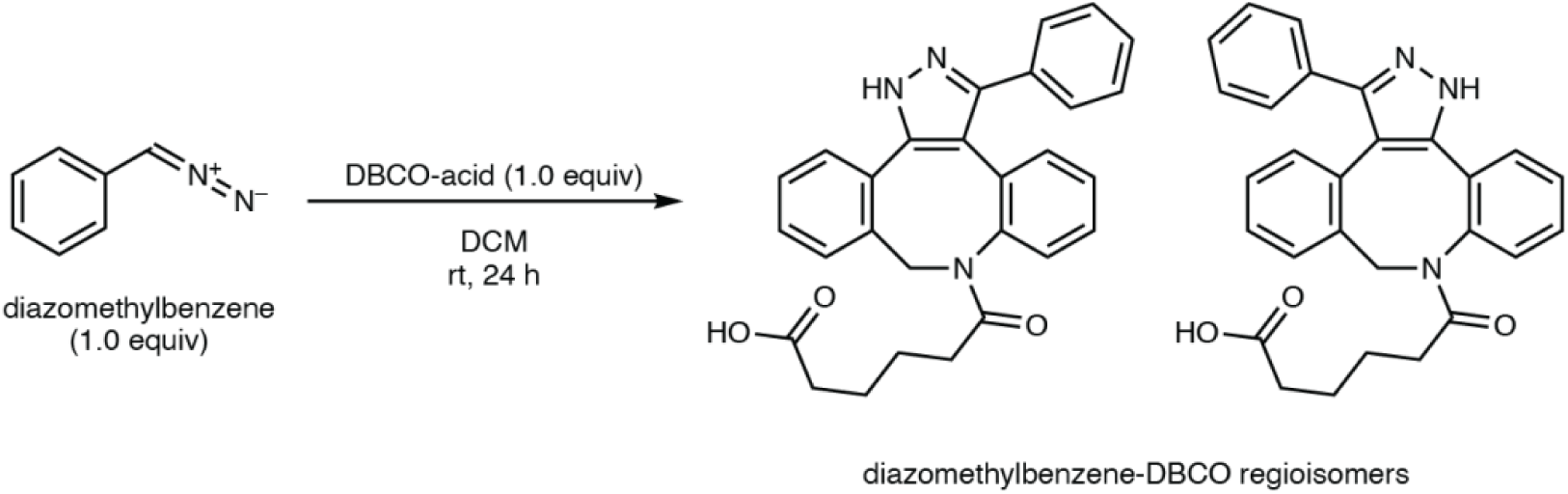

A peach-colored solution of diazomethylbenzene (19.7 mg, 167 μmol, 1.33 equiv.) in dichloromethane-diethyl ether-pentane^102^ (1:9:10 ratio, 3.34 mL) was added to DBCO-acid (41.6 mg, 125 μmol, 1.0 equiv.) in dichloromethane (333 μL) and stirred for 24 hours. The reaction mixture was concentrated and purified by flash column chromatography (0 – 5% methanol in dichloromethane) to yield an inseparable mixture of regioisomeric Diazomethylbenzene-DBCO-C_6_-acids (12.0 mg, 26.6 μmol, 16%) as a fluffy white powder after lyophilization.

**^1^H NMR:** (400 MHz, CDCl_3_) δ 7.58 – 7.27 (m, 8.6H), 7.25 – 7.07 (m, 3.4H), 7.00 (dtd, *J* = 15.1, 7.5, 1.4 Hz, 0.6H), 6.79 (dd, *J* = 7.7, 1.4 Hz, 0.4H), 6.00 (d, *J* = 16.3 Hz, 0.6H), 5.93 (d, *J* = 15.6 Hz, 0.4H), 4.51 – 4.35 (m, 1H), 2.17 – 1.96 (m, 2.6H), 1.91 – 1.78 (m, 1H), 1.54 – 1.17 (m, 4.4H).

**^13^C NMR:** (101 MHz, CDCl_3_) δ 177.69, 177.18, 173.12, 172.80, 147.85, 147.06, 143.18, 142.73, 140.94, 140.71, 134.75, 134.65, 133.51, 133.14, 132.86, 131.95, 131.90, 131.50, 130.44, 130.25, 130.12, 129.64, 129.42, 129.38, 129.33, 129.19, 128.94, 128.89, 128.81, 128.76, 128.72, 128.50, 128.40, 127.90, 127.50, 127.47, 127.11, 118.21, 115.16, 53.03, 52.88, 34.14, 33.94, 33.92, 33.75, 24.51, 24.40, 24.26.

**HRMS** (ESI+, *m/z*): [M+H]^+^ calculated for C_28_H_26_N_3_O_3_^+^, 452.1969; found 452.1974.

## Data Availability

Raw LC–MS data and LC–MS/MS data are available upon request due to the large file size. Previously published crystal structures are available in the Protein Data Bank (https://www.rcsb.org/) under accession codes 3CHH and 5HYH. All other data are available in the manuscript or Supplementary Information. Source data are provided with this paper.

## Acknowledgements

We thank Dr. Grace Kenney for her assistance with genome mining. We thank Terra Bioforge for construction of *dob*-pDualP and *dob*-pDualP ΔDob3. D.V.C. acknowledges funding from the National Science Foundation Graduate Research Fellowship Program (grant numbers DGE2140743 and DGE1745303). We acknowledge financial support from the National Institute of Health (grant number 5R01GM132564-04). E.P.B is a Howard Hughes Medical Institute Investigator. This article is subject to HHMI’s Open Access to Publications policy. HHMI lab heads have previously granted a nonexclusive CC BY 4.0 license to the public and a sublicensable license to HHMI in their research articles. Pursuant to those licenses, the author-accepted manuscript of this article can be made freely available under a CC BY 4.0 license immediately upon publication.

## Author Contributions

D.V.C. and E.P.B. initiated the study. K.P. and D.V.C. designed and optimized the reactivity-based screen. K.P. and D.V.C. performed LC–MS metabolomics and identified DAC and DOBA. K.P. and D.V.C. performed bioinformatics analyses and identified the gene cluster. K.P. and D.V.C. performed heterologous expression of the biosynthetic gene cluster. K.P. and D.V.C. performed biochemical characterization of DobA, DobB, DobC, DobE, DobF, DobG, DobM, DobQ, Dob2, and Dob3. K.P. performed the in vivo gene knockout. K.P. and D.V.C. designed substrate scope and chemoenzymatic experiments. K.P. performed substrate scope and chemoenzymatic experiments. K.W. performed chemical syntheses of substrates and standards. All authors analyzed and discussed the results and prepared the manuscript.

## Extended Data Figures and Tables

**Extended Data Figure 1:**
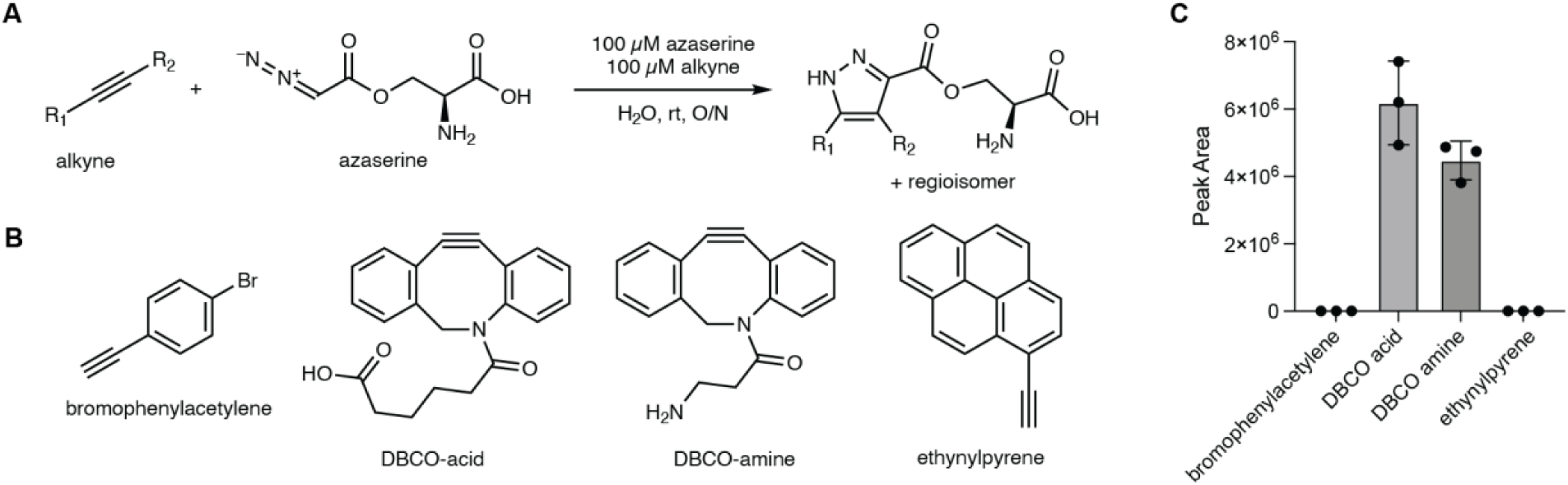
Azaserine undergoes cycloadditions with strained cyclooctynes, but not terminal alkynes. **A)** Reaction scheme of azaserine and alkynes to yield pyrazole regioisomers. **B)** Bromophenylacetylene, DBCO-acid, DBCO-amine, and ethynylpyrene were incubated with azaserine. **C)** Peak areas resulting from the cycloadditions. Error bars indicate mean ± standard deviation. Peak areas are calculated as the sum of the two regioisomer peaks. Experiments were run in biological triplicate.

**Extended Data Figure 2:**
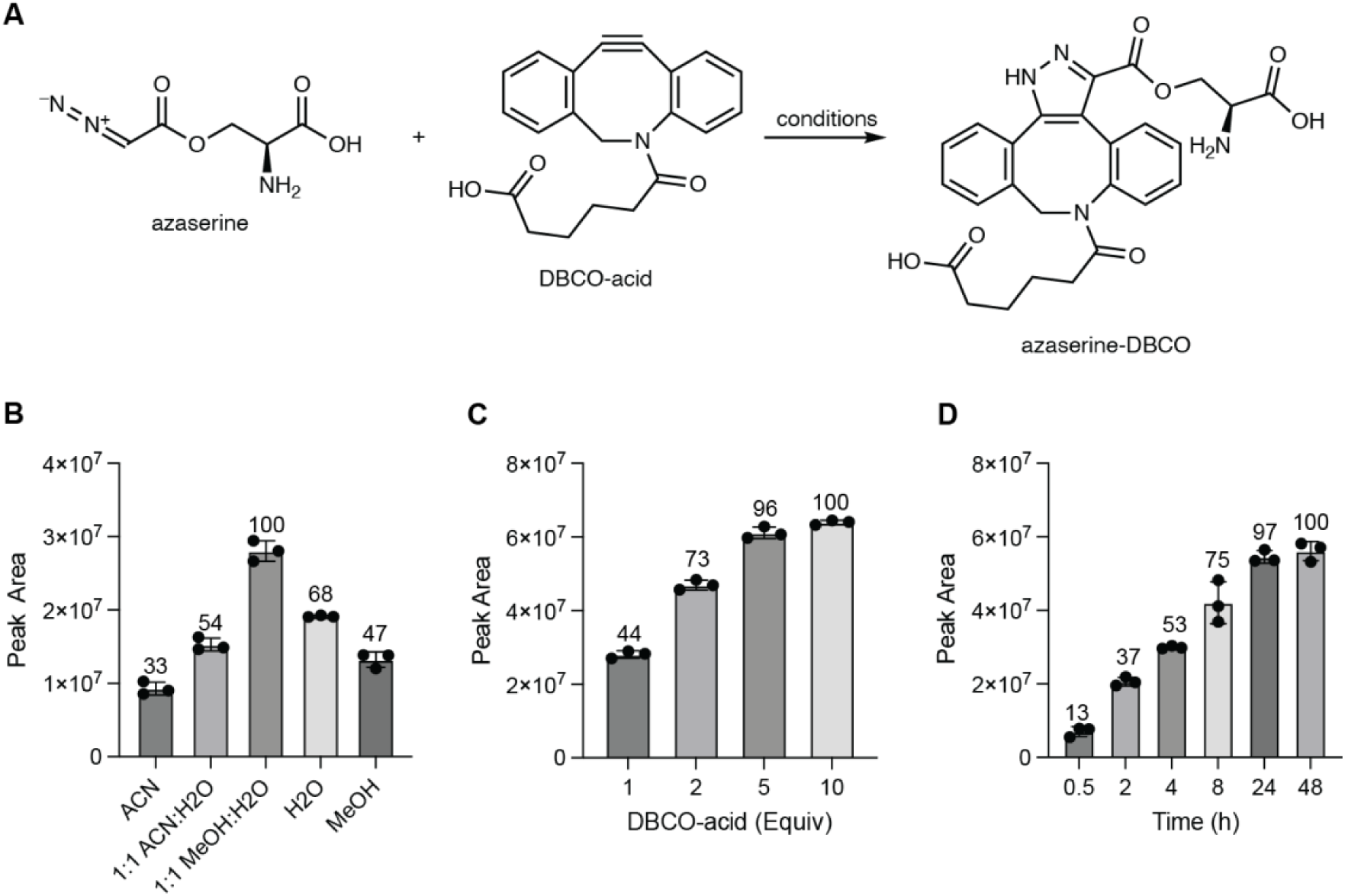
Optimization of the model reaction of azaserine with DBCO-acid. **A)** model reaction between azaserine and DBCO-acid. **B)** Solvent optimization of the model reaction. **C)** Optimization of DBCO-acid equivalents for the model reaction. **D)** Time course of the model reaction. For all panels, values are normalized to the maximum. Peak areas are calculated as the sum of the two regioisomers. Error bars indicate mean ± standard deviation. Experiments were run in biological triplicate.

**Extended Data Figure 3:**
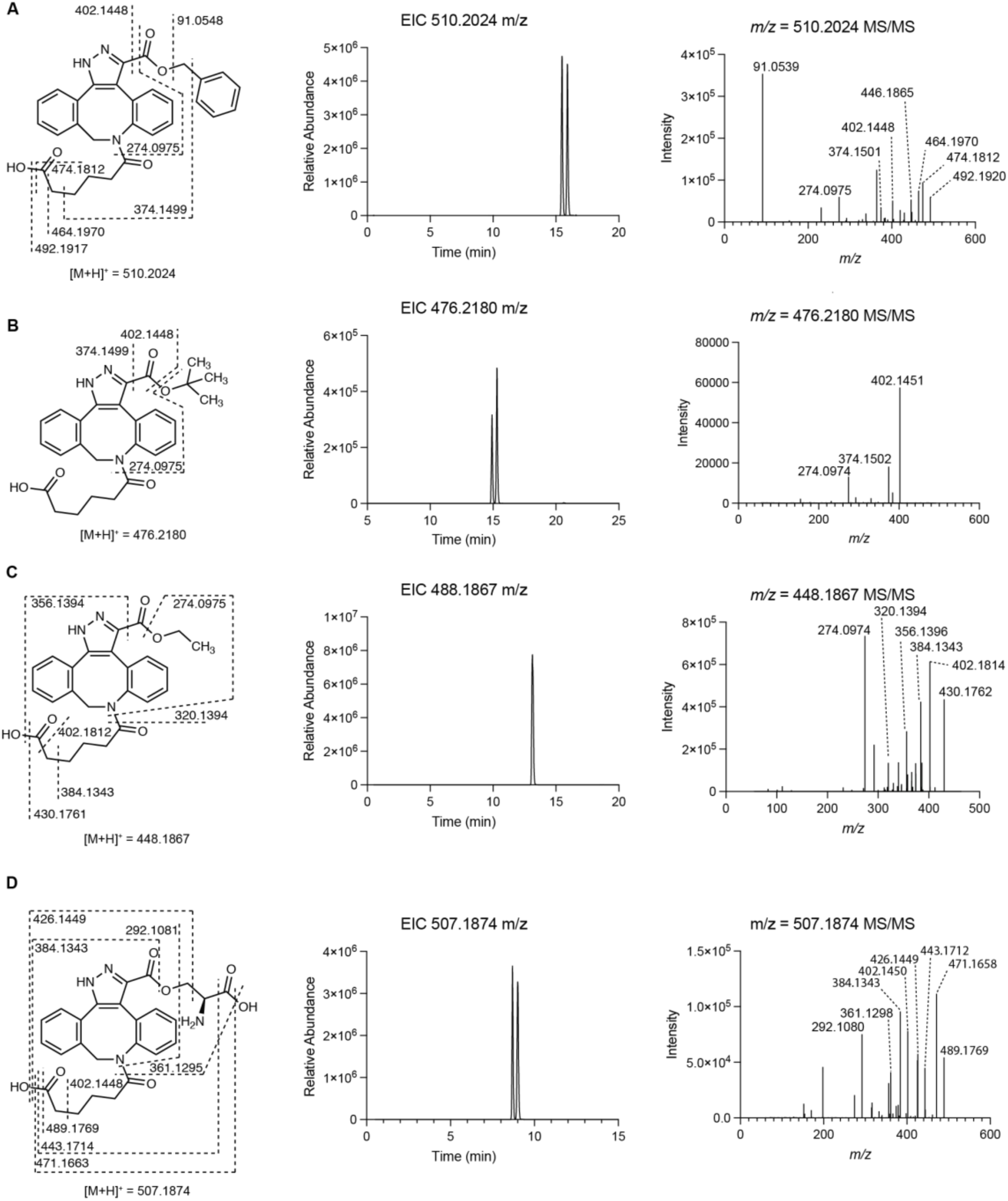
DBCO-acid substrate scope. **A**) EIC and targeted MS/MS of benzyl diazo acetate-DBCO (*m/z* = 510.2024 ± 5 ppm). Experiments were run in biological triplicate. Representative results are shown. **B**) EIC and targeted MS/MS of *tert*-butyl diazoacetate-DBCO (*m/z* = 476.2180 ± 5 ppm) **C**) EIC and targeted MS/MS of ethyl diazo acetate-DBCO (*m/z* = 448.1867 ± 5 ppm). We hypothesize only one peak is observed in the EIC due to the two regioisomers having identical retention times or the formation of one regioisomer being strongly favored. **D**) EIC and targeted MS/MS of azaserine-DBCO (*m/z* = 507.1874 ± 5 ppm). For all panels, experiments were run in biological triplicate. Representative results are shown.

**Extended Data Figure 4:**
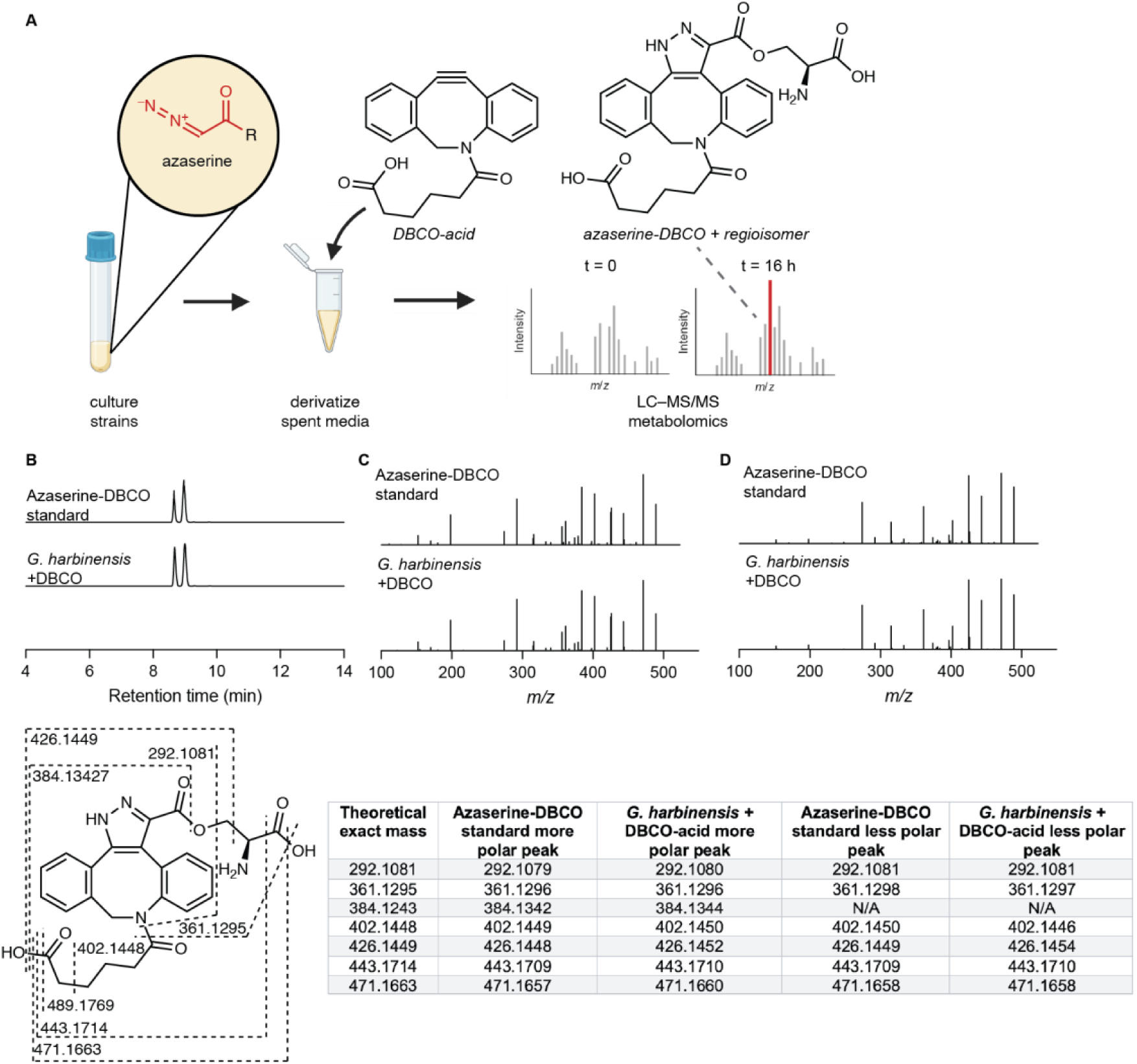
DBCO-acid reacts selectively with diazo-containing natural products in complex biological matrices. **A)** Reaction of DBCO-acid with azaserine in a complex metabolite matrix. DBCO-acid was incubated for 16 h with *G. harbinensis* spent medium. **B)** EICs of azaserine-DBCO-acid standard and azaserine-DBCO-acid (*m/z* = 507.1874 ± 5 ppm) produced by *G. harbinensis* spent media derivatized with DBCO-acid. Experiments were run in biological triplicate. Representative results are shown. **C)** MS/MS fragmentation of the more polar (earlier eluting) regioisomer of the azaserine-DBCO synthetic standard compared to *G. harbinensis* spent medium treated with DBCO-acid. **D**) MS/MS fragmentation of the less polar (later eluting) regioisomer of the azaserine-DBCO synthetic standard compared to *G. harbinensis* spent medium treated with DBCO-acid.

**Extended Data Figure 5:**
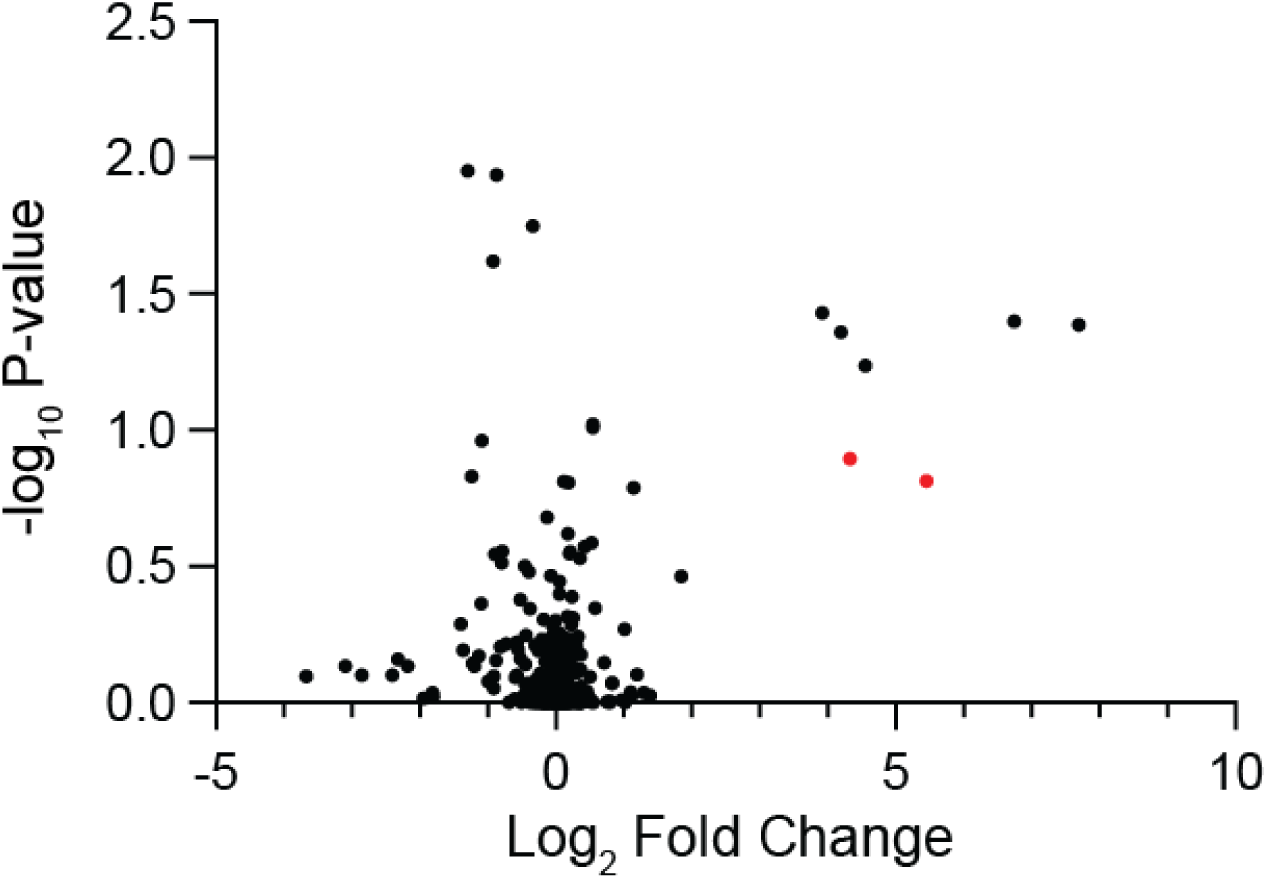
Volcano plot of comparative metabolomics for *G. harbinensis* spent media. Azaserine-DBCO regioisomers are highlighted in red.

**Extended Data Figure 6:**
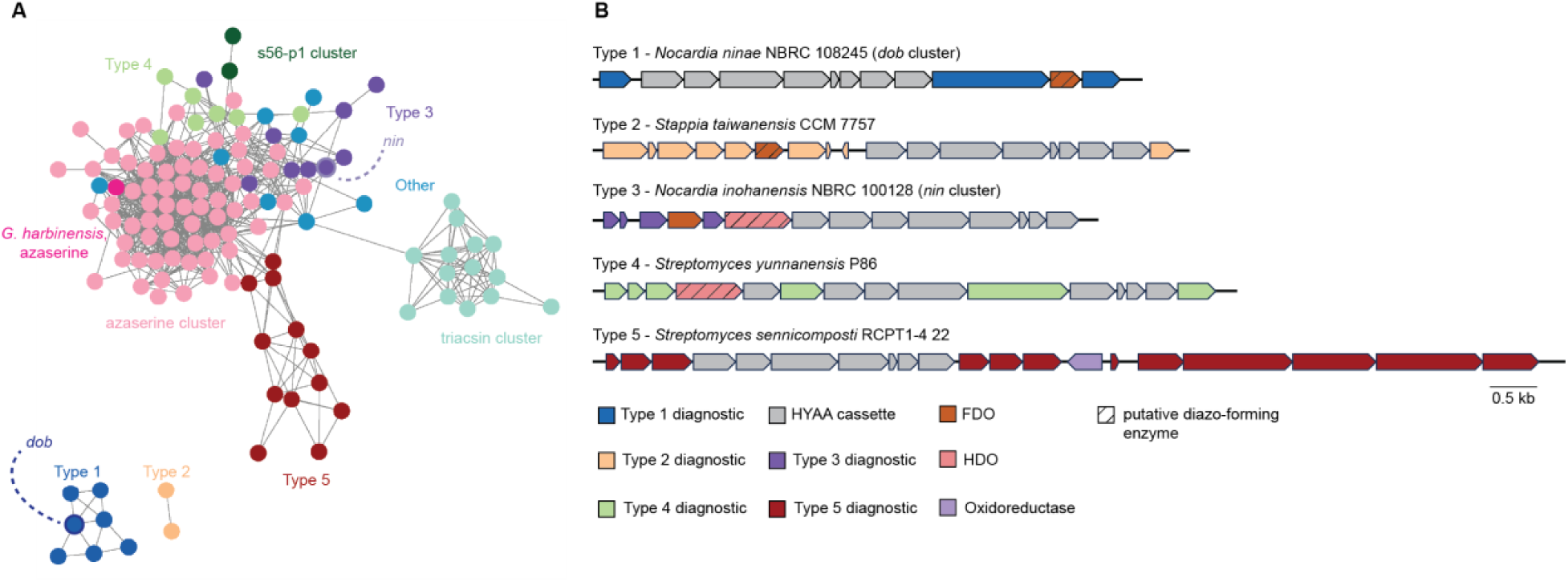
AzaE genome mining reveals diverse gene clusters containing the HYAA cassette. **A)** Biosynthetic gene cluster network constructed from gene clusters containing homologs of known hydrazone-forming enzymes^31–33,50^ and at least one additional oxidoreductase. **B)** Representative gene cluster diagrams for each gene cluster type.

## Notes

### Competing Interest Statement

The authors have declared no competing interest.

