## Supplementary Information for "Chemical capture of diazo metabolites reveals biosynthetic hydrazone oxidation"

**Supporting Information**

**Supplementary Table 1:** Primers used in this study.

| **Primers** | **Sequence (5'-3')** | **Description** |
|---|---|---|
| DobA_ninae_NcoI_HindIII_F | ctttaagaaggagatatacATGACCGC GGTCCTGC | HindIII |
| DobA_ninae_NcoI_HindIII_R | ggtgctcgagtgcggccgcaCGTTGCT CCTGGGACAGC | NcoI |
| DobB_ninae_NdeI_HindIII_F | ggtgccgcgcggcagccatATGGGAT GGGCTGAGCTGG | NdeI |
| DobB_ninae_NdeI_HindIII_R | tgctcgagtgcggccgcaTCATTCGAC GGCGCCC | HindIII |
| DobC_ninae_NdeI_HindIII_F | ggtgccgcgcggcagccatATGGATTC GGTGATCAACC | NdeI |
| DobC_ninae_NdeI_HindIII_R | tgctcgagtgcggccgcaTCAGGCATG TGTTCTTCC | HindIII |
| DobE_ninae_NdeI_HindIII_F | ggtgccgcgcggcagccatATGATCAT CAATCGTTTCGACAC | NdeI |
| DobE_ninae_NdeI_HindIII_R | tgctcgagtgcggccgcaTCAAGCGCT CCCCAC | HindIII |
| DobF_ninae_NdeI_HindIII_F | ggtgccgcgcggcagccatATGACGAA CAACACCGTCGC | NdeI |

| | | |
|---|---|---|
| DobF_ninae_NdeI_HindIII_R | tgctcgagtgcggccgcaTCACGCGG GACTCATCTC | HindIII |
| DobG_ninae_NdeI_HindIII_F | ggtgccgcgcggcagccatATGAGCC GGAGCTCGC | NdeI |
| DobG_ninae_NdeI_HindIII_R | tgctcgagtgcggccgcaTCATCGCTC TACGCTCCGG | HindIII |
| DobM_ninae_NdeI_HindIII_F | ggtgccgcgcggcagccatATGAGCTT CGATGCGGGC | NdeI |
| DobM_ninae_NdeI_HindIII_R | tgctcgagtgcggccgcaTCATTTCGC CCGCCCC | HindIII |
| DobQ_ninae_NdeI_HindIII_F | ggtgccgcgcggcagccatATGCCTGA TCTGGAAAAGGCC | NdeI |
| DobQ_ninae_NdeI_HindIII_R | tgctcgagtgcggccgcaTCAGACCG CACTTTCGGAG | HindIII |
| Dob2_ninae_NdeI_HindIII_F | ggtgccgcgcggcagccatATGAGTG CGTTCCTGTCG | NdeI |
| Dob2_ninae_NdeI_HindIII_R | tgctcgagtgcggccgcaCTATCCCTC CATTTCTCTCCGG | HindIII |
| Dob3_ninae_NdeI_HindIII_F | ggtgccgcgcggcagccatATGACATT GCCGCAATATCC | NdeI |
| Dob3_ninae_NdeI_HindIII_R | tgctcgagtgcggccgcaTCACTGTCC CTCGATCGTC | HindIII |

| | | |
|---|---|---|
| Dob3_E137A_F | ggtgtggtactg**cgc**gtcgagcatcgc | ctc → cgc |
| Dob3_E137A_R | gcgatgctcgac**gcg**cagtaccacacc | gag → gcg |
| Dob3_E101A_F | tcaccacgtgctg**cgc**ggtgtccatgatg | ctc → cgc |
| Dob3_E101A_R | catcatggacacc**gcg**cagcacgtggtga | gag → gcg |
| Dob3_H140A_F | gagatgcatgagggt**ggc**gtactgctcgtcg agc | gtg → ggc |
| Dob3_H140A_R | gctcgacgagcagtac**gcc**accctcatgca tctc | cac → gcc |
| Dob3_H225A_F | ctcgtcgcggtt**ggc**catggtcgcggtg | gtg → ggc |
| Dob3_H225A_R | caccgcgaccatg**gcc**aaccgcgacgag | cac → gcc |
| Dob3_E198A_F | gttgatcgagat**cgc**ggcgaccgtggc | ctc → cgc |
| Dob3_E198A_R | gccacggtcgcc**gcg**atctcgatcaac | gag → gcg |
| Dob3_E229A_F | ggagtggcagta**cgc**gtcgcggttgtg | ctc → cgc |
| Dob3_E229A_R | cacaaccgcgac**gcg**tactgccactcc | gag → gcg |
| Dob3_H232A_F | cggcgatcgagga**ggc**gcagtactcgtcg c | gtg → ggc |
| Dob3_H232A_R | gcgacgagtactgc**gcc**tcctcgatcgccg | cac → gcc |

**Supplementary Table 2:** Vectors used in this study

| Vector | Description | Source |
|---|---|---|
| pET-28a(+) | Protein expression vector | Invitrogen |
| pDualP | Dual inducible BAC for whole-cluster expression | Terra Bioforge |

**Supplementary Table 3:** Strains used in this study.

| Strain | Description | Source |
|---|---|---|
| *Glyomyces harbinensis* ATCC 43155 | Azaserine producer | ATCC |
| *Streptomyces coelicolor* M1152 | Heterologous host for *dob* gene cluster expression | Lab stock |
| *Escherichia coli* Top10 | Maintenance of protein expression vectors | Invitrogen |
| *Escherichia coli* BL21(DE3) | Protein expression | Invitrogen |
| *Escherichia coli* BAP1 | Expression of phosphopantetheinylated proteins | Lab stock |
| *Escherichia coli* BacOpt2.0 | Dh10B derivative – maintenance of *dob*-pDualP and *dob*-pDualP ∆Dob3 | Terra Bioforge |

| | | |
|---|---|---|
| *Escherichia coli* ET12567/pUZ8002 | Methylation deficient donor for *dob*-pDualP and *dob*-pDualP ΔDob3 conjugation | Lab Stock |
| *Nocardia ninae* NBRC 108245 | DOBA/DAC production | DSMZ |
| *Nocardia tenerifensis* DSM 44704 | DOBA/DAC production | DSMZ |
| *Streptomyces coelicolor dob*-pdualP | Cluster confirmation | This study |
| *Streptomyces coelicolor dob*-pdualP ΔDob3 | In vivo investigation | This study |

**Supplementary Table 4:** Annotations of proteins in the *dob* biosynthetic gene cluster in *Nocardia ninae*.

| Name | Size (aa) | Annotation | Homolog origin | Proposed function | Accession ID | ID (% aa) | Query coverage (%) |
|---|---|---|---|---|---|---|---|
| DobA | 350 | C45 peptidase | *Nocardia* sp. CS682 | HYAA-formation | WP_135232797.1 | 96.0 | 100 |
| DobB | 204 | GCN5 *N*-acetyl transferase | *Nocardia* sp. XZ_19_369 | HYAA-formation | WP_194834407.1 | 99.0 | 100 |
| DobC | 495 | AMP-dependent synthease | *Nocardia* sp. CS682 | HYAA-formation | WP_135232795.1 | 98.0 | 100 |
| DobE | 671 | Methionine | *Nocardia* sp. | HYAA- | WP_206055239.1 | 97.8 | 100 |

| | | | | | | | |
|---|---|---|---|---|---|---|---|
| | | tRNA ligase/cupin domain | CS682 | formation | | | |
| DobF | 358 | FAD-binding oxidoreductase | *Nocardia* sp. XZ_19_369 | HYAA-formation | WP_194834410.1 | 96.1 | 100 |
| DobG | 439 | L-lysine 6-monooxygenase | *Nocardia* sp. CS682 | HYAA-formation | WP_135232793.1 | 98.4 | 100 |
| DobM | 391 | Acyl-CoA dehydrogenase | *Nocardia* sp. CS682 | HYAA-formation | WP_135232798.1 | 97.4 | 100 |
| DobQ | 86 | Acyl carrier protein | *Nocardia suismassiliense* | HYAA-formation | WP_107658748.1 | 98.8 | 100 |
| Dob1 | 342 | ParB transcriptional regulator | *Nocardia* sp. CS682 | regulation | WP_135232792.1 | 98.3 | 100 |
| Dob2 | 1239 | Polyketide synthase | *Nocardia* sp. XZ_19_369 | PKS | WP_194834404.1 | 95.8 | 100 |
| Dob3 | 322 | Ferritin-like diiron oxidase or oxygenase | *Nocardia* sp. CS682 | Diazo formation | WP_135232800.1 | 98.8 | 100 |
| Dob4 | 405 | Major facilitator superfamily 1 | *Nocardia suismassiliense* | Export/resistance | WP_107658754.1 | 97.8 | 100 |

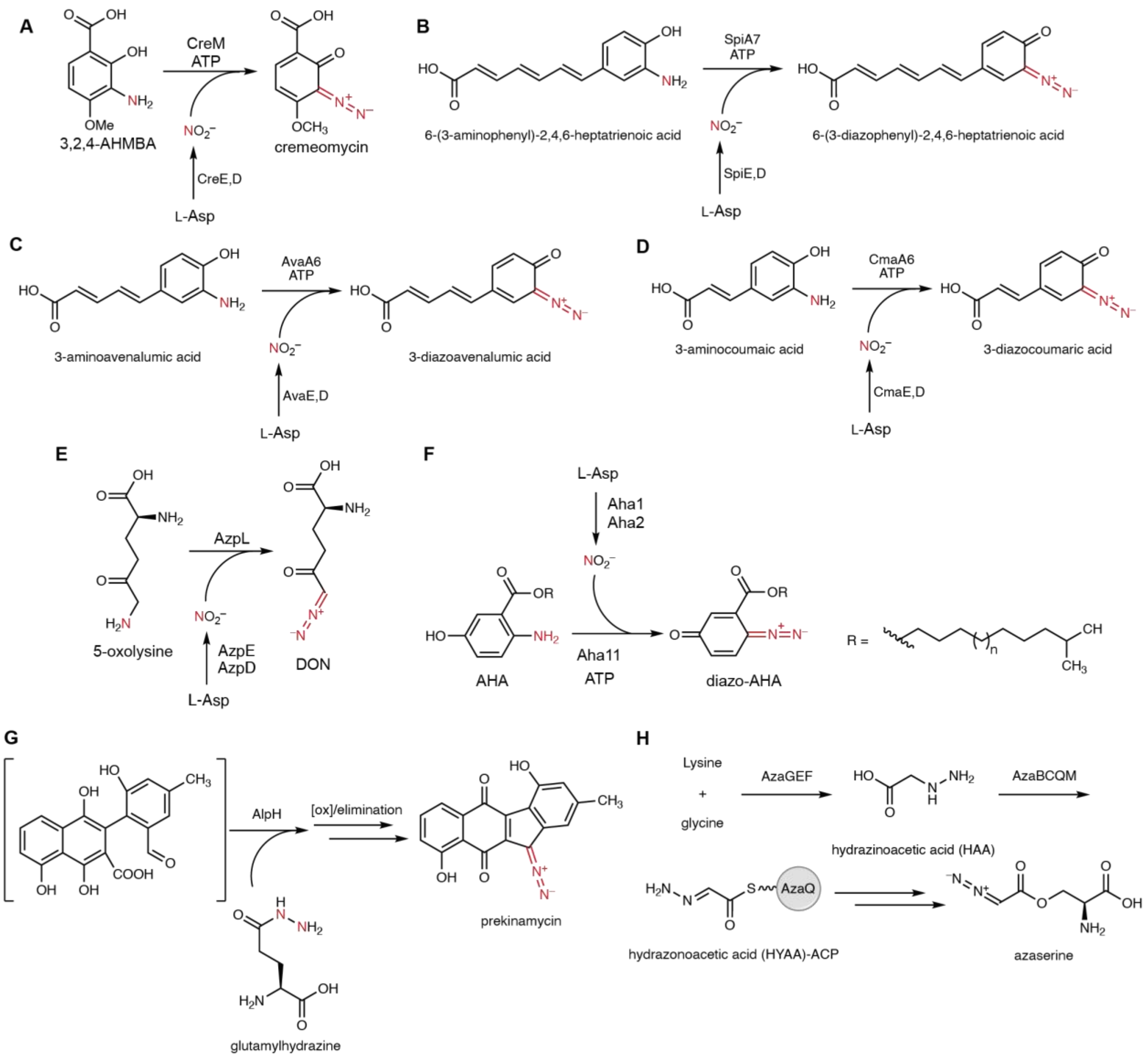


**Supplementary Figure 1:** Characterized and putative diazo biosynthetic enzymes. **A)** CreM catalyzes formation of the diazo functionality in cremeomycin biosynthesis. 3,2,4-AHMBA = 3-amino-2-hydroxy-4-methoxybenzoic acid.[1,2] **B)** SpiA7 catalyzes diazo formation in spinamycin biosynthesis.[3] **C)** AvaA6 catalyzes 3-diazoavenalumic acid formation in avenalumic acid biosynthesis.[4] **D)** CmaA6 catalyzes 3-diazocoumaric acid formation in *p*-coumaric acid biosynthesis.[5] **E)** AzpL catalyzes DON formation in alazopeptin biosynthesis.[6] **F)** Aha11 catalyzes diazo formation in tasikamide biosynthesis. AHA = alkyl 5-hydroxylanthranilate.[7] **G)** AlpH catalyzes condensation of an acyl hydrazide with a carbonyl in kinamycin biosynthesis.[8] AzaGEF catalyzes conversion of L-lysine and glycine to hydrazinoacetic acid and subsequent transformations by AzaBCQM yields hydrazonoacetic acid-AzaQ. The enzyme responsible for oxidation of the hydrazone to the diazo has not yet been elucidated.

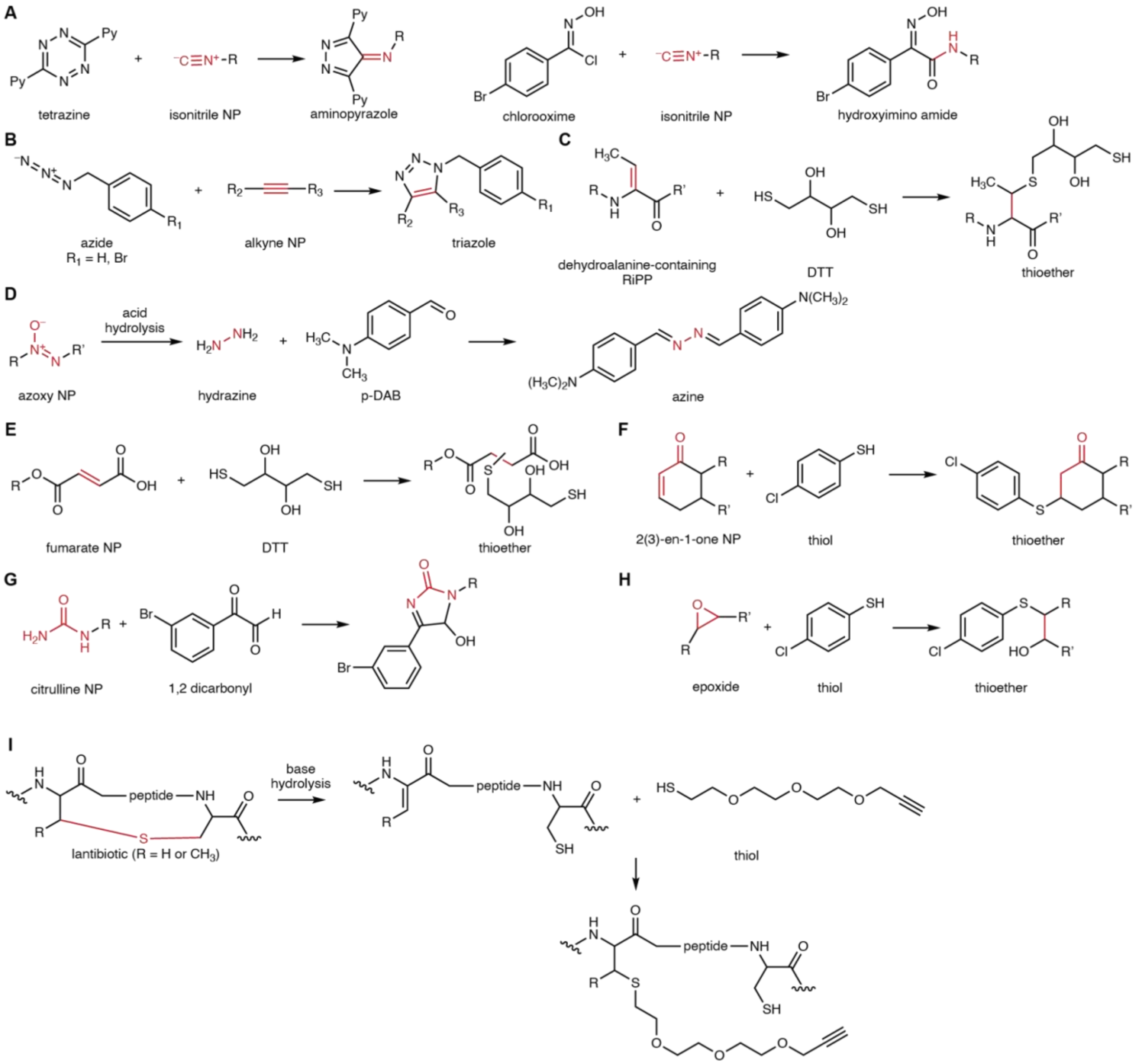


**Supplementary Figure 2:** Chemical trapping facilitates identification of natural products. **A)** Tetrazine[9] and chlorooxime[10] probes identify isonitrile-containing natural products. **B)** Azide probes undergo 1,3-dipolar cycloadditions to identify alkyne-containing natural products.[11,12] **C)** Dithiothreitol (DTT) undergoes 1,4-nucleophilic additions to identify dehydroalanine-containing RiPPs.[13,14] **D**) Acid hydrolysis of azoxy natural products yields hydrazine which can be derivatized by para-dimethylamino benzaldehyde (p-DAB) to identify azoxy natural products.[15] **E)** DTT undergoes 1,4-nucleophilic additions to identify fumarate-containing natural products.[16] **F)** Thiol probes undergo 1,4-nucleophilic additions to identify 2(3)-en-1-one-containing natural products.[17] **G)** 1,2-dicarbonyl probes identify citrulline-containing natural products.[18] **H)** Thiol probes identify epoxide-containing natural products.[19] **I)** Base hydrolysis of lantibiotics yields dehydrated amino acids which undergo 1,4-nucleophilic additions with thiol probes to introduce an alkyne functionality. A subsequent click reaction with an azide enables enrichment of the derivatized natural product.[20]

**Supplementary Table 5:** Organisms containing *dob* biosynthetic gene clusters.

| **Organism** | **Description** | **Accession** |
|---|---|---|
| *Actinobacteria* sp. 051321 | Soil microbe | IMG 2931719839 |
| *Actinomadura sp. J1-007* | Industrial maduramicin producer | IMG 3003019599 |
| *Nocardia ninae* NBRC 108245 | Human pathogen | NZ_BJXA00000000.1 |
| *Nocardia tenerifensis* DSM 44704 | Animal pathogen | NZ_QJKF01000002.1 |
| *Nocardia pseudobrasiliensis* DSM 44290 | Human pathogen | NZ_QQBC01000001.1 |
| *Nocardia colli* CICC11023 | Human pathogen | NZ_VXLC01000004.1 |
| *Nocardia suismassiliense* S-137 | Boar gut microbe | NZ_LT985361.1 |
| *Nocardia* sp. CS682 | Soil microbe | CP029710.1 |
| *Nocardia* sp. NPDC 6044 | Soil microbe | NZ_JBIAID010000001.1 |
| *Nocardia* sp. NPDC 50175 | Soil microbe | NZ_JBITJI010000016.1 |
| *Nocardia* sp. NPDC 51321 | Soil microbe | NZ_JBITEM010000001.1 |

| | | |
|---|---|---|
| *Nocardia* sp. NPDC 51756 | Soil microbe | NZ_JBFAYV010000001.1 |
| *Nocardia* sp. NPDC 57030 | Soil microbe | NZ_JBHUSN010000373.1 |
| *Nocardia* sp. NPDC 60255 | Soil microbe | NZ_JBHXAU010000066.1 |
| *Nocardia* sp. NPDC 52316 | Soil microbe | NZ_JBIUBB010000001.1 |
| *Nocardia* sp. NPDC 46473 | Soil microbe | NZ_JBEYZJ010000003.1 |
| Halostreptopolyspora alba YIM 96095 | Soil microbe | IMG 2861794813 |

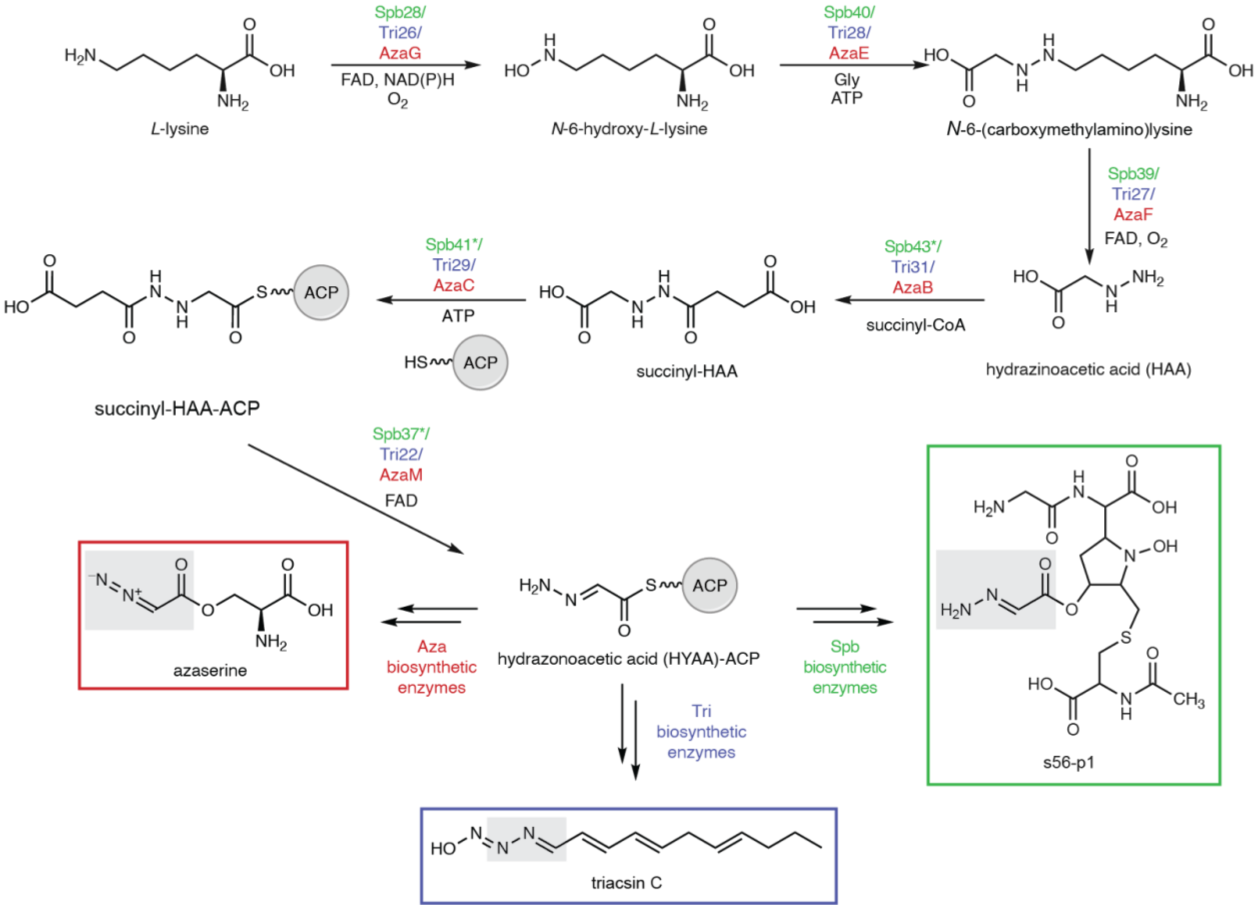


**Supplementary Figure 3:** Previously reported hydrazone biosynthetic pathways. Atoms highlighted in gray boxes originate from HAA. * = enzymes were identified bioinformatically and have not been biochemically characterized.

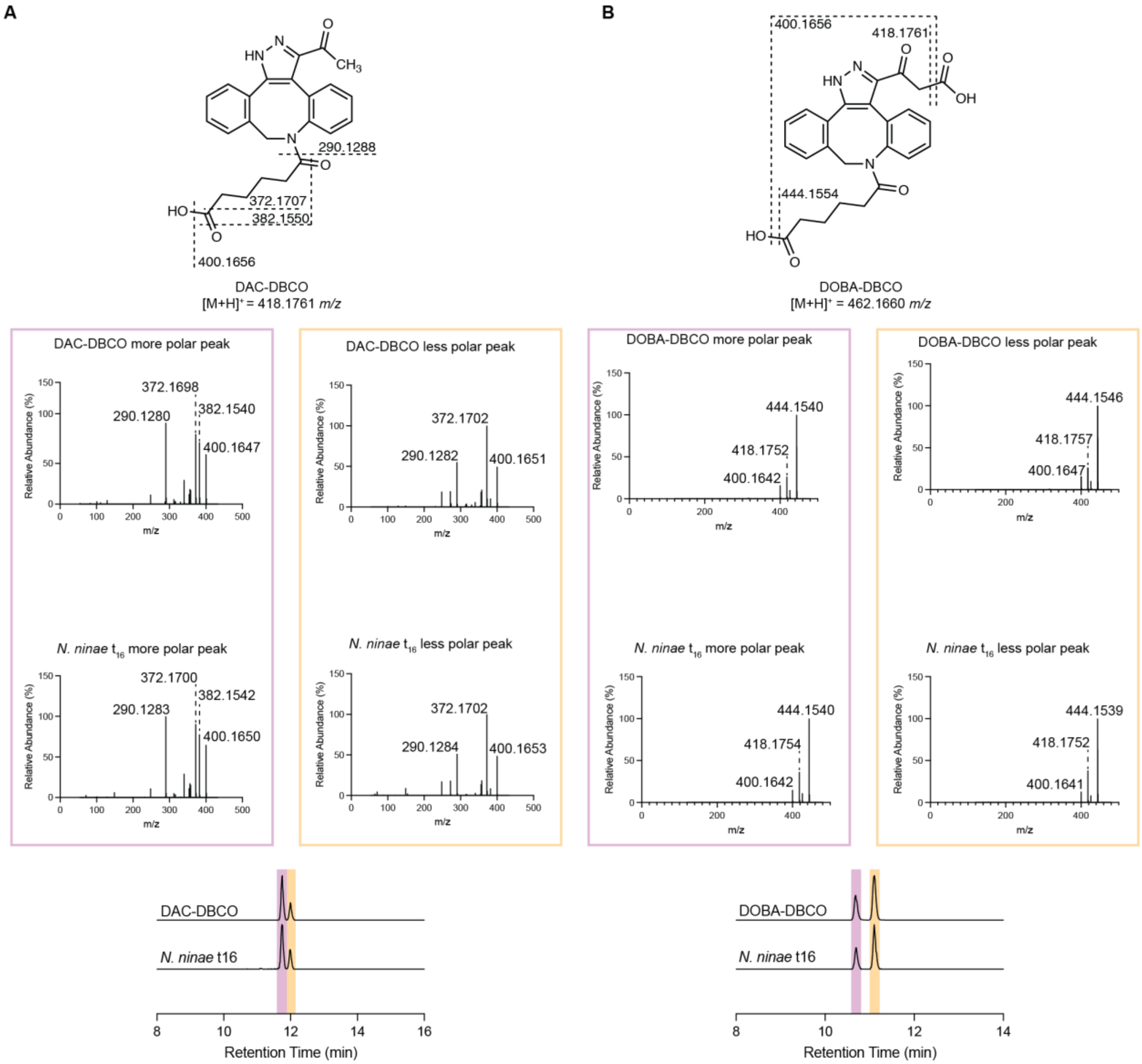


**Supplementary Figure 4:** MS/MS spectra of both regioisomers of synthetic DAC-DBCO and DOBA-DBCO, and comparison with derivatized spent media from *N. ninae*. **A)** Fragmentation pattern of DBCO-acid derivatized *N. ninae* spent media matches that of synthetic DAC-DBCO. Extracted ion chromatogram (*m/z* = 418.1761 ± 5 ppm) of the DBCO-acid derivatized *N. ninae* spent media compared to a synthetic standard. **B)** Fragmentation pattern of DBCO-acid derivatized *N. ninae* spent media matches that of synthetic DOBA-DBCO. DOBA-DBCO was prepared through incubation of OMe-DOBA-DBCO with PLE. Extracted ion chromatogram (*m/z* = 462.1660 ± 5 ppm) of the DBCO-acid derivatized *N. ninae* spent media compared to a synthetic standard.

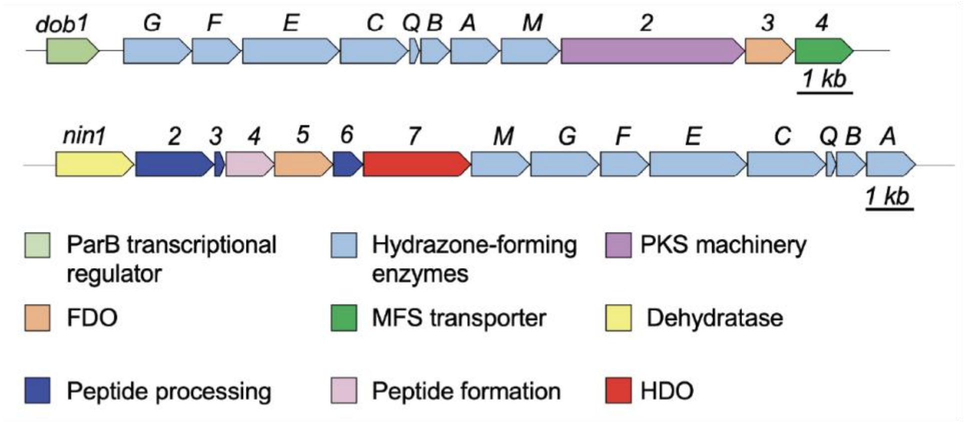


**Supplementary Figure 5:** Genome-mining for hydrazone-forming enzymes identified the *dob* and *nin* biosynthetic gene clusters in *N. ninae*. The PKS gene encoded in the *dob* gene cluster suggests *dob* is responsible for DOBA/DAC production.

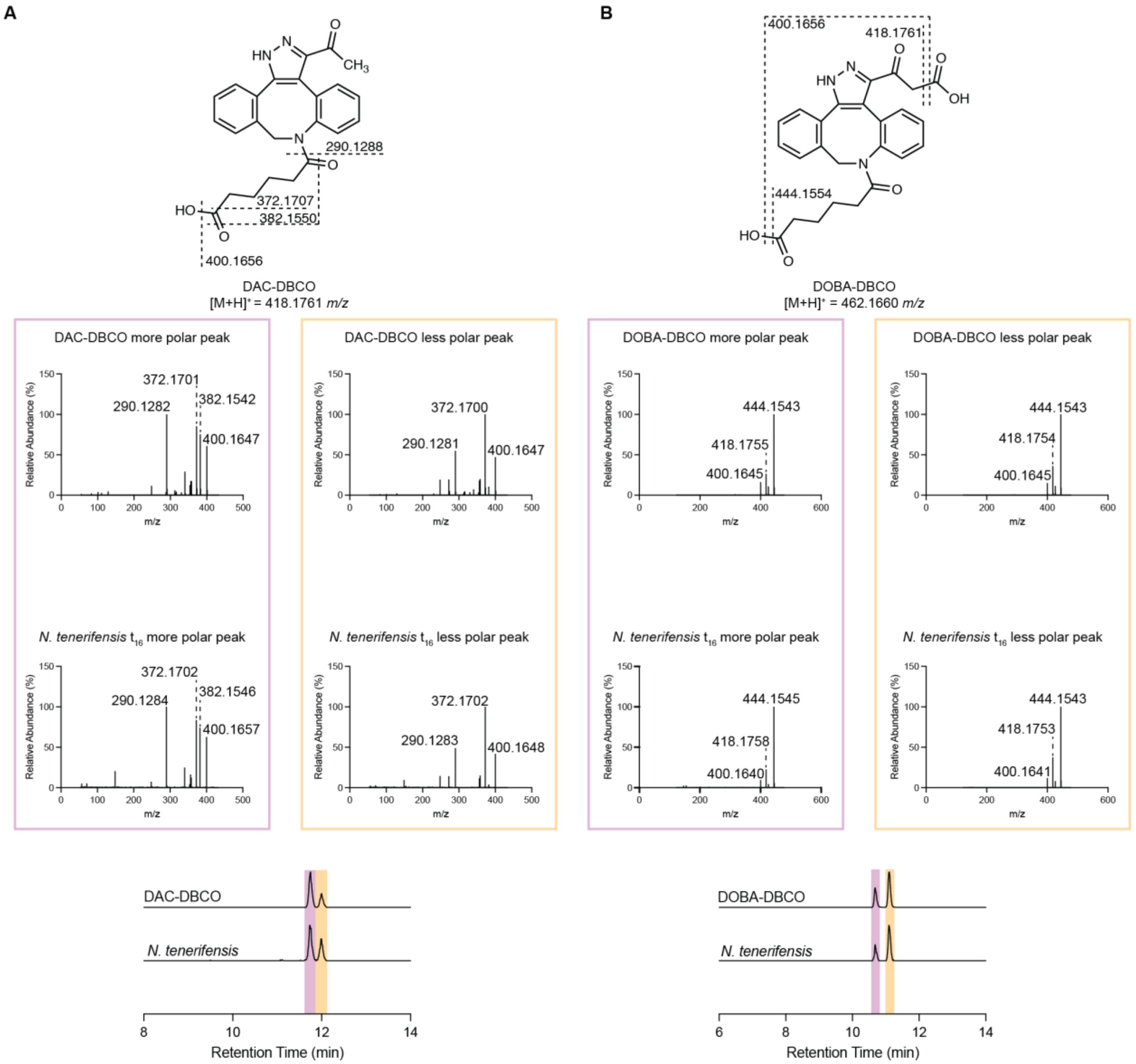


**Supplementary Figure 6:** MS/MS spectra of both regioisomers of synthetic DAC-DBCO and DOBA-DBCO, and comparison with derivatized spent media from *N. tenerifensis*. **A)** Fragmentation pattern of DBCO-acid derivatized *N. tenerifensis* spent media matches that of synthetic DAC-DBCO. Extracted ion chromatogram (*m/z* = 418.1761 ± 5 ppm) of the DBCO-acid derivatized *N. tenerifensis* spent media compared to a synthetic standard. **B)** Fragmentation pattern of DBCO-acid derivatized *N. tenerifensis* spent media matches that of synthetic DOBA-DBCO. DOBA-DBCO was prepared through incubation of OMe-DOBA-DBCO with PLE. Extracted ion chromatogram (*m/z* = 462.1660 ± 5 ppm) of the DBCO-acid derivatized *N. tenerifensis* spent media compared to a synthetic standard.

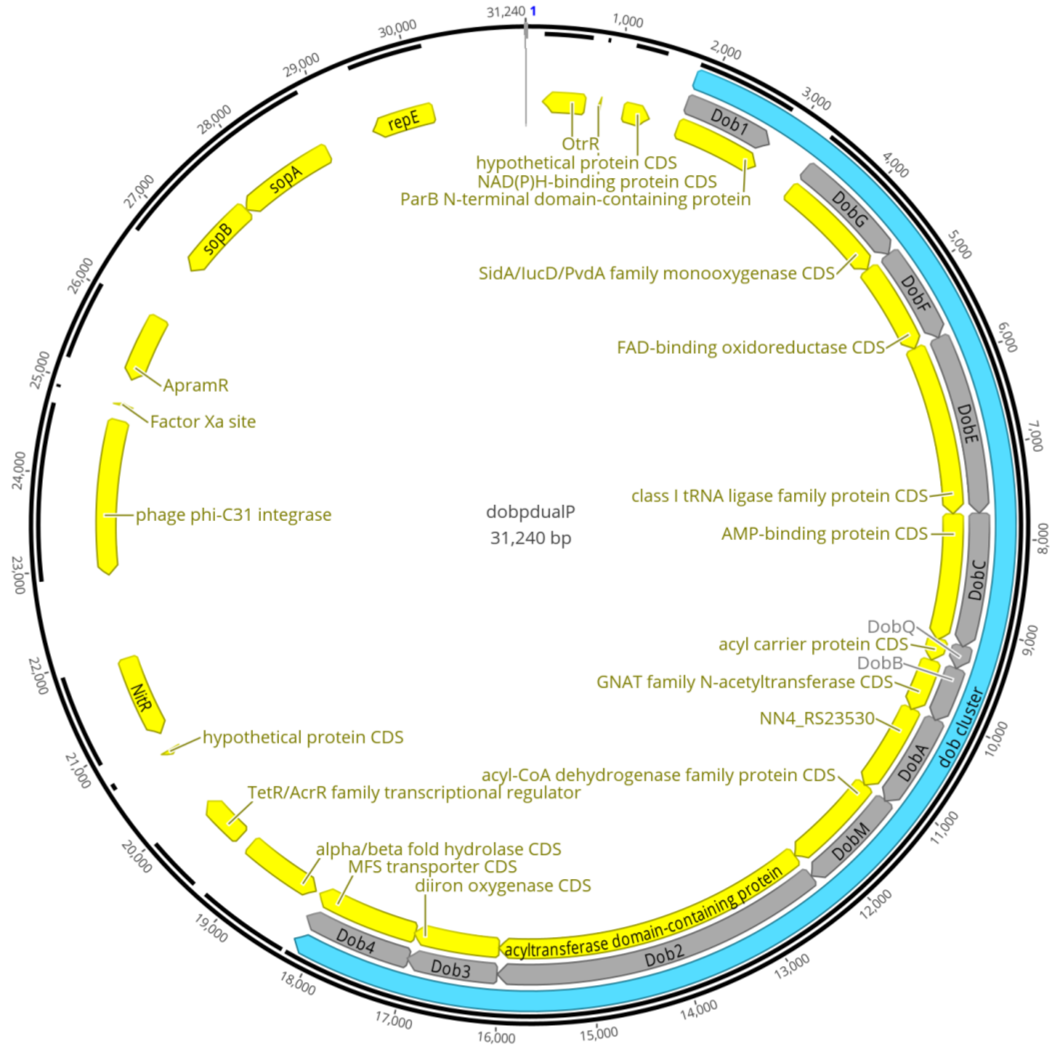


**Supplementary Figure 7:** Vector map of *dob*-pDualP

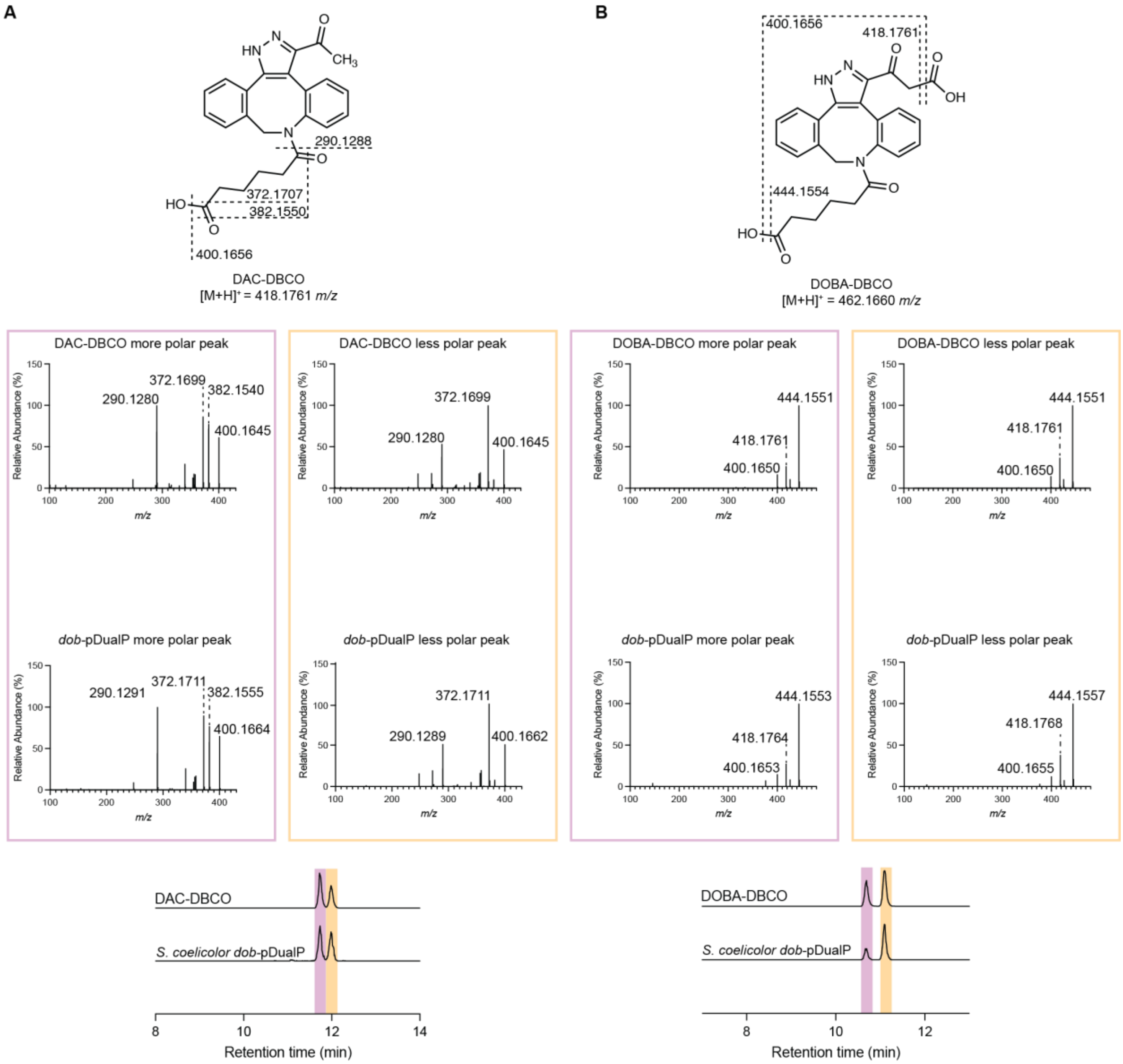


**Supplementary Figure 8:** MS/MS spectra of both regioisomers of synthetic DAC-DBCO and DOBA-DBCO, and comparison with derivatized spent media from *S. coelicolor dob*-pDualP. **A)** Fragmentation pattern of DBCO-acid derivatized *S. coelicolor dob*-pDualP spent media matches that of synthetic DAC-DBCO. Extracted ion chromatogram (*m/z* = 418.1761 ± 5 ppm) of the DBCO-acid derivatized *S. coelicolor dob*-pDualP spent media compared to a synthetic standard. **B)** Fragmentation pattern of DBCO-acid derivatized *S. coelicolor dob*-pDualP spent media matches that of synthetic DOBA-DBCO. DOBA-DBCO was prepared through incubation of OMe-DOBA-DBCO with PLE. Extracted ion chromatogram (*m/z* = 462.1660 ± 5 ppm) of the DBCO-acid derivatized *S. coelicolor dob*-pDualP spent media compared to a synthetic standard.

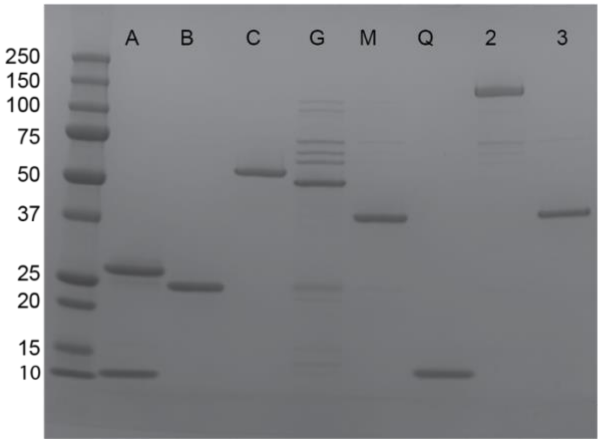


**Supplementary Figure 9:** SDS-PAGE gel of purified Dob biosynthetic enzymes. Molecular weights for $His_6$-tagged constructs were calculated using ExPASy ProtParam[21]: DobA (39.3 kDa; 12.1 kDa and 27.2 kDa after autoproteolysis at C112 to produce the mature C45 peptidase), DobB (25.1 kDa), DobC (56.1 kDa), DobG (51.8 kDa), DobM (43.7 kDa), DobQ (11.5 kDa), Dob2 (133.9 kDa), Dob3 (40.2 kDa). Precision Plus All Blue Molecular Weight Standard (Bio-Rad) was used as the ladder.

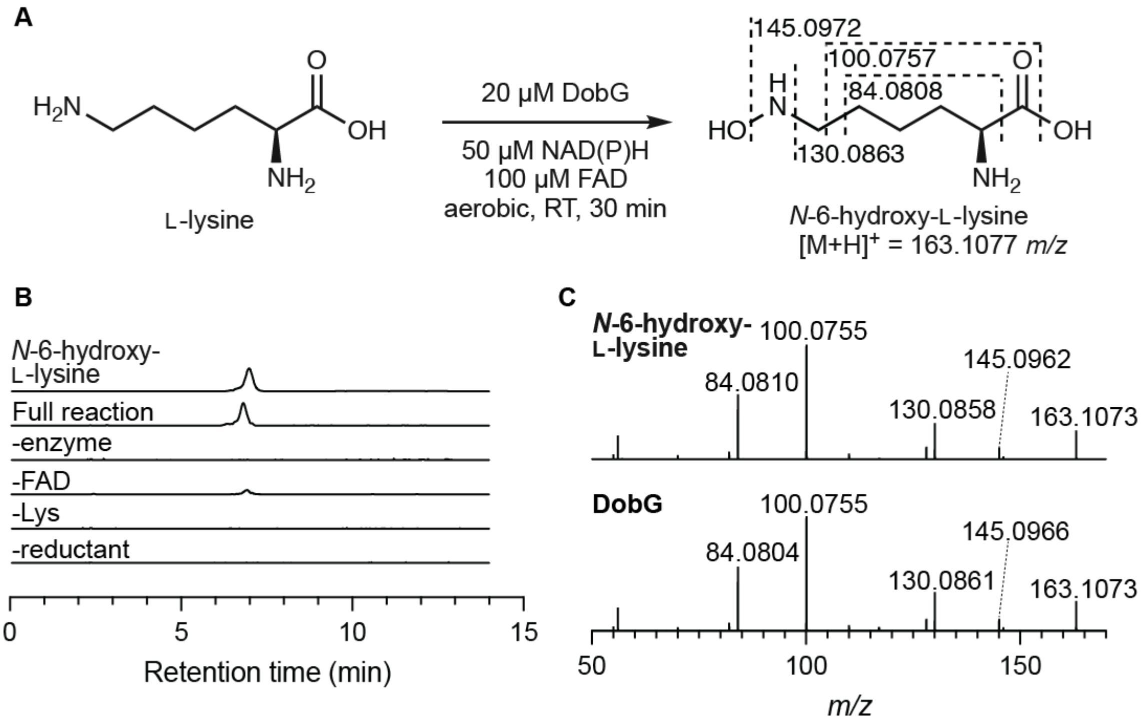


**Supplementary Figure 10:** In vitro activity assays of DobG confirm its role as an L-lysine *N*-monooxygenase. **A)** In vitro reaction of DobG. Calculated masses of predicted MS/MS fragments are shown. **B)** Extracted ion chromatogram (*m/z* = 163.1077 ± 10 ppm) of the DobG reaction compared to a synthetic standard of *N*-6-hydroxylysine and no enzyme, no FAD, no NADH/NADPH, and no L-lysine controls. **C)** MS/MS fragmentation of the 163.1077 ion from the DobG reaction matches the fragmentation pattern of the standard. Assays were run in biological triplicate. Representative results are shown.

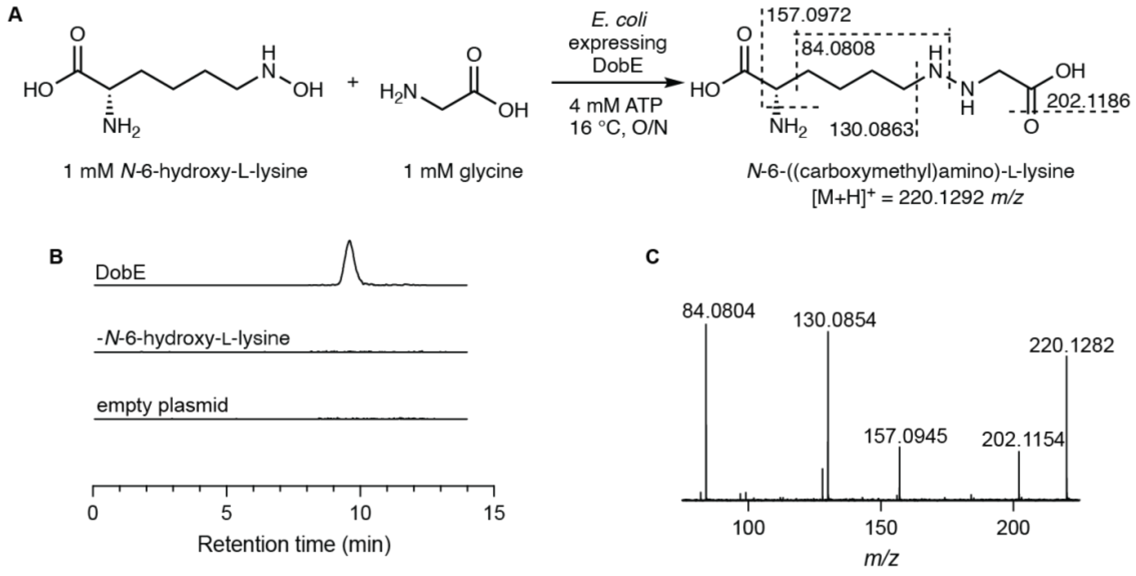


**Supplementary Figure 11:** In vivo activity assays demonstrate that DobE catalyzes N–N bond formation between glycine and *N*-6-hydroxy-L-lysine. **A)** *In vivo* reaction of DobE. Calculated masses of predicted fragments are shown. **B)** Extracted ion chromatogram (*m/z* = 220.1292 ± 10 ppm) of DobE cultures compared to an empty pET28a control. **C)** MS/MS of the 220.1292 ion from the DobE reaction. Fragmentation patterns matched previously reported spectra for *N*-6-(carboxymethylamino)lysine.[22] Assays were performed in biological triplicate and representative results are shown.

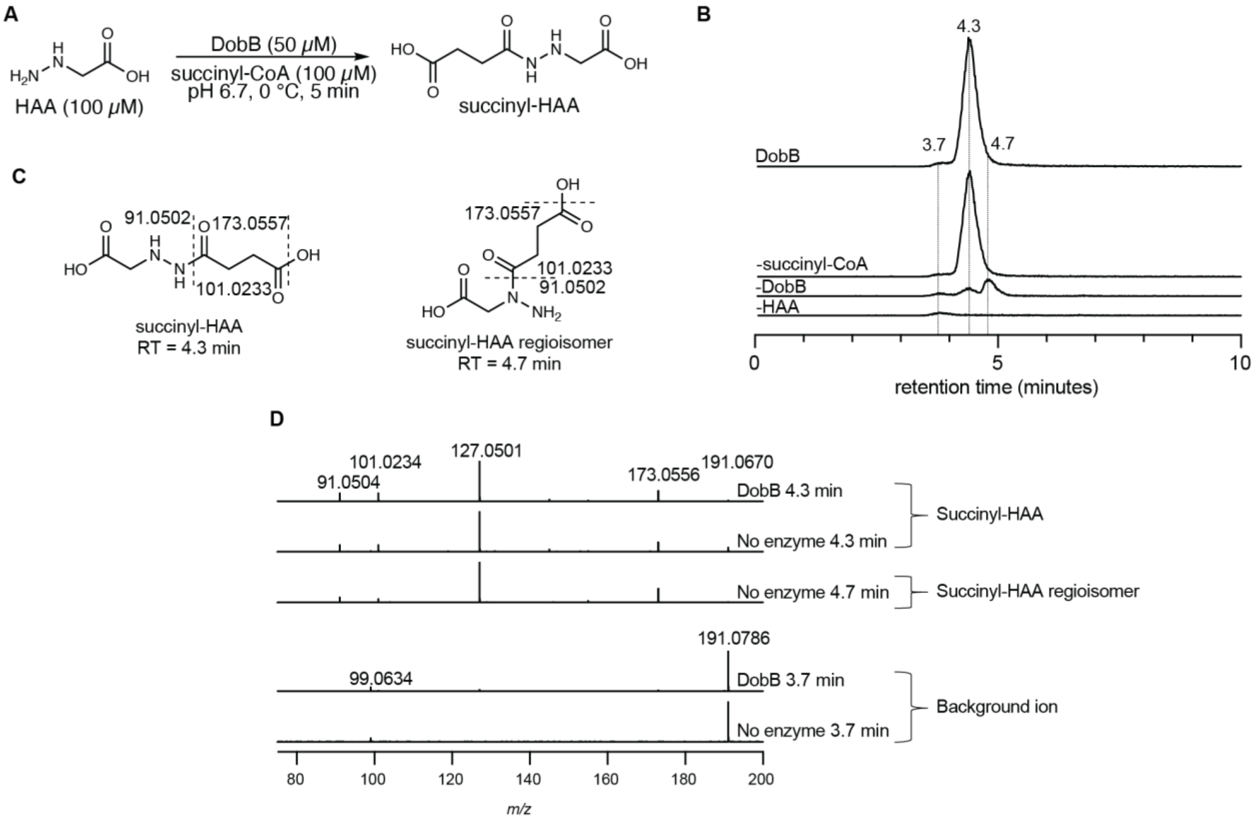


**Supplementary Figure 12:** In vitro activity assays demonstrate that DobB catalyzes succinylation of HAA. **A)** In vitro reaction of DobB. **B)** EIC (*m/z* = 191.0662 ± 10 ppm) of the DobB reaction compared to no enzyme, no succinyl-CoA, and no HAA controls. Omission of succinyl-CoA did not abolish production of succinyl-HAA, likely due to copurification of DobB with succinyl-CoA as previously reported for close homologs.[23,24] **C)** Putative structures of succinyl-HAA and the succinyl-HAA regioisomer are shown with expected fragment masses. **D)** MS/MS fragments of the 191.0662 mass from the DobB reaction are consistent with the expected fragmentation of succinyl-HAA, but do not distinguish between regioisomers. The MS/MS fragmentation patterns of the 4.3 min and 4.7 min peaks in the no enzyme control are identical to the fragmentation pattern of the AzaB reaction product. MS/MS fragmentation of the background ion peak at 3.7 min is distinct from succinyl-HAA. Experiments were performed in biological triplicate and representative results are shown.

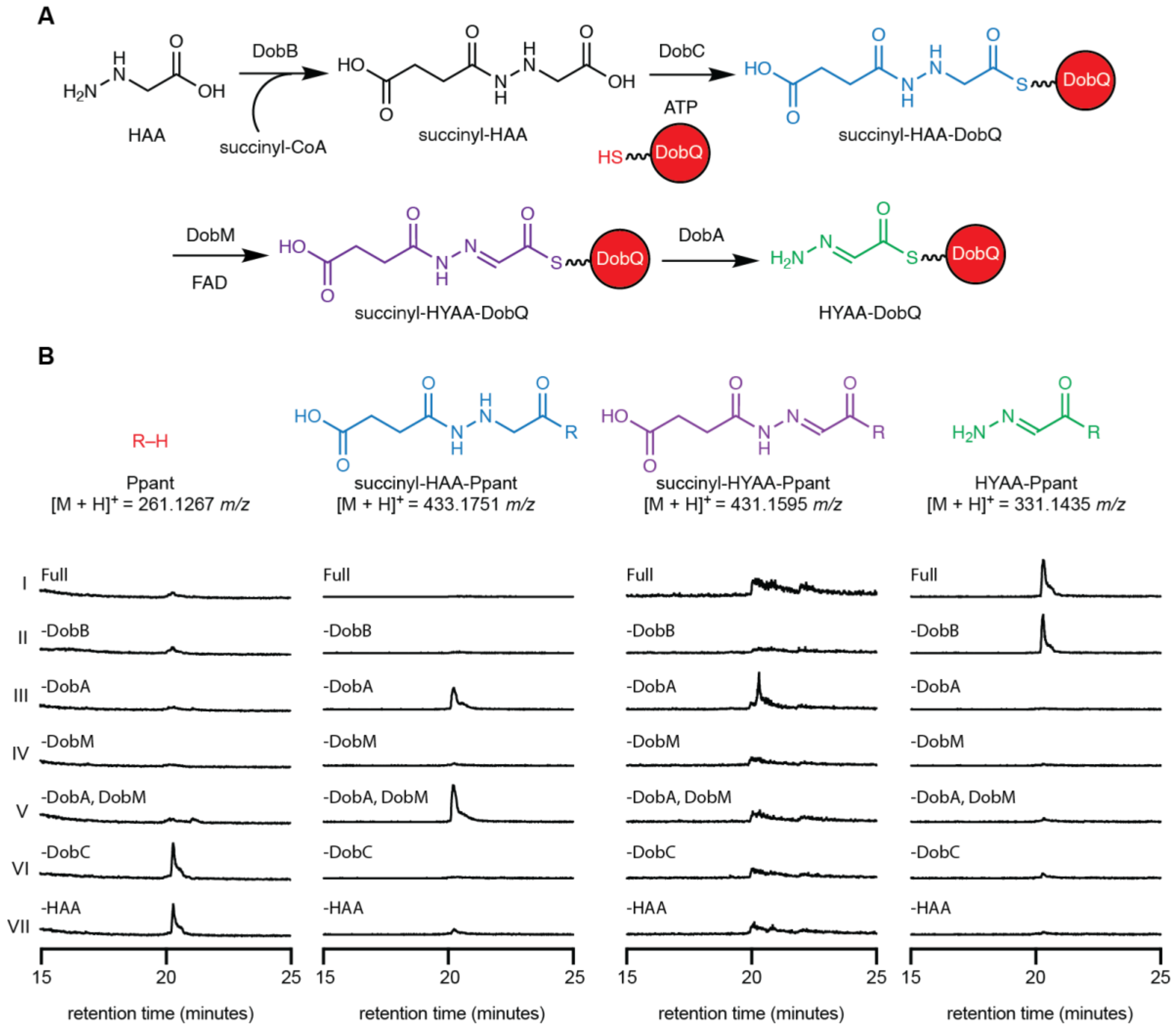


**Supplementary Figure 13:** In vitro reactions of DobABCMQ suggest DobM may catalyze succinyl-HAA-AzaQ oxidation prior to hydrolysis by DobA. **A)** DobABCMQ cascade reaction scheme. **B)** EICs of **(I)** Full DobABCMQ reaction. **(II)** Omission of DobB did not abolish production of HYAA-Ppant, likely due to non-enzymatic succinylation which was previously observed in the in vitro AzaB assay[23] (**Supplementary Figure 12**). **(III)** Omission of DobA resulted in accumulation of a mass consistent with succinyl-HYAA-Ppant. **(IV)** Interestingly, omission of DobM resulted in no observed products. We hypothesized that DobA may additionally act in a proof-reading capacity to hydrolyze stalled/aberrant DobQ thioester intermediates. **(V)** Omission of DobM and DobA led to accumulation of succinyl-HAA-Ppant, potentially supporting this hypothesis. **(VI)** Omission of DobC or **(VII)** substrate resulted in accumulation of free Ppant. HAA-Ppant (*m/z* = 333.1591) was not observed in any of the reactions. EICs are ± 10 ppm. Experiments were performed in biological triplicate and representative results are shown.

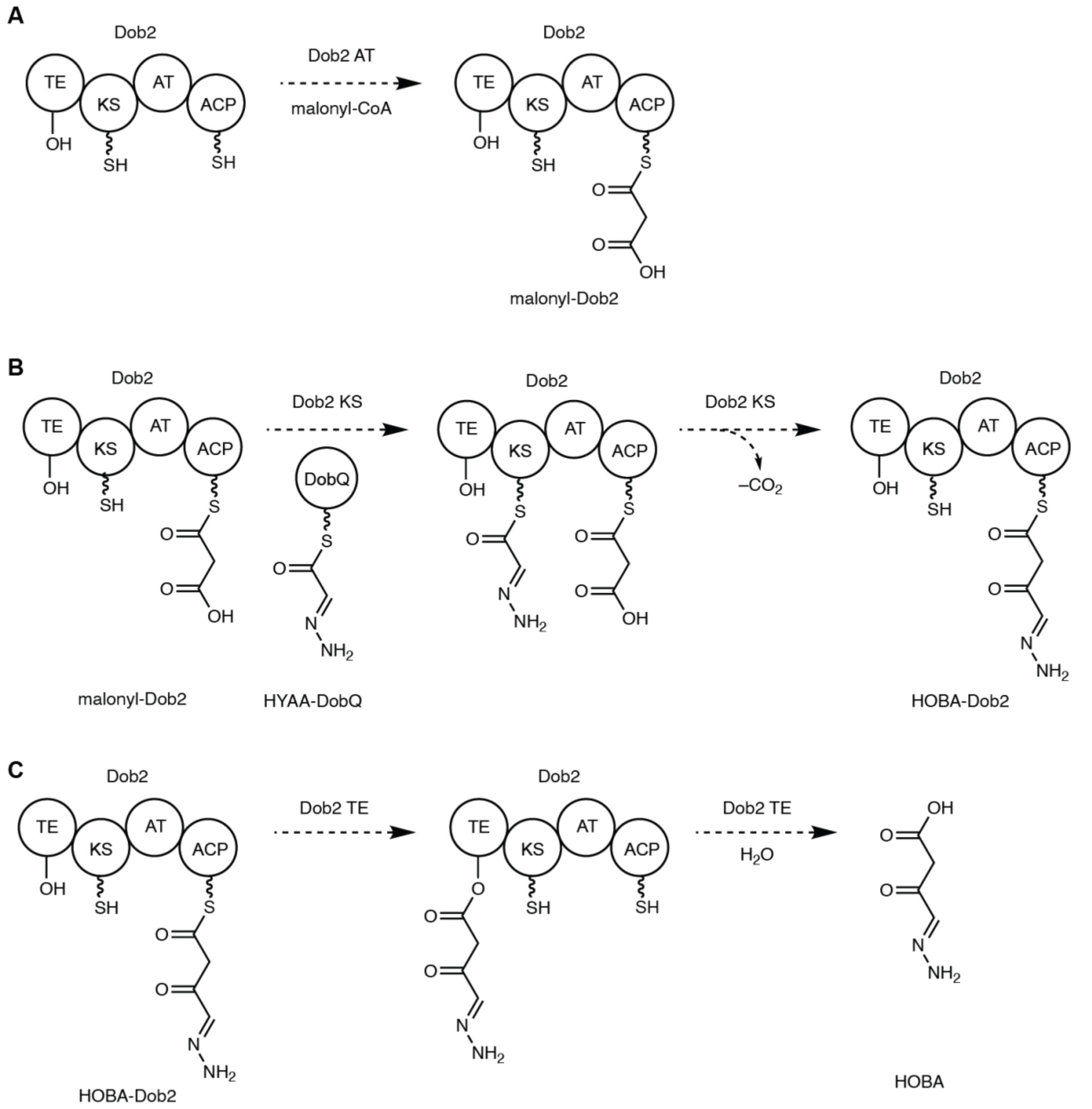


**Supplementary Figure 14:** Predicted functions of Dob2 domains. **A)** The Dob2 AT domain likely loads a malonyl extender unit. **B)** The KS domain likely catalyzes translocation of HYAA from DobQ to a conserved Cys residue, followed by C–C bond-forming decarboxylative Claisen condensation between the malonyl extender unit and the HYAA starter unit to produce HOBA-Dob2. **C)** The thioesterase domain likely translocates the mature PKS intermediate to a conserved serine residue (S88) prior to hydrolytic release of HOBA.

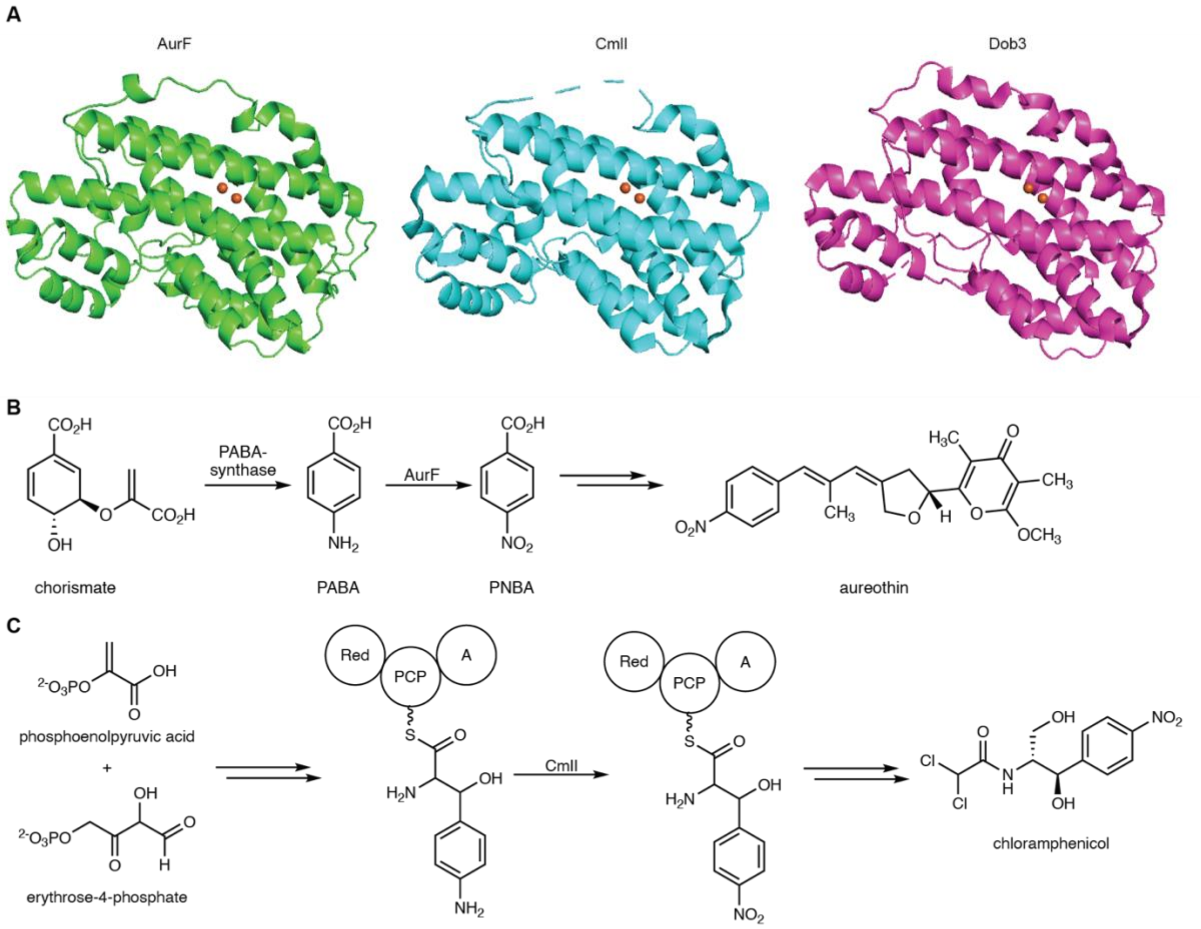


**Supplementary Figure 15:** Dob3 is structurally homologous to ferritin-like diiron *N*-oxygenases (FDOs) AurF and CmlI. **A)** Comparison of the AlphaFold predicted Dob3 structure to the X-ray crystal structures of AurF (PDB: 3CHH) and CmlI (PDB: 5HYH). Nitro-forming *N*-oxygenation reactions catalyzed by **B)** AurF[25] and **C)** CmlI[26] during the biosynthesis of aureothin and chloramphenicol, respectively.

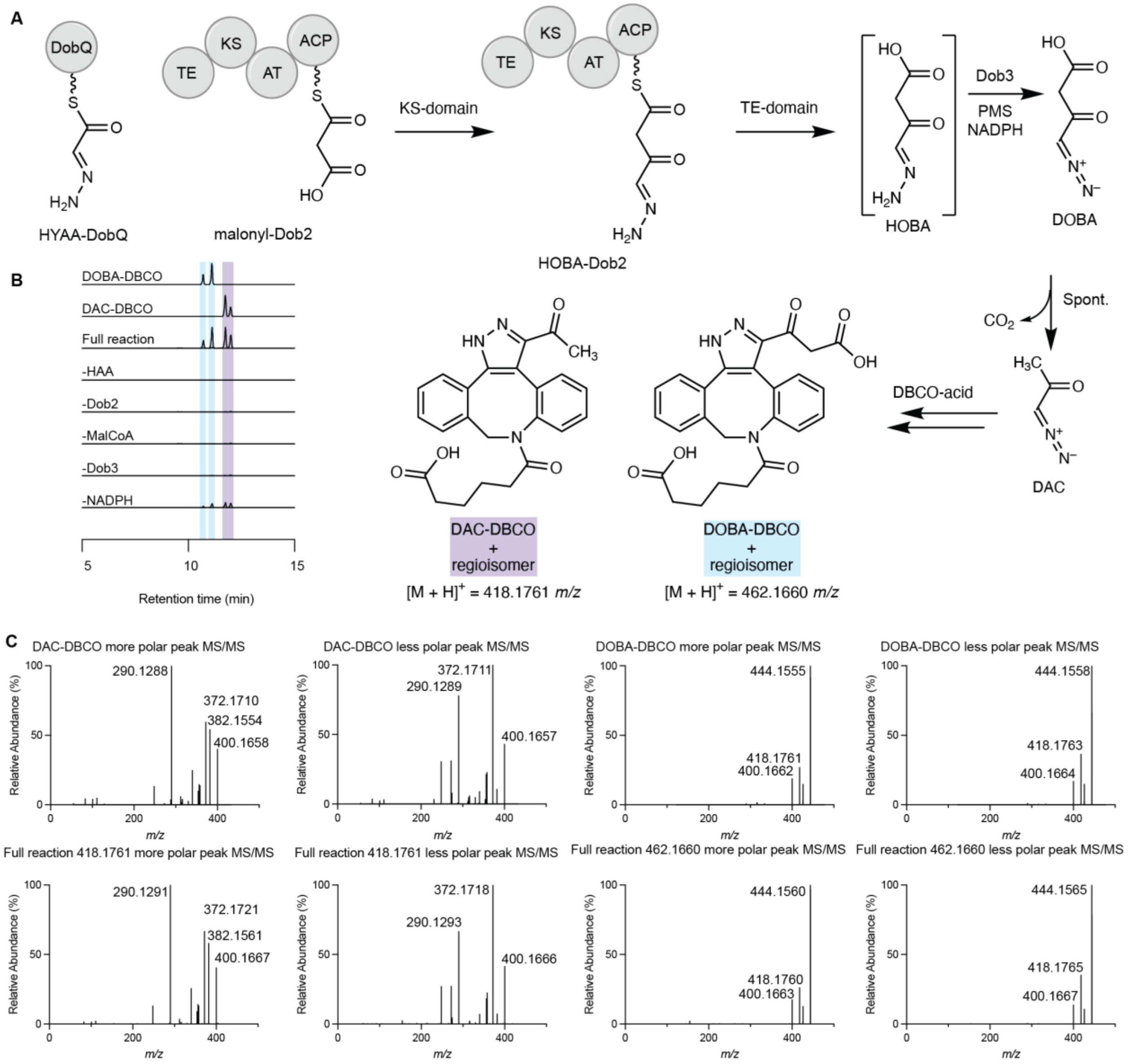


**Supplementary Figure 16:** The enzymatic cascade reaction of DobABCMQ23 converts HAA to DOBA and DAC in vitro. **A)** DobABCMQ23 reaction scheme. HYAA-DobQ is generated enzymatically from HAA by the hydrazone biosynthetic enzymes DobABCMQ. Reaction mixtures were incubated at pH 8.0. **B)** Extracted ion chromatograms for DOBA-DBCO (*m/z* = 462.1660 ± 5 ppm) and DAC-DBCO (*m/z* = 418.1761 ± 5 ppm) in 1 mM HAA, 20 μM DobB, 200 μM succinyl-CoA, 50 μM DobQ, 20 μM DobC, 5 mM ATP, 20 μM DobM, 200 μM FAD, 40 μM DobA, 10 μM Dob2, and 200 μM malonyl-CoA, as well as controls. EICs for 462.1660 and 418.1761 are superimposed. Full reaction and control data is normalized to the full reaction maximum separately for DOBA-DBCO and DAC-DBCO. **C)** MS/MS fragmentation of DAC-DBCO and DOBA-DBCO vs the products observed in the assay. Experiments were performed in biological triplicate and representative results are shown.

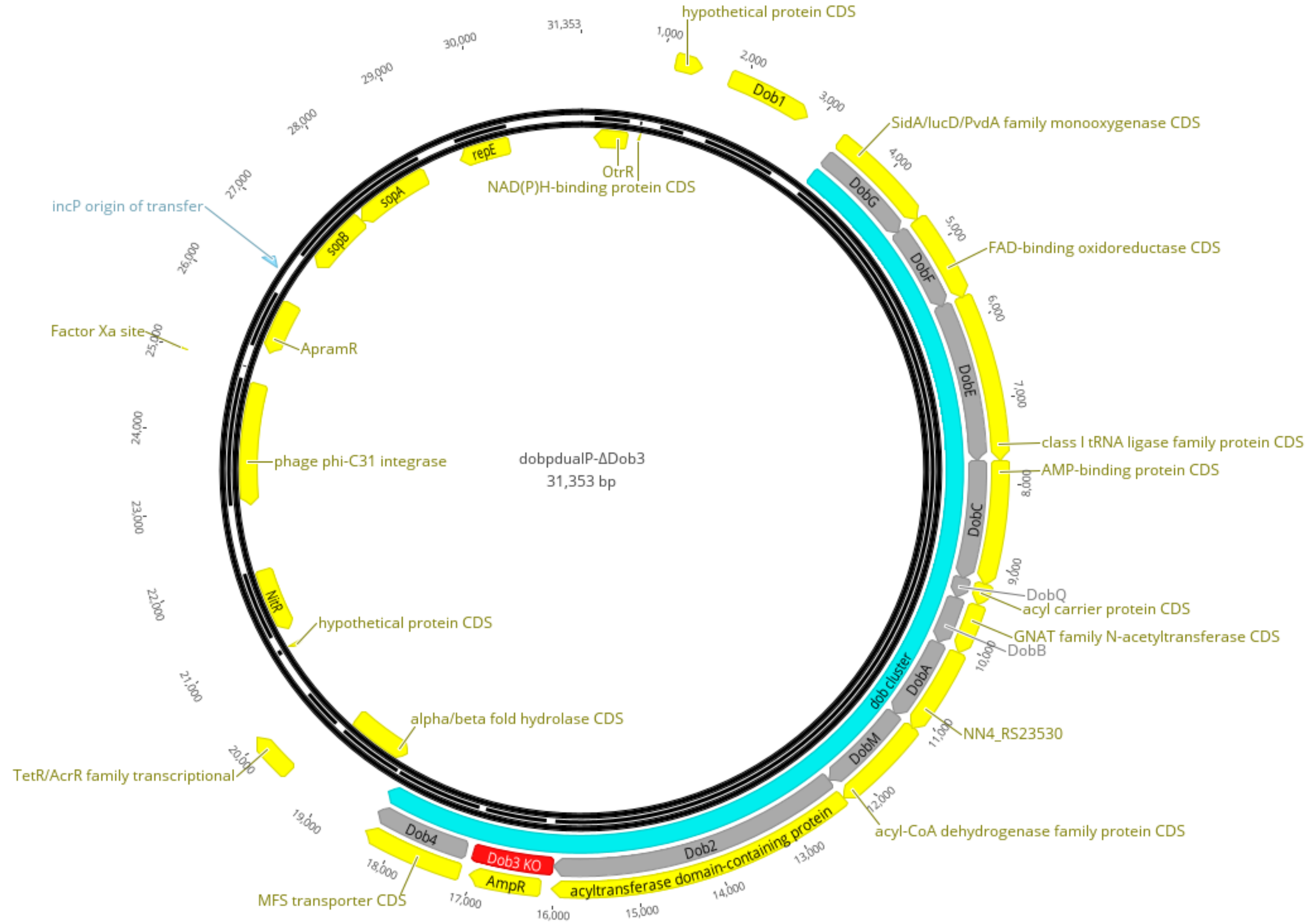


**Supplementary Figure 17:** Vector map of *dob*-pDualP ΔDob3.

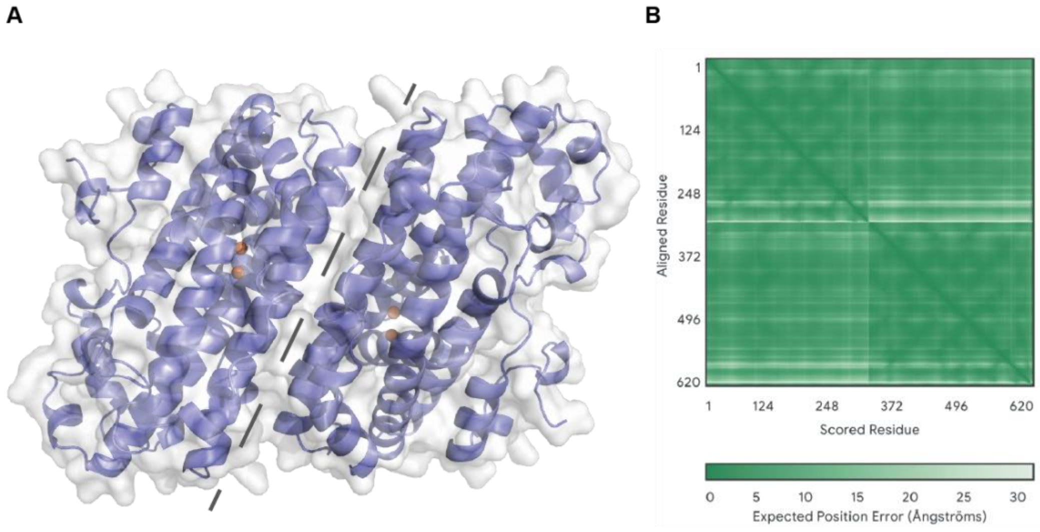


**Supplementary Figure 18**: AlphaFold 3 predicts that Dob3 is a homodimer.**A)** AlphaFold 3 predicted structure of the Dob3 homodimer. The dimer interface is marked with a dashed line. Residues 1-26 form a long, unfolded chain and are omitted for clarity. **B)** Expected Position Error Plot for the predicted Dob3 homodimer structure.

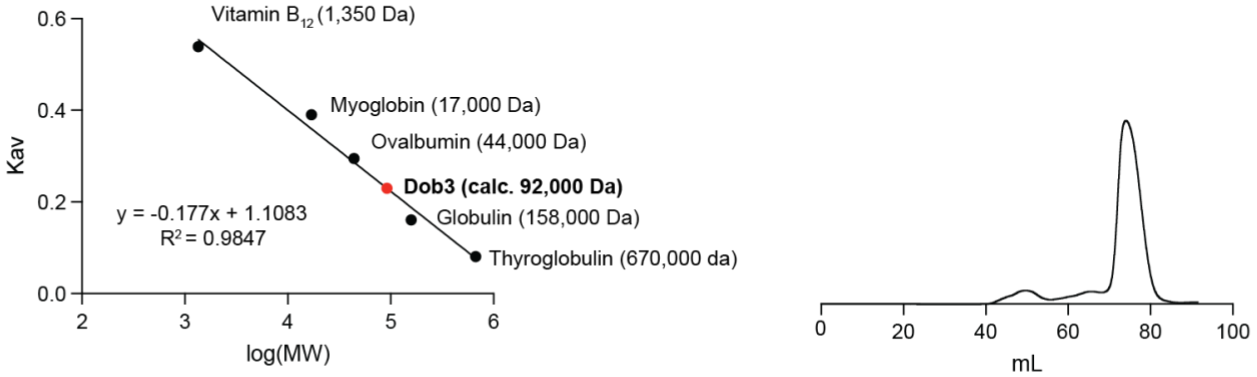


**Supplementary Figure 19:** Size exclusion chromatography is consistent with a Dob3 homodimer ($MW_{dimer}$ = 80.4 kDa, $MW_{calc}$ = 92.0 kDa).

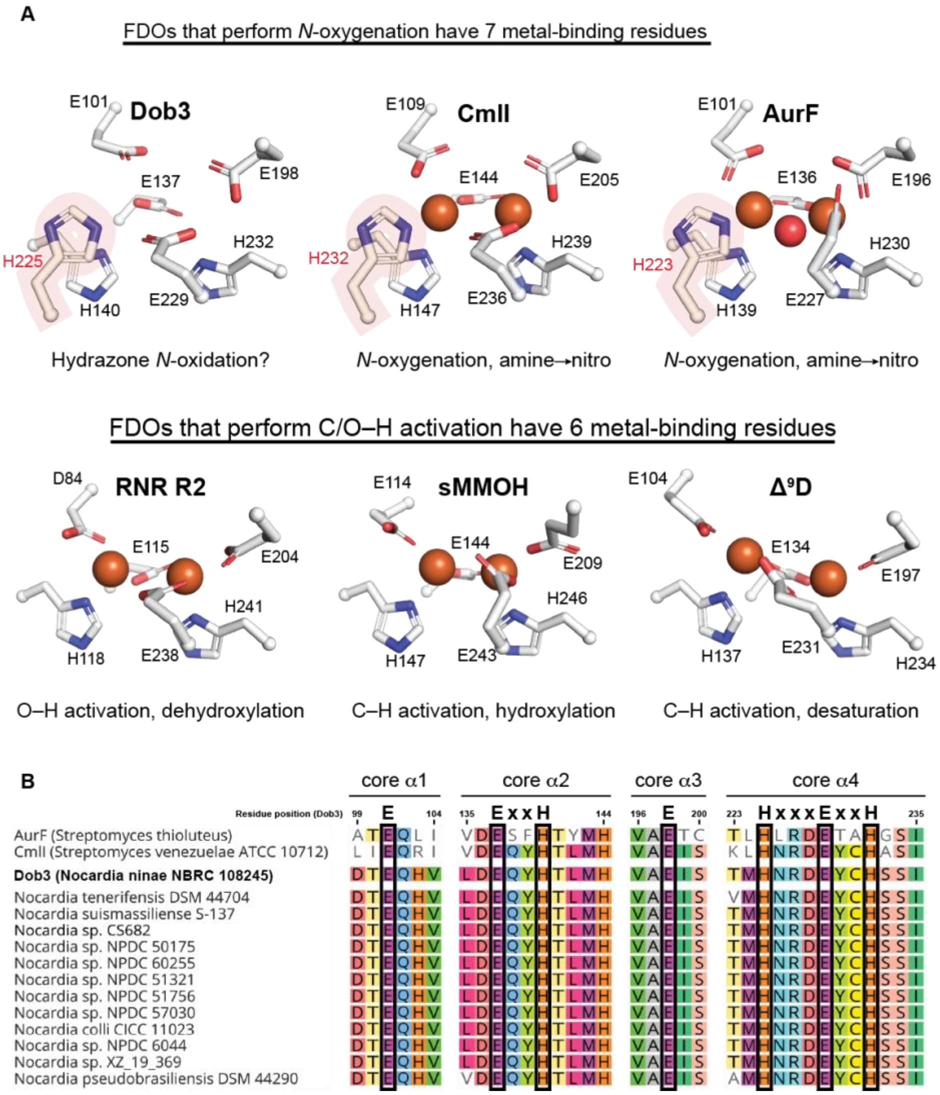


**Supplementary Figure 20:** Comparison of Dob3 to known FDOs suggests its activity as an *N*-oxygenase. **A)** The AlphaFold 3 predicted active site structure of Dob3 compared to the X-ray crystal structures of CmlI (PDB: 5HYH), AurF (PDB: 3CHH), RNR R2 (PDB: 1PIY), sMMOH (PDB: 6YD0), and $\Delta^9$-stearoyl-ACP desaturase (PDB: 1AFR). **B)** Multiple sequence alignment of Dob3 and its homologs from *Nocardia* strains with the FDO *N*-oxygenases CmlI and AurF demonstrates conservation of Fe-binding (boxed) residues.

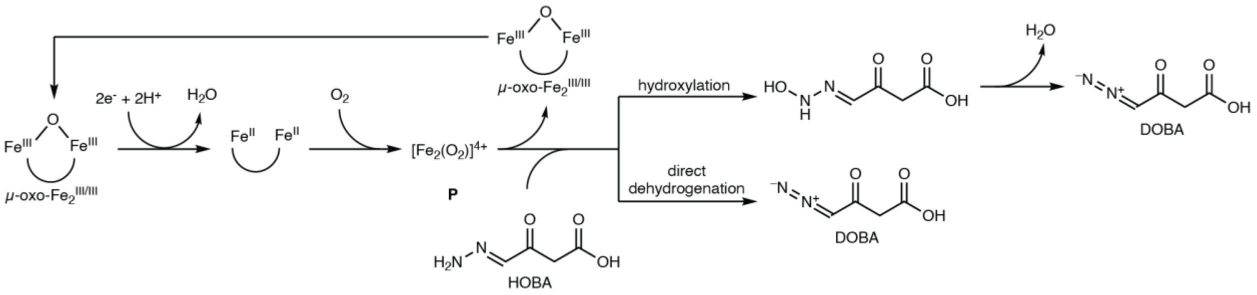


**Supplementary Figure 21:** Proposed mechanism of Dob3. Analogous to the mechanism of AurF, the μ-oxo-diferric cofactor is reduced to the diferrous state and subsequent oxidation by molecular oxygen yields a reactive differic-peroxo intermediate (**P**).[27,28] Hydroxylation and subsequent dehydration, or direct dehydrogenation of HOBA yields DOBA.

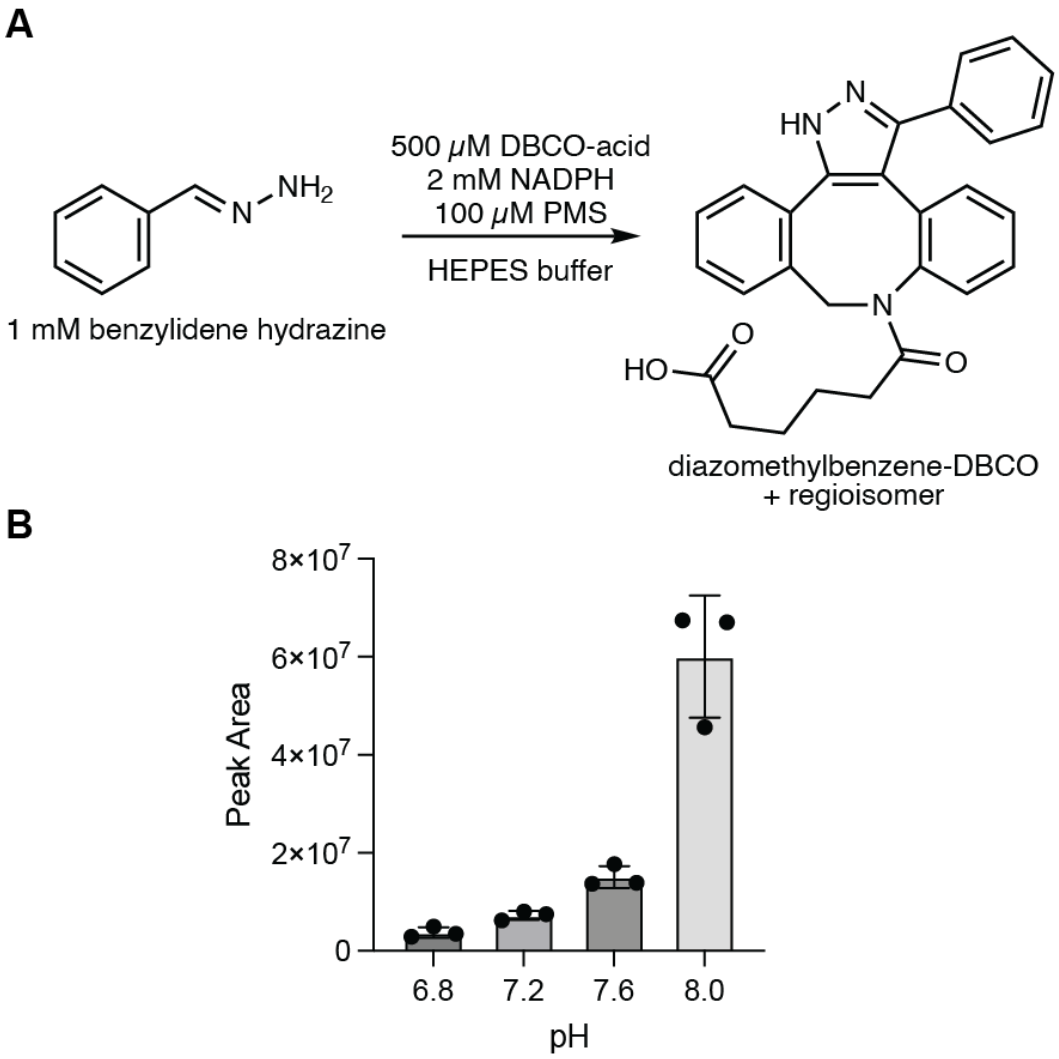


**Supplementary Figure 22:** pH dependence of autoxidation of benzylidenehydrazine. **A)** Reaction scheme. **B)** Combined LC–MS peak areas of both regioisomers of diazomethylbenzene-DBCO. Experiments were performed in triplicate. Error bars indicate mean ± standard deviation. Product identity was confirmed by comparison of LC–MS/MS data to a synthetic standard of diazomethylbenzene-DBCO.

**A**

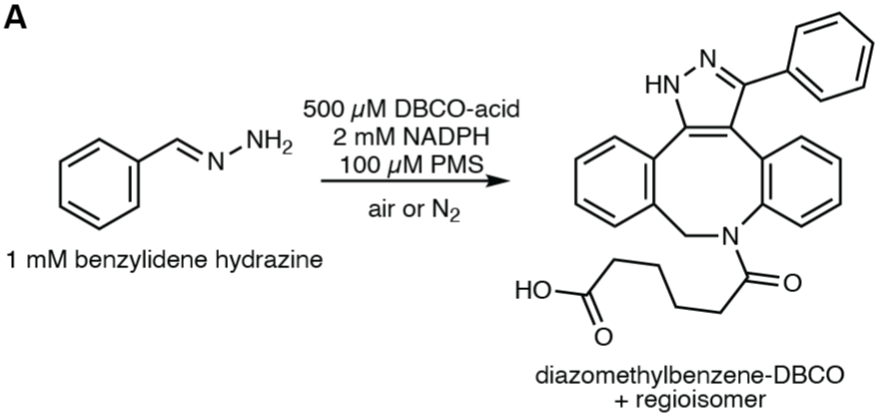


**B**

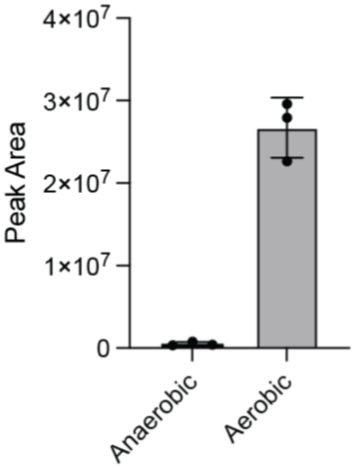


**Supplementary Figure 23:** Aerobic incubation of benzylidenehydrazine with DBCO-acid, PMS, and NADPH yields more product than anaeobic incubations. **A)** Reaction scheme. Reactions were incubated at pH 8.0. **B)** Combined LC–MS peak areas of both regioisomers of diazomethylbenzene-DBCO. Experiments were performed in triplicate. Error bars indicate mean ± standard deviation. Product identity was confirmed by comparison to a synthetic standard of diazomethylbenzene-DBCO.

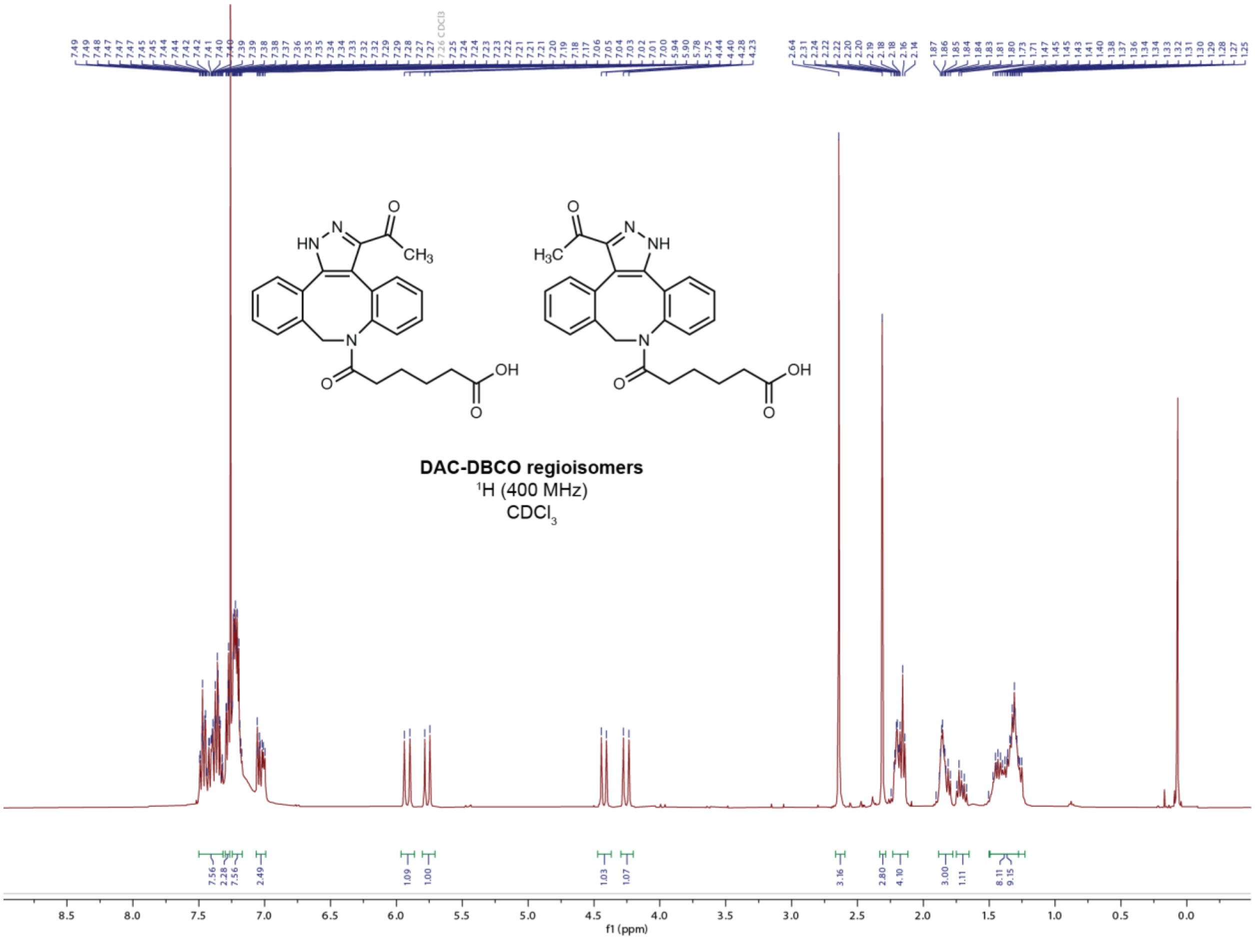


**Supplementary Figure 24:** $^{1}H$ NMR spectrum of DAC-DBCO regioisomers

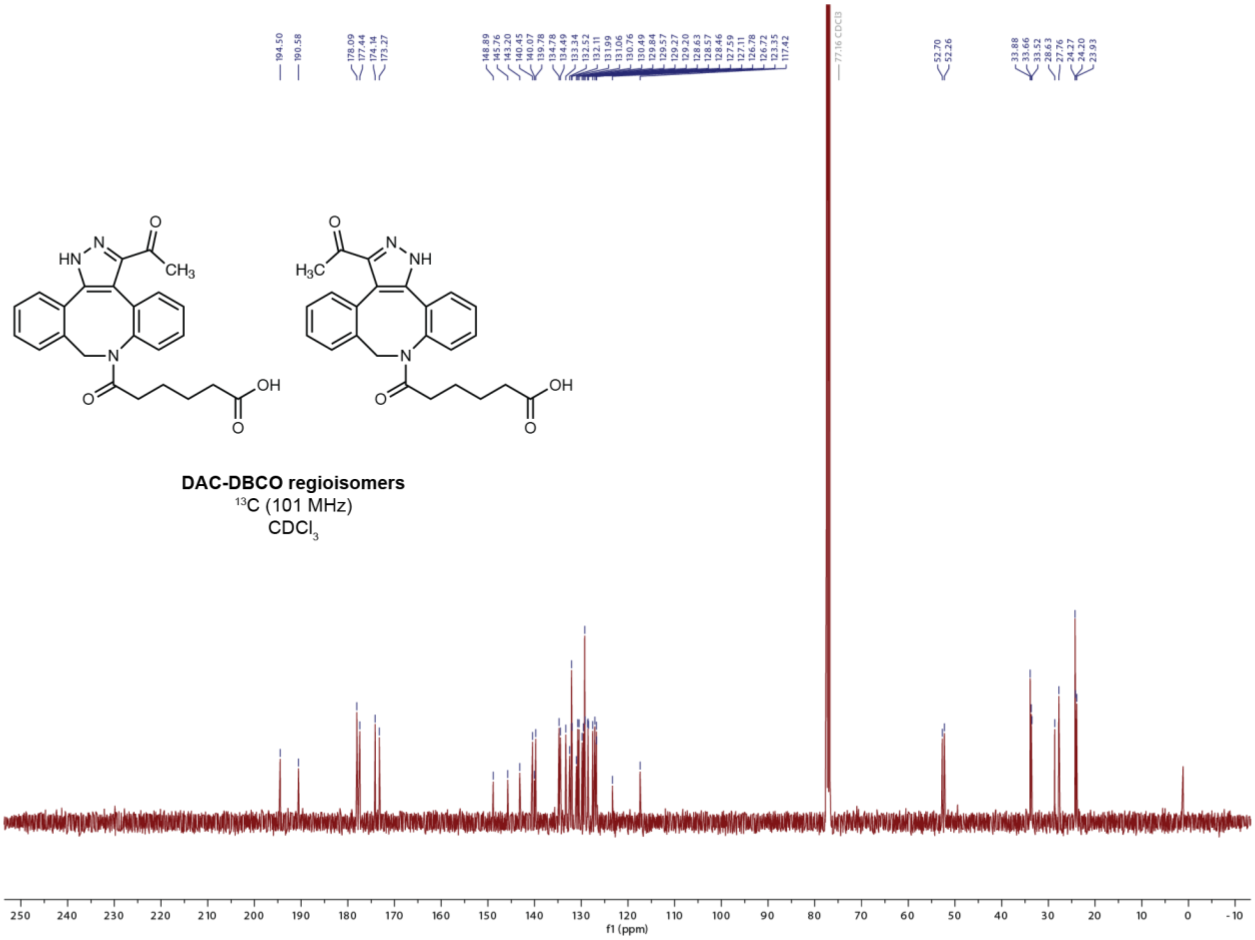


**Supplementary Figure 25:** $^{13}$C NMR spectrum of DAC-DBCO regioisomers

6
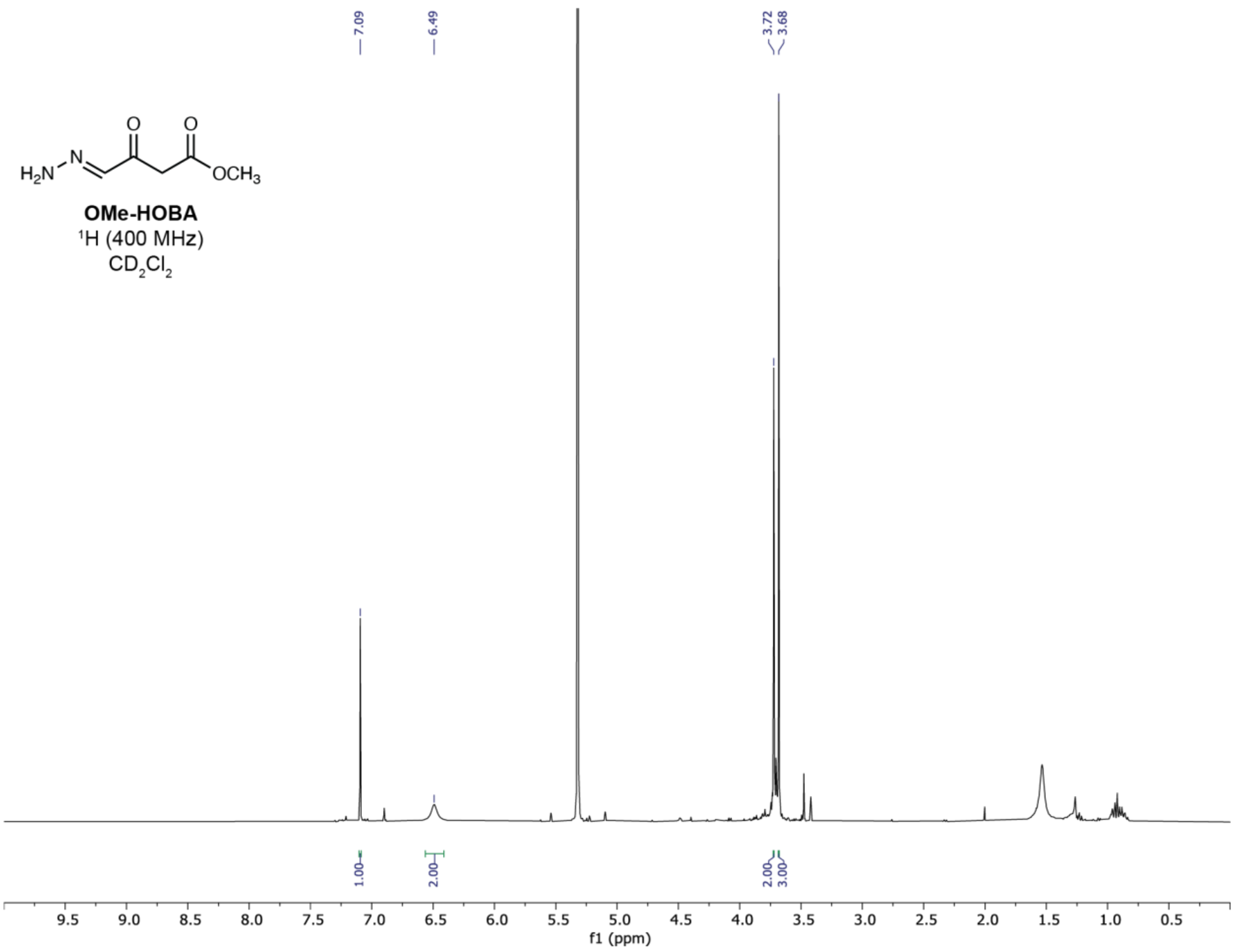


7 **Supplementary Figure 26:** $^{1}$H NMR spectrum of OMe-HOBA

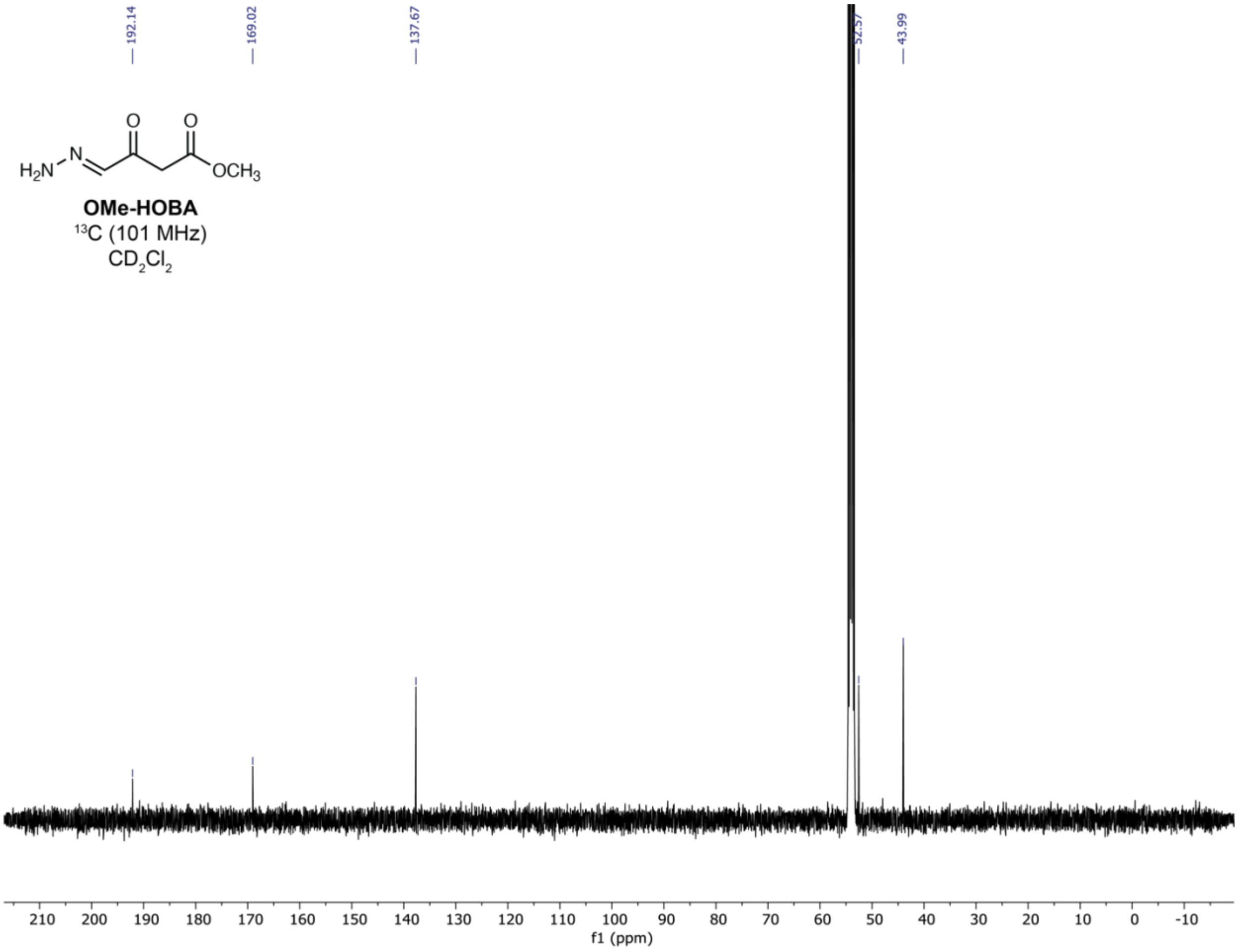


**Supplementary Figure 27:** $^{13}C$ NMR spectrum of OMe-HOBA

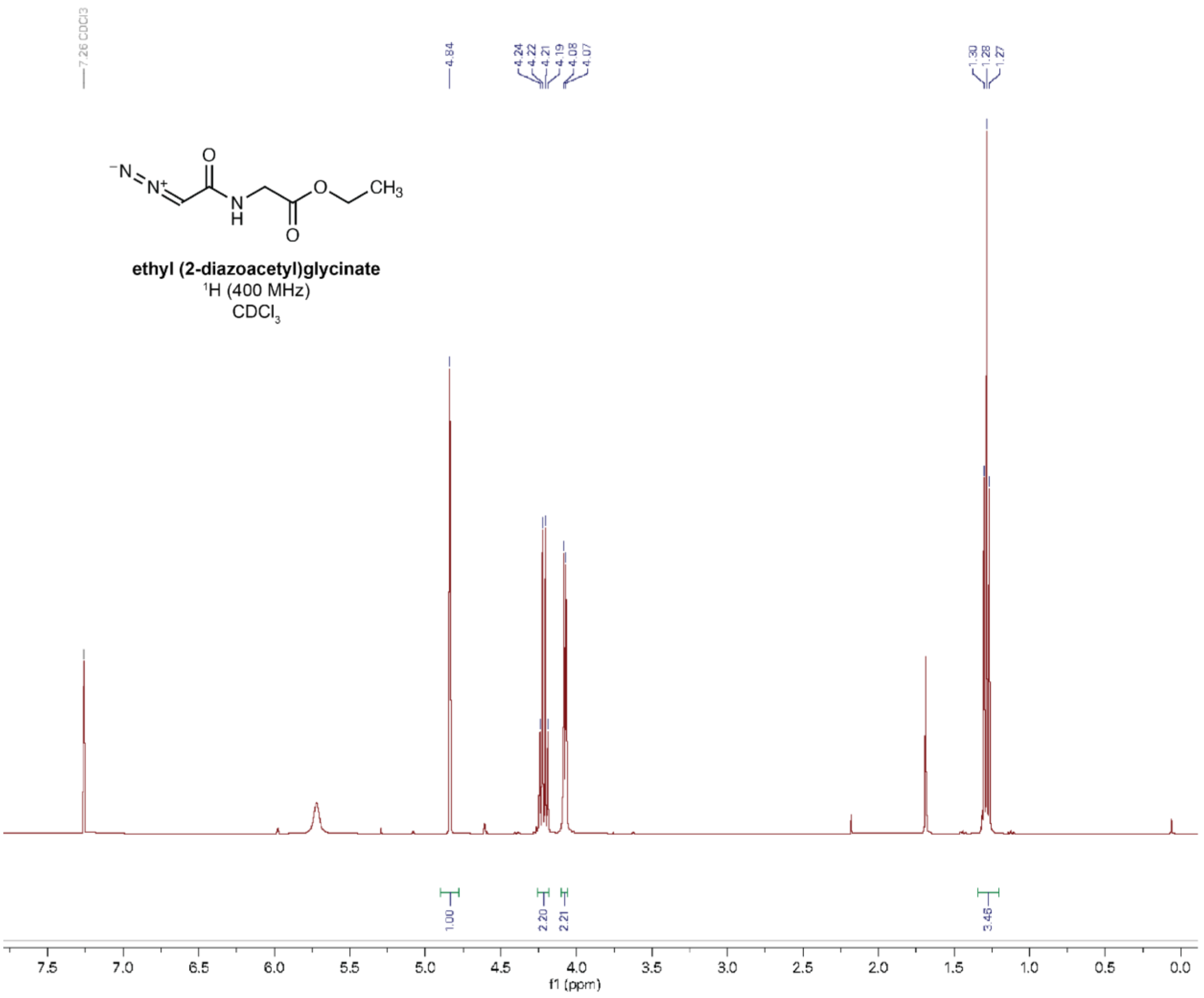


**Supplementary Figure 28:** $^{1}$H NMR spectrum of ethyl (2-diazoacetyl)glycinate

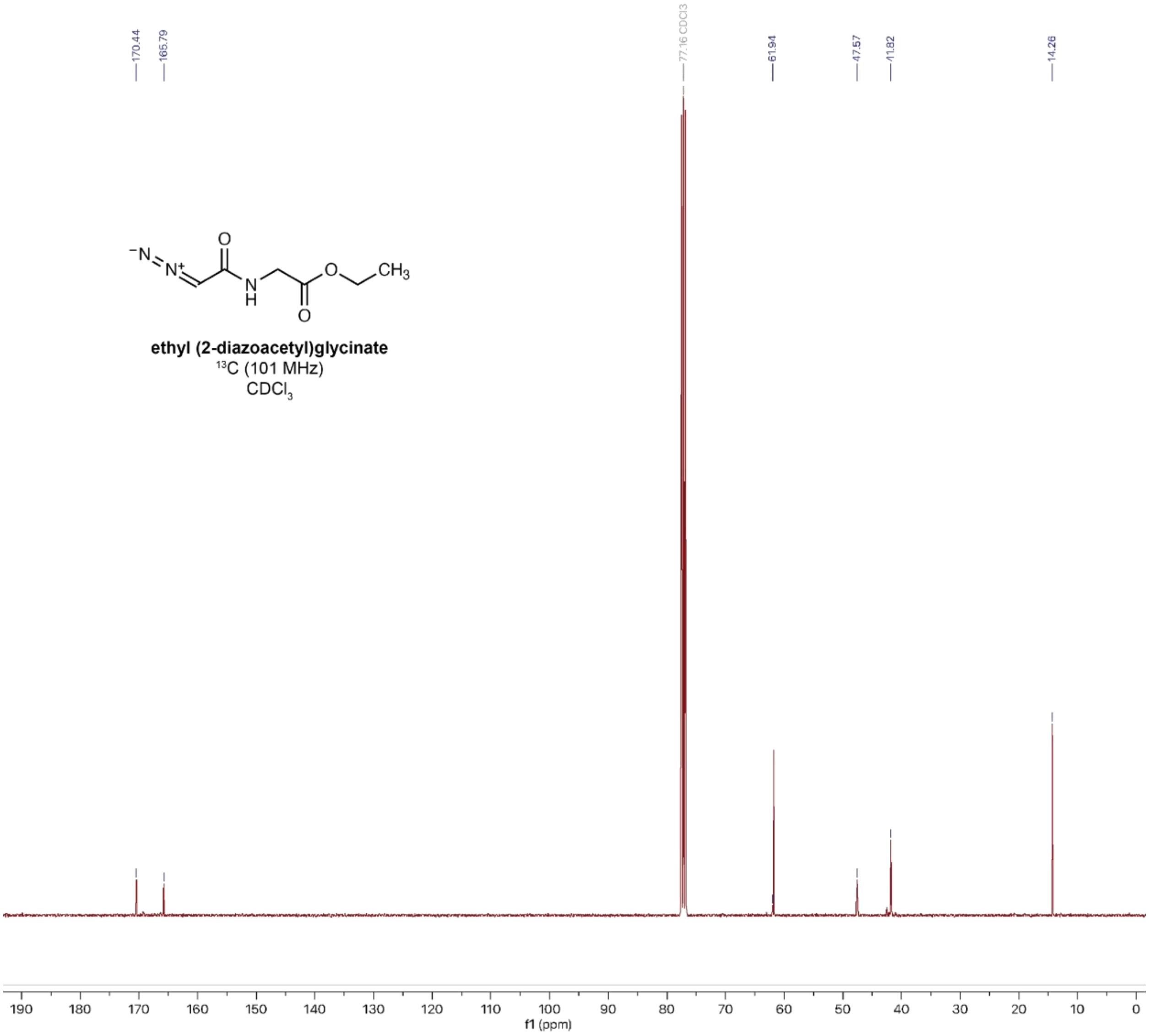


**Supplementary Figure 29:** $^{13}$C NMR spectrum of ethyl (2-diazoacetyl)glycinate

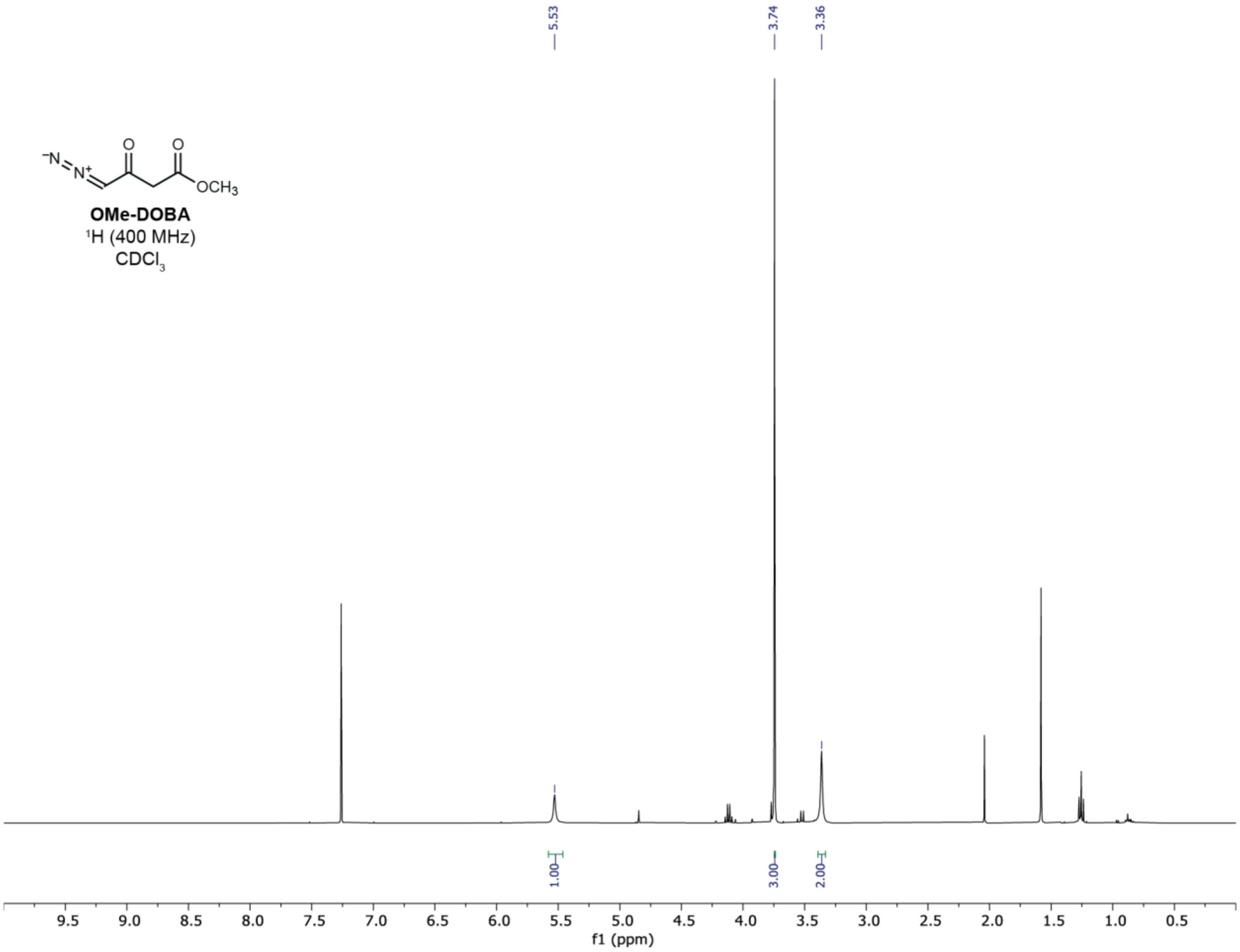


**Supplementary Figure 30:** $^{1}H$ NMR spectrum of OMe-DOBA

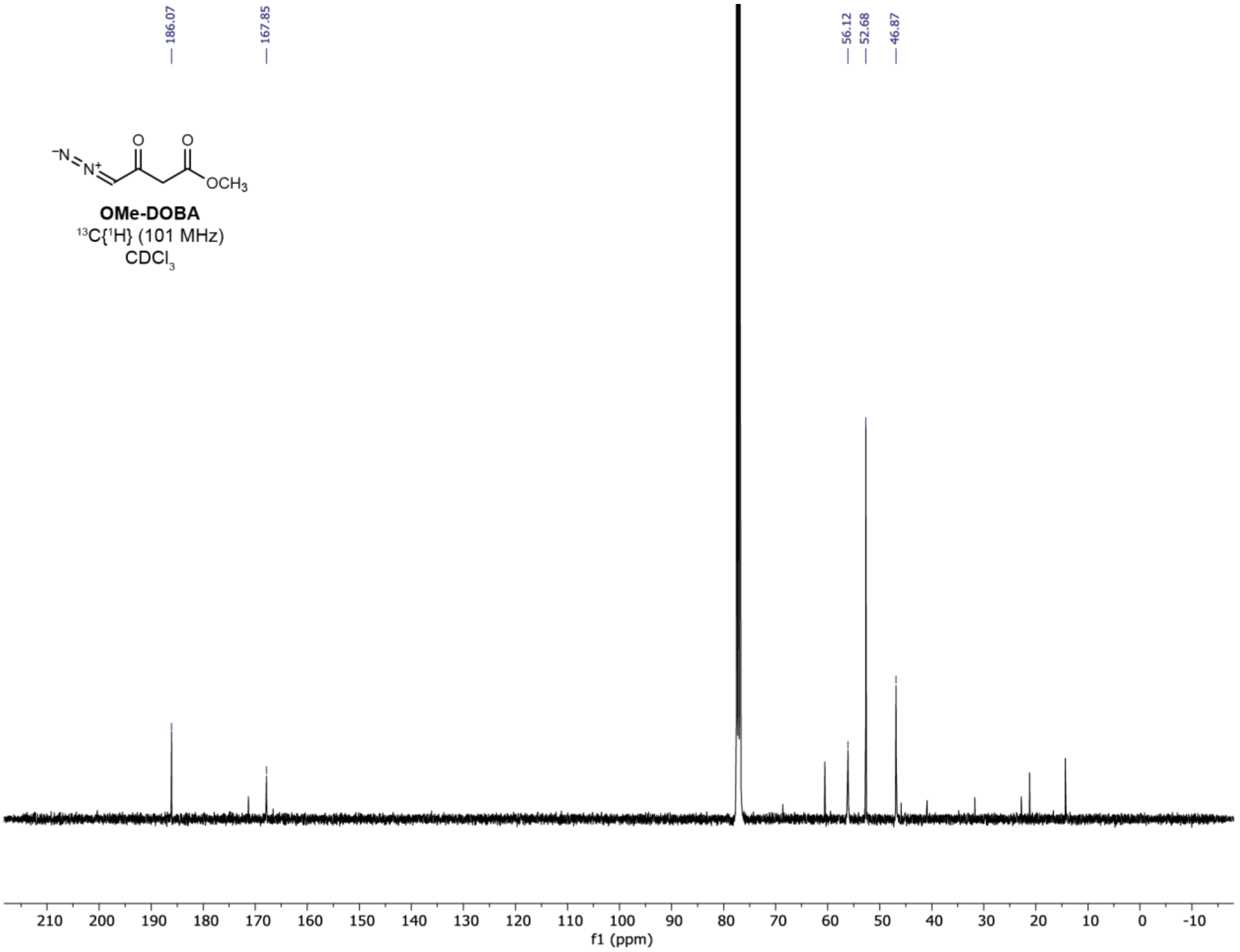


**Supplementary Figure 31:** $^{13}C$ NMR spectrum of OMe-DOBA

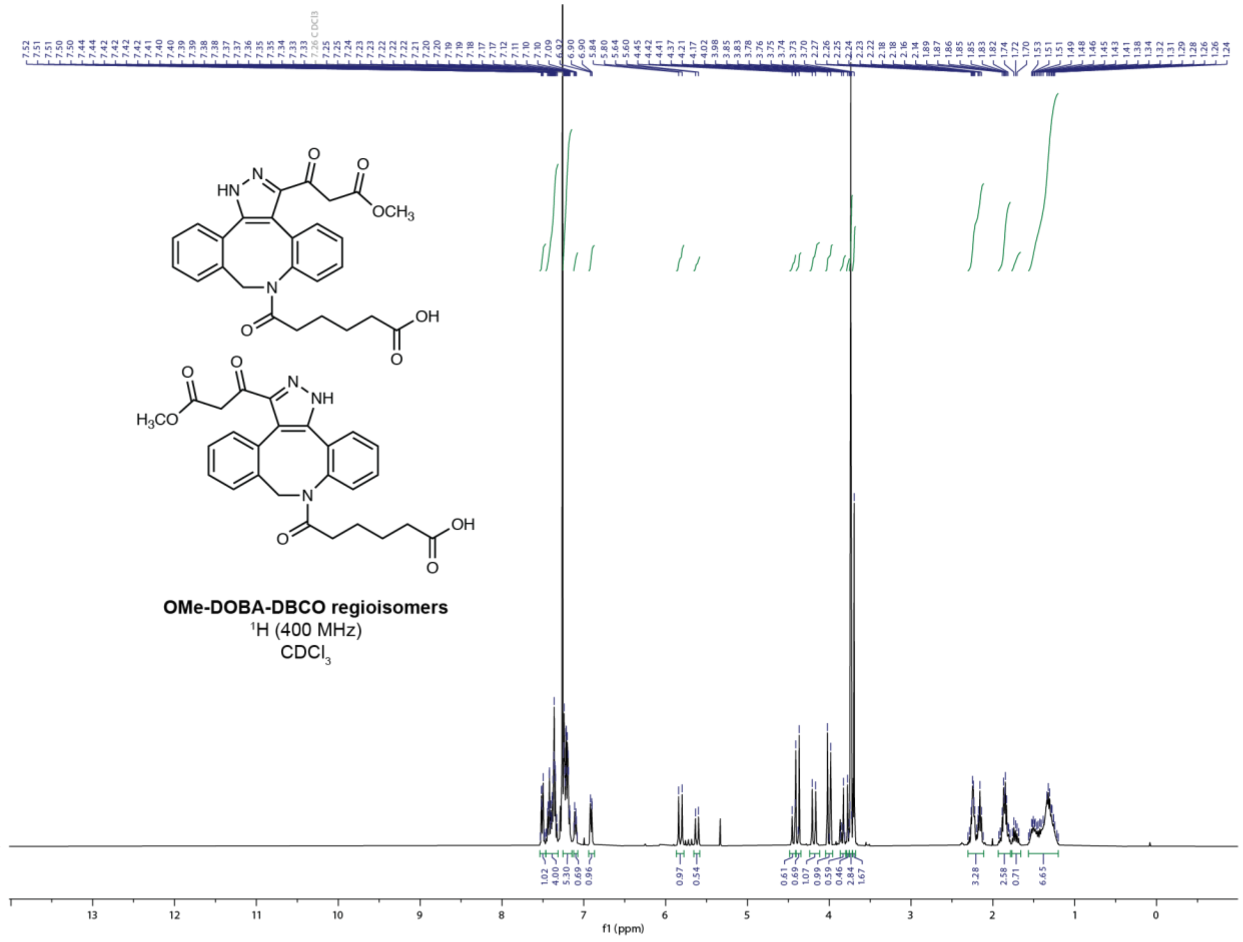


**Supplementary Figure 32:** $^1$H NMR spectrum of OMe-DOBA-DBCO

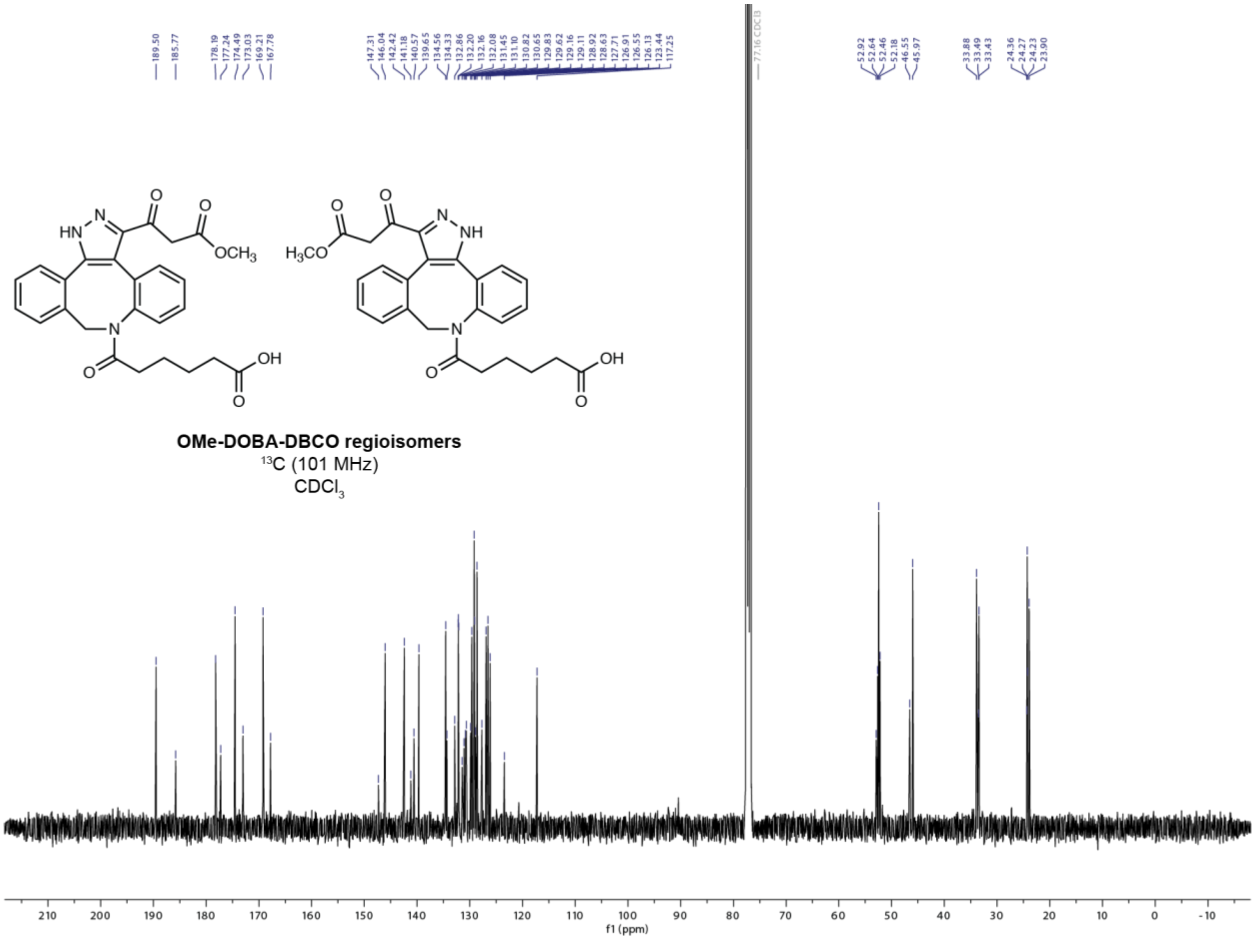


**Supplementary Figure 33:** $^{13}C$ NMR spectrum of OMe-DOBA-DBCO

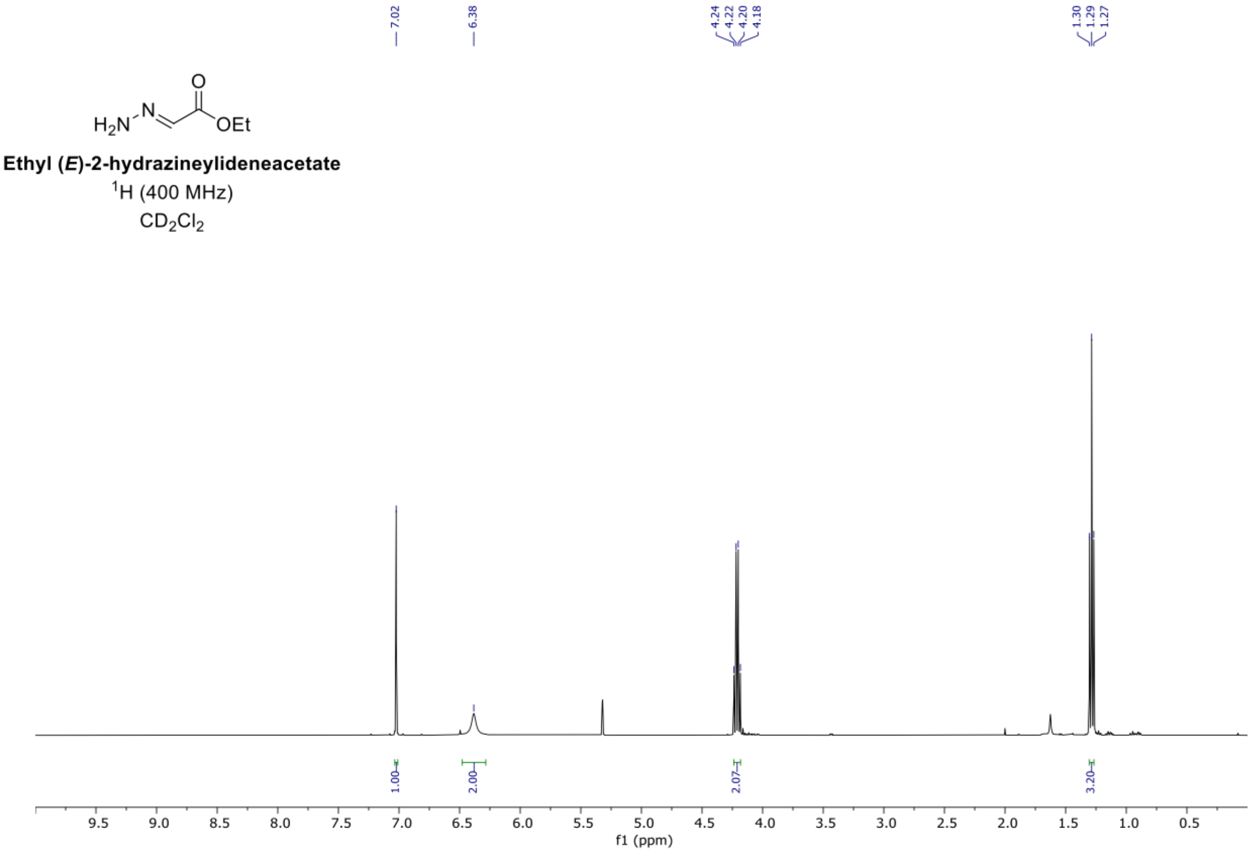


**Supplementary Figure 34:** $^{1}H$ NMR spectrum of ethyl (*E*)-2-hydrazineylideneacetate

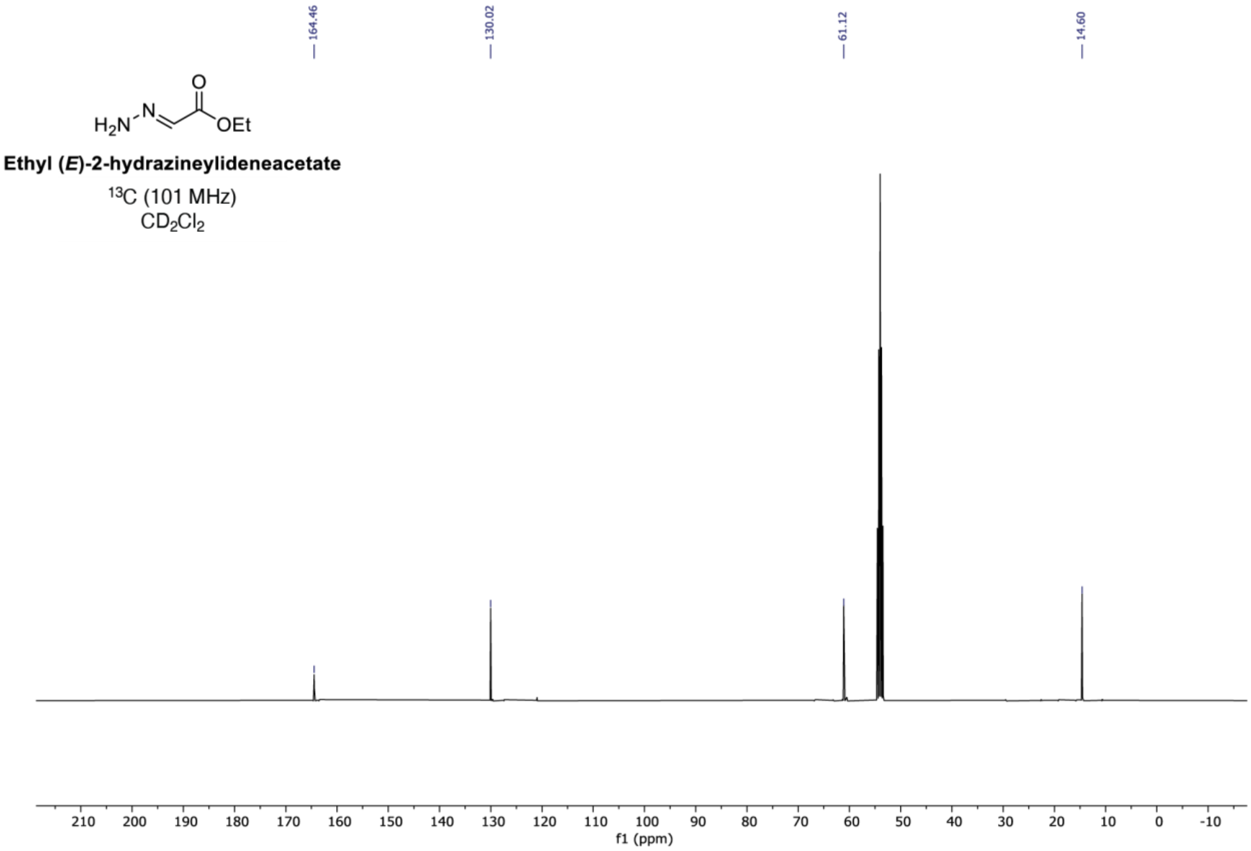


**Supplementary Figure 35:** $^{13}C$ NMR spectrum of ethyl (*E*)-2-hydrazineylideneacetate

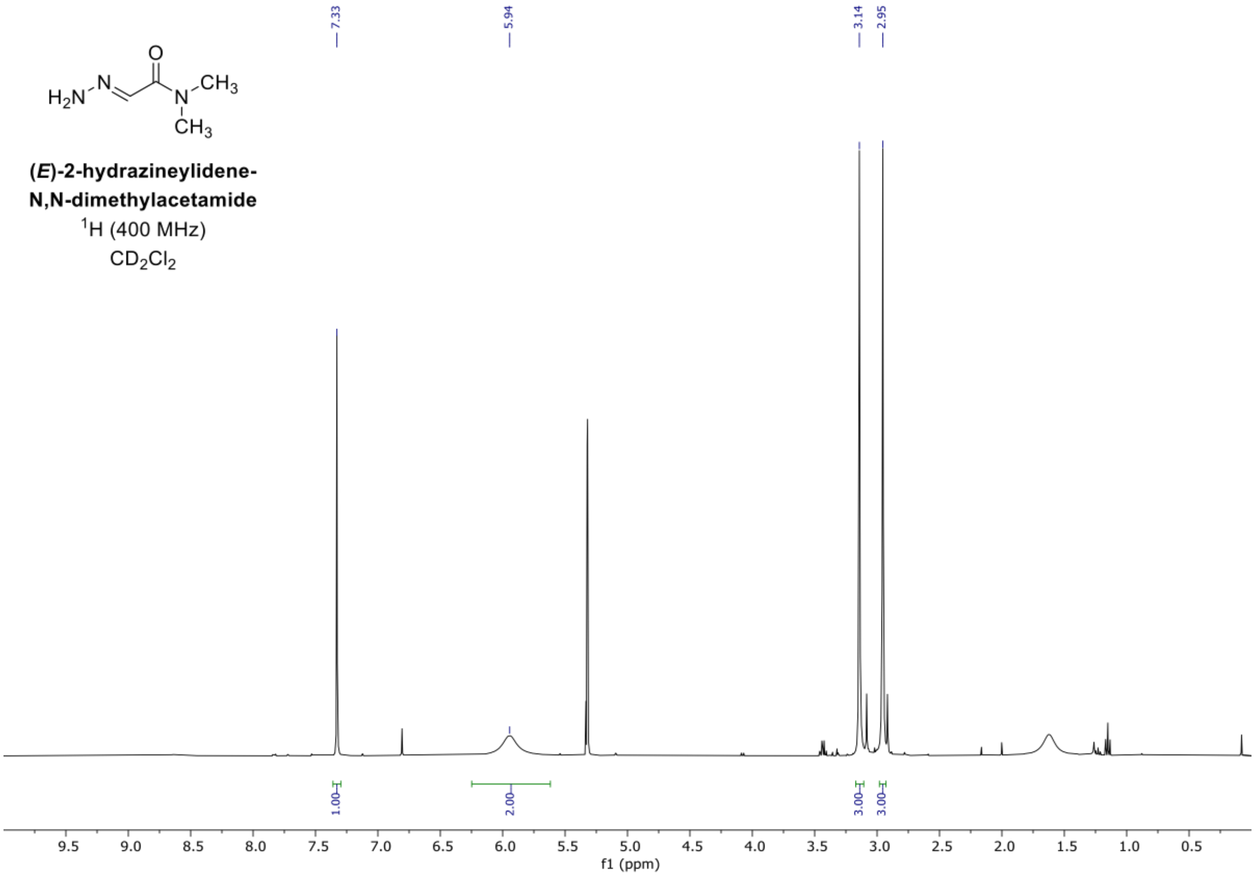


**Supplementary Figure 36:** $^{1}H$ NMR spectrum of (*E*)-2-hydrazineylidene-N,N-dimethylacetamide.

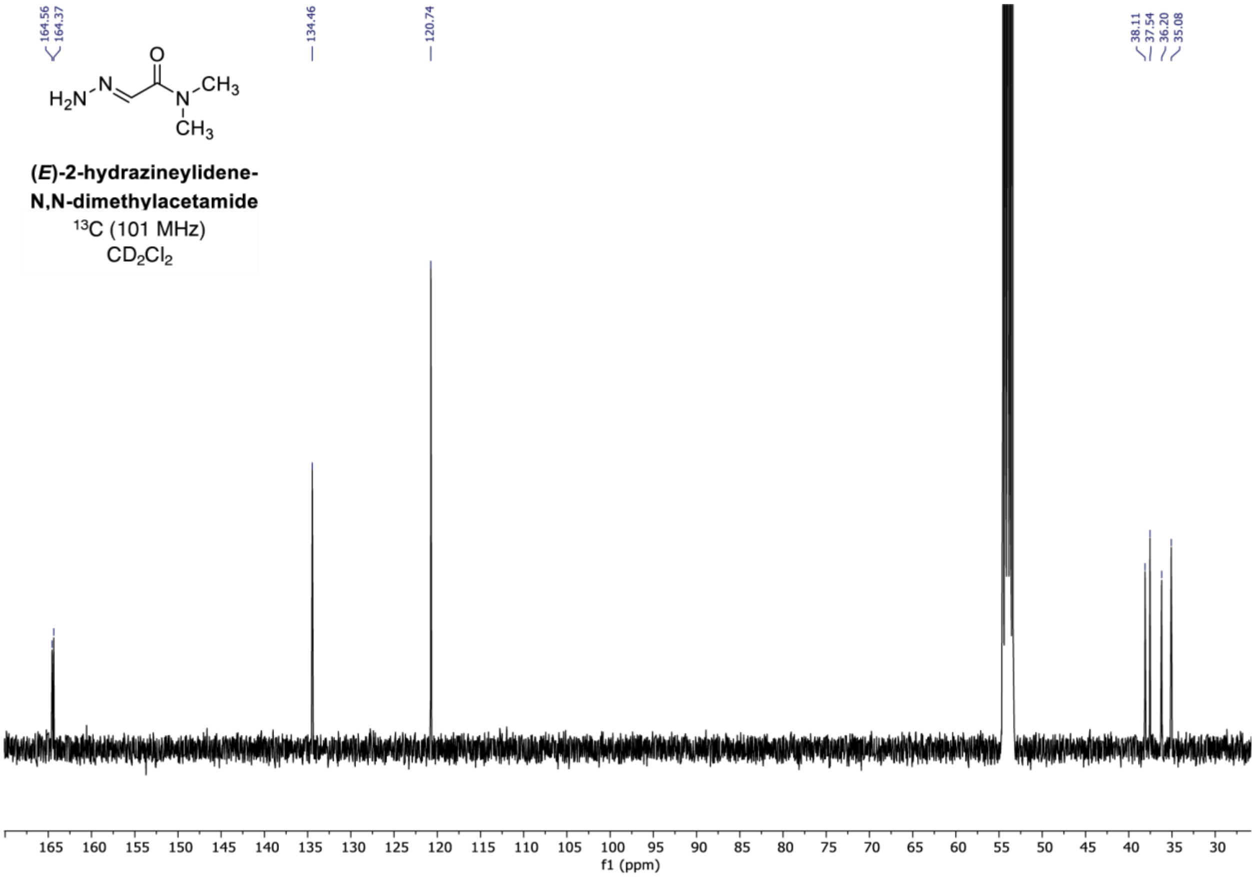


**Supplementary Figure 37:** $^{13}$C NMR spectrum of (*E*)-2-hydrazineylidene-N,N-dimethylacetamide.

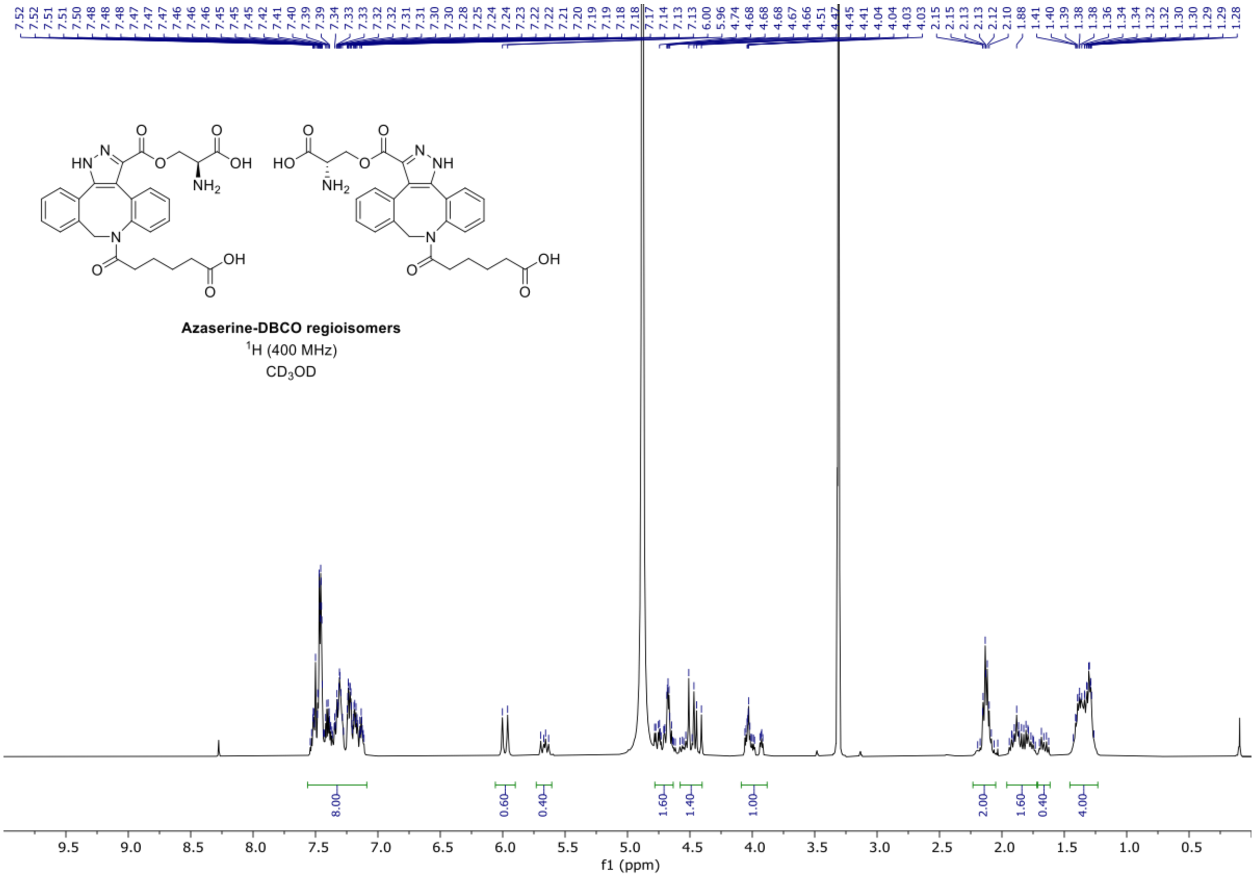


**Supplementary Figure 38:** $^1$H NMR spectrum of azaserine-DBCO

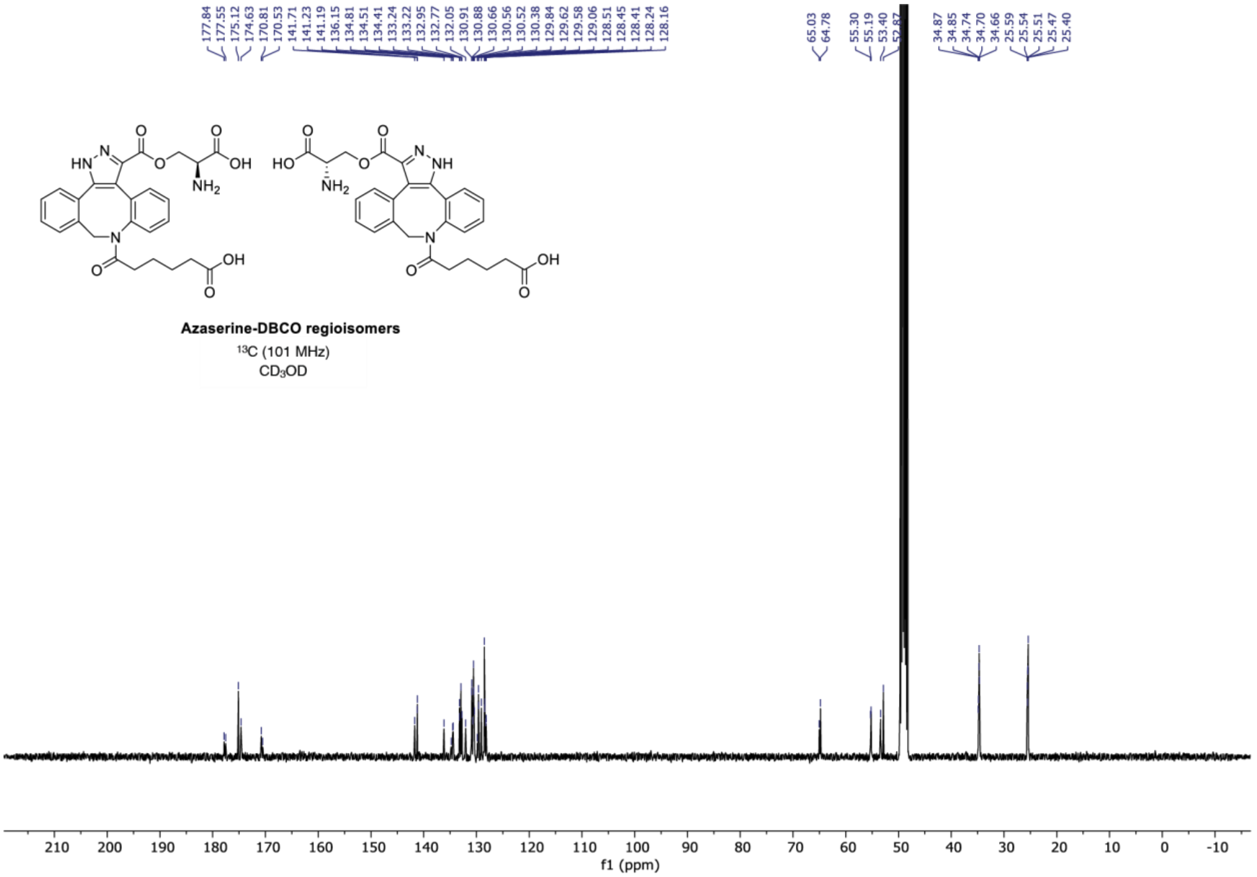


**Supplementary Figure 39:** $^{13}$C NMR spectrum of azaserine-DBCO

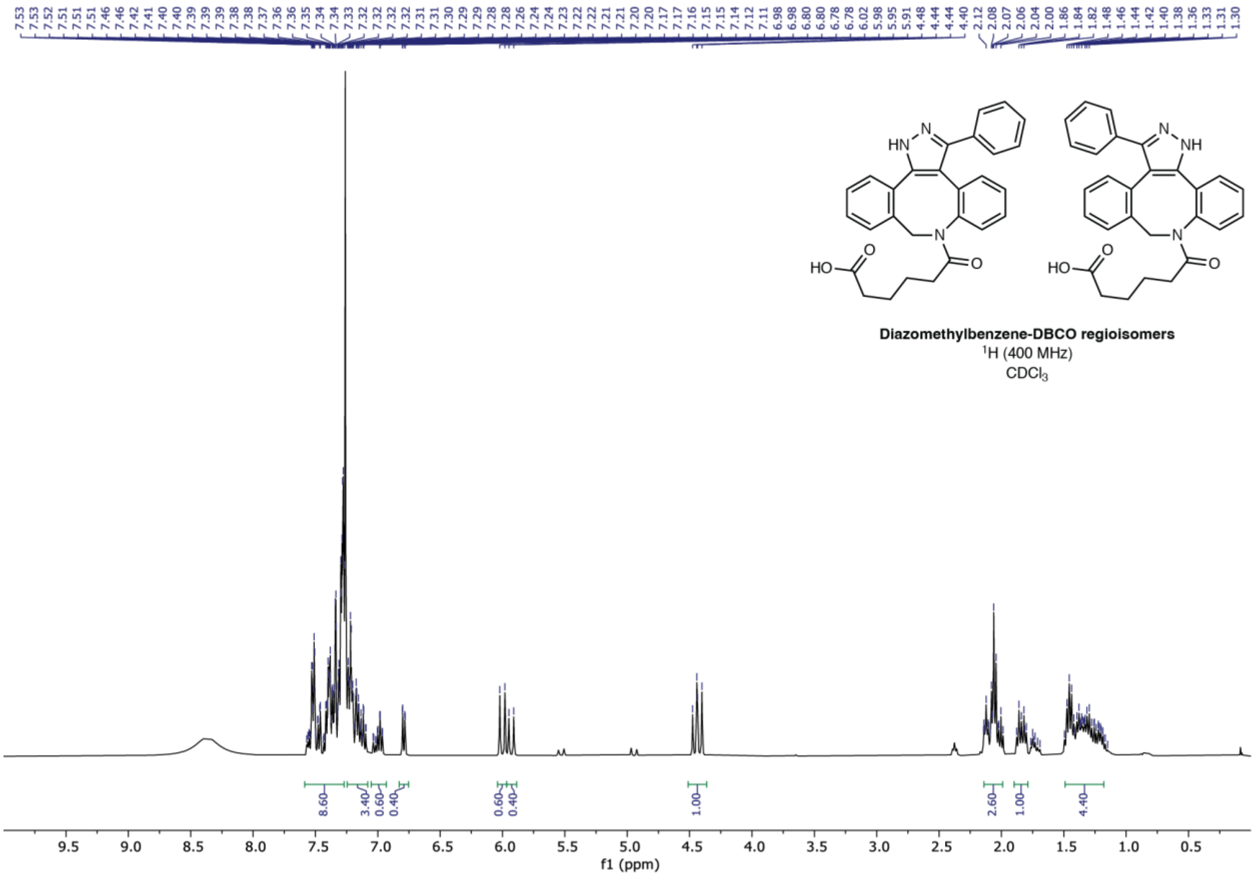


**Supplementary Figure 40:** $^{1}$H NMR spectrum of diazomethylbenzene-DBCO

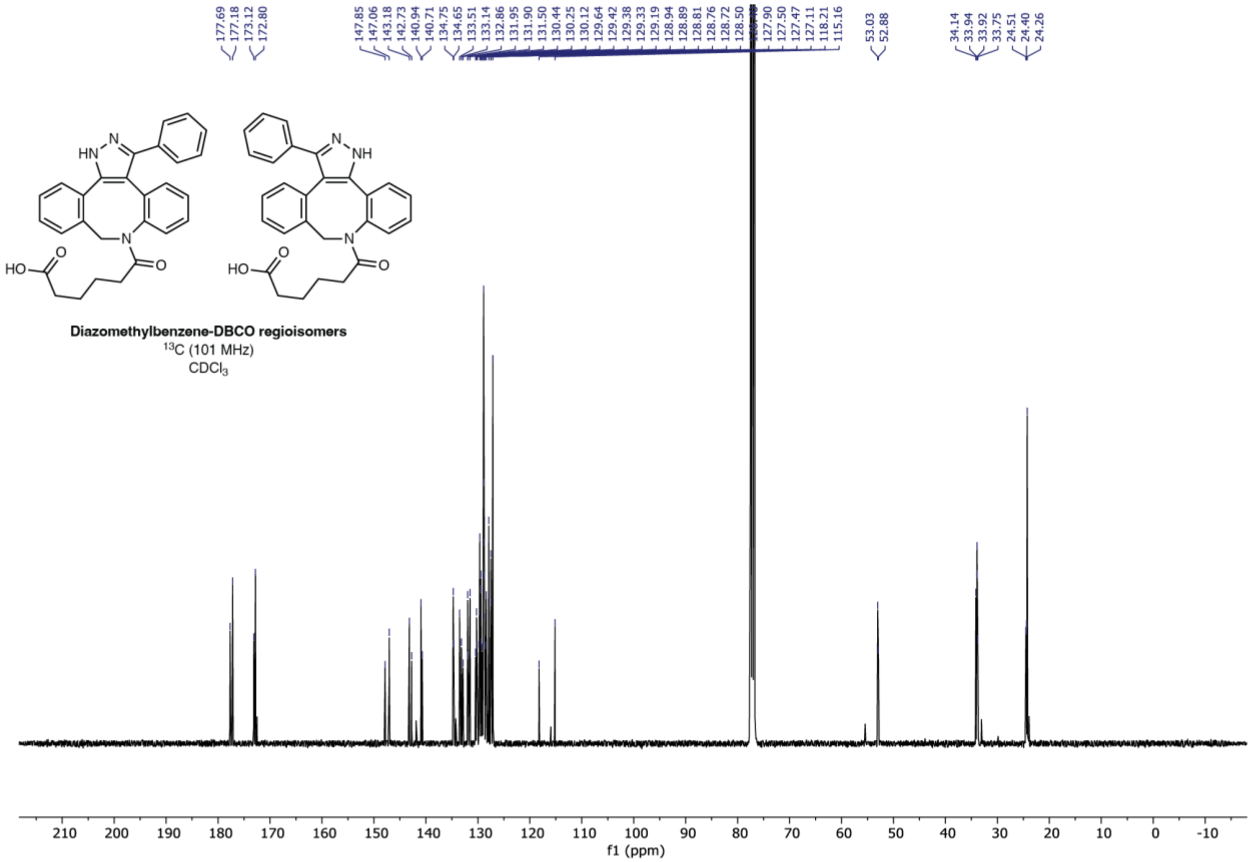


**Supplementary Figure 41:** $^{13}$C NMR spectrum of diazomethylbenzene-DBCO
